# Nanometer-Resolution Long-term Tracking of Single Cargos Reveals Dynein Motor Mechanisms

**DOI:** 10.1101/2022.01.05.475120

**Authors:** Chunte Sam Peng, Yunxiang Zhang, Qian Liu, G. Edward Marti, Yu-Wen Alvin Huang, Thomas C. Südhof, Bianxiao Cui, Steven Chu

## Abstract

Cytoplasmic dynein is essential for intracellular transport, but because of its complexity, we still do not fully understand how this 1.5 megadalton protein works. Here, we used novel optical probes that enable single-particle tracking (SPT) of individual cargos transported by dynein motors in live neurons over 900 *μm*. Analyses using the Fluctuation Theorem (FT) showed that the number of dynein molecules switches between 1-5 motors during the transport. Clearly resolved single-molecular steps revealed that the dwell times between individual steps were accurately described by an enzymatic cycle dominated by two equal and thermally-activated rate constants. Based on these data, we propose a new molecular model whereby each step requires the hydrolysis of 2 ATPs. The model is consistent with extensive structural, single-molecule and biochemical measurements.

## Introduction

Axonal transport of biomolecules and organelles is essential for neuronal function, especially since the length of the narrow axons prohibits transport by diffusion over distances from millimeters to meters. Proteins synthesized in the cell body are anterogradely transported by kinesin motors. Retrograde transport by dynein motors is used to recycle anterogradely transported proteins and to convey neurotrophic signals that report the condition of the distal axon terminals to the cell body (Fig. 1A). Robust bidirectional transport relies on coordination between multiple copies of the opposing motors, their regulatory mechanisms within the cell, and ultimately the fundamental biophysical properties of these motors. Dysregulation of axonal transport is thought to lead to neurodegenerative disorders (*1*).

**Figure 1.**
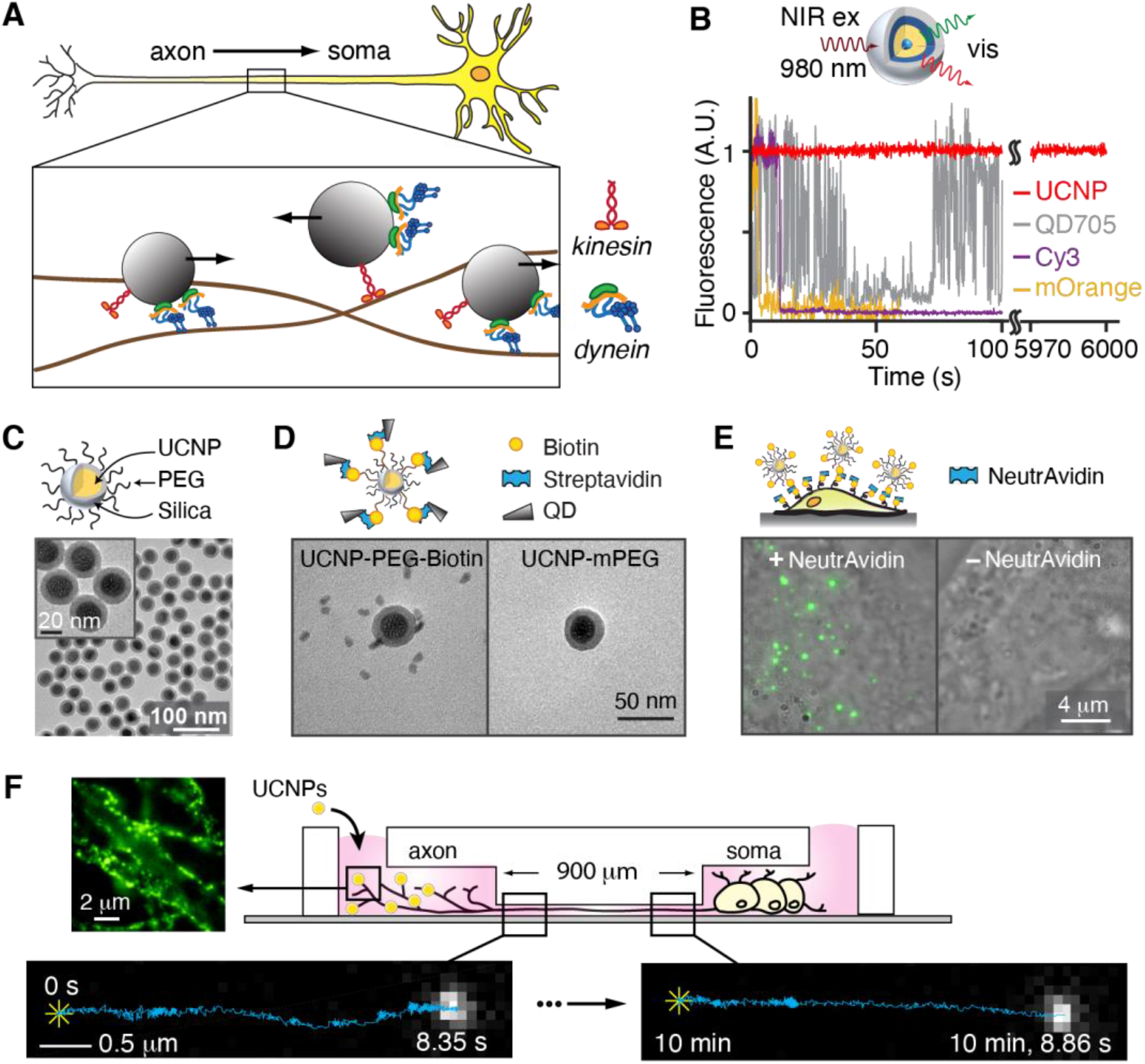
UCNPs used in single-particle imaging. (A) Schematic of axonal transport which involves endosomes being transported by kinesin and dynein motors. (B) (Top) Schematic design of the silica-coated core-shell-shell UCNPs used in live-neuron single-particle tracking. (Bottom) Fluorescence time traces of single UCNP (red), QD705 (grey), Cy3 dye (purple) and mOrange fluorescent protein (orange). (C) Transmission electron microscopy (TEM) images of the PEGylated NaYF_4_: 20% Yb^3+^, 2% Er^3+^@SiO_2_. (D) Satellite assembly of UCNPs and QDs via biotin-streptavidin binding. TEM images of mixture of streptavidin-conjugated QD705 and UCNPs bearing biotin (left) and without biotin (right). (E) Live-cell imaging of btn-UCNPs on HeLa cells whose membrane proteins have been biotinylated. Overlay of brightfield images of HeLa cells and maximum-intensity-projected luminescence images of biotinylated-UCNPs (btn-UCNPs) in the presence (left) and absence (right) of NeutrAvidin. (F) Schematic design of single cargo tracking during axonal transport in live neurons. Neurons are cultured in microfluidic devices with 900 μm long microchannels. Each field of view observes transport over 5 – 15 *μm*. Wheat-germ-agglutinin conjugated core-shell-shell UCNPs are incubated in the axon chamber to induce endocytosis. Representative luminescence image of DRG neuron axons labeled with UCNPs (left). (Bottom) Representative luminescence images of the same cargo during axonal transport from distal axon (t = 0 s, left) to near-soma (t ~10 min, right). Single-particle trajectories are overlaid on the fluorescent images where * indicates the start positions.

Despite the discovery of axonal dynein over 50 years ago (*2*) and of cytoplasmic dynein over 30 years ago (*3*), and despite the extensive and elegant structural, biochemical and single-molecule studies over the subsequent decades, the fundamental mechanism of dynein’s motor activity remain unresolved (*4*–*6*). Many pioneering *in vitro* experiments using purified motor proteins have offered significant insights (*7*–*11*), and much progress is being made in studies that include accessory proteins to these studies such as dynactin, adaptors, and dynein regulators such as Lis1-NudEL (*12*, *13*). Nevertheless, it is still desirable to perform live-cell neuronal studies of dynein dynamics to complement the reconstituted *in vitro* studies. However, the complex cellular environment and the high ATP concentration which allows the motors to move at speeds > 1 *μm/sec* complicates these experiments (*14*–*18*). A noninvasive imaging technique with high spatiotemporal resolution without photobleaching and blinking is essential to unambiguously monitor the rapid dynamics of neuronal cargoes over the entirety of their transport from the distal axon to soma.

Single-molecule fluorescence microscopy has been an essential tool in capturing the structure, interactions and dynamics of molecular machines such as dynein motors (*7*, *19*). However, the currently used fluorescent probes are not ideal. Organic fluorophores or fluorescent proteins exhibit fluorescence intermittency (blinking) and irreversibly photobleaching within tens of seconds (Fig. 1B). Numerous strategies have been developed to reduce fluorophore photobleaching and blinking, such as employing enzymatic oxygen-scavenging systems (*20*) or addition of triplet-state quenchers either in solution (*20*) or directly conjugated to the fluorophores (*21*). A combination of low concentrations (2%) of dissolved oxygen and a reducing-plus-oxidizing system was shown to enable single-particle tracking (SPT) in live cells for several minutes (*22*), but toxicity remains a concern for using these additives in live cells. Longer SPT was obtained by using a protein scaffold that recruits multiple fluorescent proteins or dyes (*23*–*25*), but these methods are still limited to the timescale of few minutes. Quantum dots (QDs) are more photostable, but their blinking prevents continuous imaging (Fig. 1B). Even when more benign materials are used, the photoreactive toxicity due to the generation of reactive oxygen species remains an issue (*26*).

Here we demonstrate the long-term single-particle tracking in live neurons enabled by photostable rare-earth ion doped ***u***p***c***onverting ***n***ano***p***articles (UCNPs). Their non-blinking luminescence is extremely stable, allowing single-particle imaging for many hours (Fig. 1B, Fig. S1). The near-infrared excitation into visible and UV emissions via multi-photon energy transfer among the sensitizer Yb^3+^ and emitters such as Er^3+^ and Tm^3+^ ions eliminate cellular autofluorescence, resulting in essentially background-free imaging (*27*–*33*). The sharp emissions can be tuned by varying both the type of emitter ions, their concentrations and ratios to perform multi-color imaging (*34*–*36*). Moreover, the use of near-IR excitation minimizes photodamage and permits deep-tissue imaging (*37*) and stimulation (*38*). Long-term tracking of UCNPs was demonstrated in HeLa cells and human lung carcinoma cells (*32*, *39*).

## Results and Discussion

### Functionalization of UCNPs

Ideally, nanoparticle functionalization should allow easy derivatization of various targeting molecules on the nanoparticle surface while maintaining low non-specific binding to cells (*40*). To this end, we used the reverse microemulsion method (*41*) to grow a 5-nm thin layer of silicon dioxide on 23 nm β-NaYF_4_ nanoparticles co-doped with 20% Yb^3+^ and 2% Er^3+^, resulting in monodispersed UCNPs in aqueous solution (Fig. 1C, Figs. S2, S3). The uniform silica coating allows direct conjugation of silane derivatives, such as silane-polyethylene glycol (PEG500), for mitigating nonspecific binding, and silane-PEG_3400_-biotin as the targeting group. Silica-coated UCNPs have been shown to exhibit no observable cytotoxicity when injected at high concentrations into mouse brain (*38*). Moreover, since UCNPs luminescence is due to inner-shell atomic-like transitions, the generation of reactive ion species such as singlet oxygen does not occur (*26*).

To validate the presence of the biotin moiety on the surface of the UCNPs, transmission electron microscopy (TEM) images were taken for a mixture of biotinylated UCNPs (btn-UCNPs) and streptavidin-coated QDs (Fig. 1D and Fig. S4). The btn-UCNPs were found to be decorated with QDs, forming a satellite assembly. In contrast, PEGylated UCNPs lacking biotin ligands did not bind QDs. An *in vitro* single-particle binding assay on PEGylated glass coverslip was used to quantitate the degree of nonspecific sticking of PEGylated UCNPs (Fig. S5). Specific binding of individual UCNPs via the biotin-NeutrAvidin interaction was evident, while control experiments without NeutrAvidin showed no binding. Fluorescence images of multiple fields of views (FOVs) established a lower boundary of 2000:1 specific versus nonspecific binding. To further assess the specificity of btn-UCNPs on live cells, we biotinylated membrane proteins on HeLa cells and performed similar binding experiments. Images of HeLa cells incubated with NeutrAvidin followed by btn-UCNPs showed bright fluorescent spots (Fig. 1E and Fig. S6), indicating successful targeting. In contrast, upconversion signals were absent in experiments without the addition of NeutrAvidin.

### Millimeter tracking of axonal transport

The photostable and background-free UCNPs are ideal for long-term monitoring of axonal transport in live neurons at high spatiotemporal resolution. To reduce the excitation power density for live cell imaging, we used our recently reported Yb^3+^-rich core-shell-shell (CSS) UCNPs, NaYF_4_@NaYbF_4_: 8% Er^3+^@NaYF_4_ (Fig. S7) (*30*). These 28 nm diameter particles are much brighter than the canonical core-only NaYF_4_: 20% Yb^3+^, 2% Er^3+^ due to the increased 980-nm light absorption by higher levels of Yb^3+^ ions, the improved crystal quality, and the protection against energy loss to the solvent by an inert NaYF_4_ shell.

For long-distance tracking of neuronal transport, DRG neurons were cultured in compartmentalized microfluidic devices so that their axons grew aligned within microchannels (*15*). This geometry allows simple differentiation between retro- and anterograde transport (Fig. 1F). The btn-UCNPs were conjugated to wheat germ agglutinin (WGA) which targets glycosylated membrane proteins and triggers receptor-mediated endocytosis (*16*). UCNP-containing endosomes that are transported by dynein motors towards the soma were imaged in the microchannels where there are no free UCNPs to interfere with the measurements. The same cargo was tracked through the entire length of the microchannel (900 μm) by recording several movies at 100 frames per second (f.p.s.) over a period of 10-15 minutes. Fig. 1F displays the unprocessed raw images. Since there is no cell auto-fluorescence, we were able to achieve the theoretical spatial localization precision 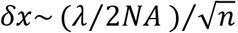, where *λ* is the wavelength of the detected light, NA is the numerical aperture of the microscope objective and *n* is number of detected photons.is achieved. For *n* = 1000,*δx* ~ 6.4 *nm*.

The representative trajectory (Fig. 2A) reveals a dynamic and complex behavior of a single cargo in the “stop-and-go” fashion, as previously reported (*15*, *16*). The non-blinking, photostable luminescence provides continuous records of retrograde transport that includes frequent motion reversals as well as cargo switching to neighboring microtubules (Fig. 2B). To quantitatively characterize these motions, a hidden Markov modeling (HMM)-Bayes approach was used to automatically annotate heterogenous motion states locally along a single trajectory (*42*). In contrast to the commonly used mean-square displacement (MSD) analysis, the HMM-Bayes method reveals stochastic switching between different motion states without the need to time-average over a significant number of particle displacement steps. Long-term SPT is ideally suited for HMM-Bayes analysis since the accuracy of the model prediction increases with the length of the trajectory.

**Figure 2.**
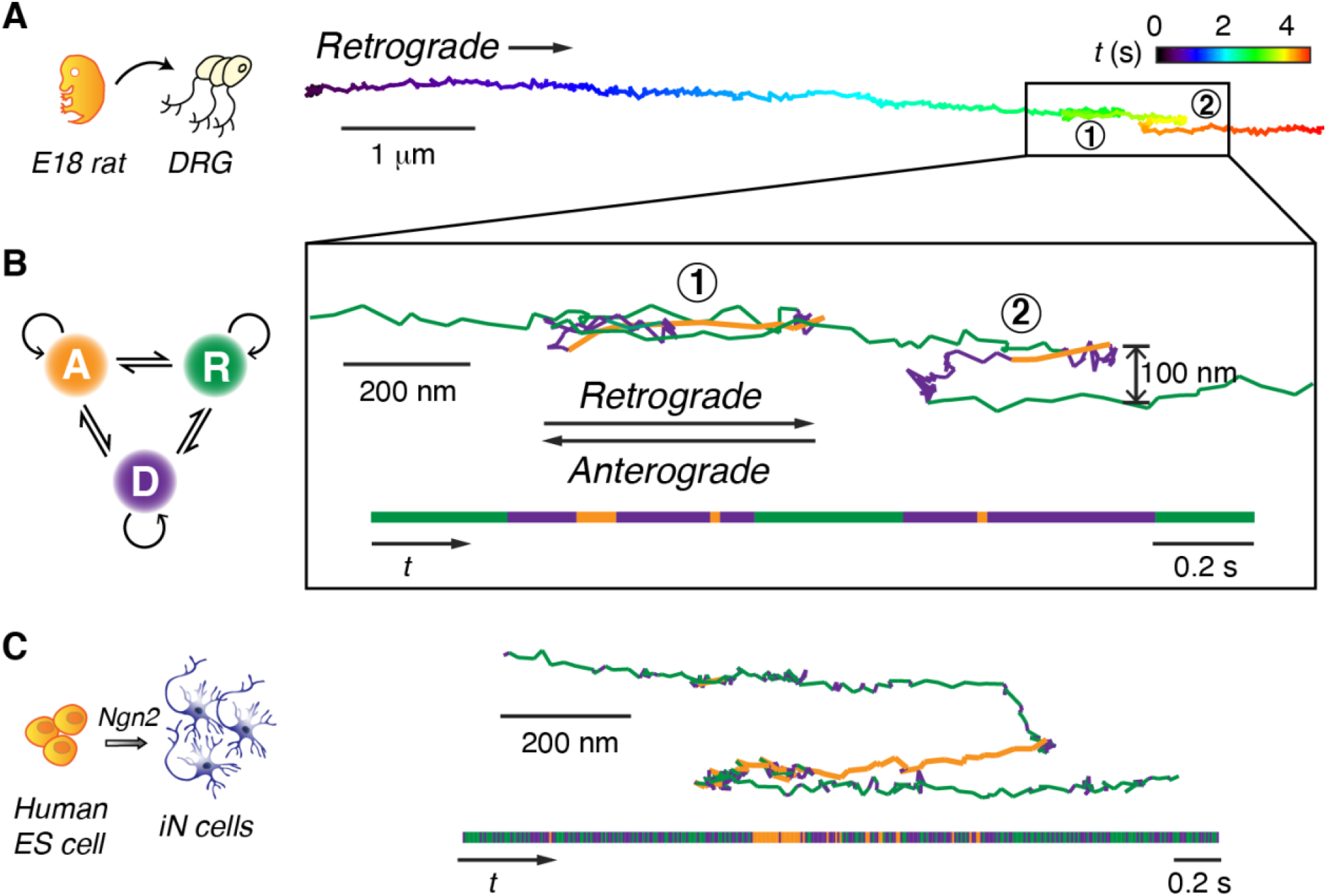
Long-term tracking of retrograde transport at high spatiotemporal resolution. (A) One example trajectory of retrograde cargo in rat DRG neurons that is color-coded in time, showing cargo reversal and track-switching (black box). (B) Hidden Markov model (HMM)-Bayes analysis of the single cargo trajectory in the boxed region in (A). The inferred motions include one diffusive state (blue) and two active transport states: retrograde (magenta) and anterograde (green) transport. The temporal sequence of motion states is illustrated in the horizontal bar on the bottom. The trajectory color-coded by motion states clearly shows reversal (Step 1) and track-switching (Step 2). (C) One example of single cargo tracking in human induced neurons (iNs) also illustrates track switching and rapid conversions between the diffusive, retrograde, and anterograde transport states. Equal scales used for horizontal and vertical dimensions.

HMM-Bayes analysis for the trajectory fragment boxed in Fig. 2A identifies one diffusive state along with two active transport states, which corresponded to retrograde and anterograde transport (Fig. 2B, Fig. S8). The retrograde transport had a lower average velocity of 2.81+0.18 μm/sec compared to that of the anterograde transport (5.06+0.02 μm/sec). Cargoes were often observed to come to a full stop and pause for a few seconds before continuing in the same direction (Fig. S8).

More complicated scenarios involving cargo pausing were also observed. Region 1 in Fig. 2B showcases an example where a short pause was followed by a rapid reversal on the same track. Such reversals should involve detachment of the opposing motors rather than the tug-of-war model (*16*, *43*). In region 2, the cargo was initially transported retrogradely. After pausing for 0.14 s, the cargo reversed direction and remarkably diffused to another microtubule that was ~100 nm away from the original one and resumed retrograde transport. While track changes may be explained by the presence of obstacles or cargo reaching microtubule termini (*43*–*45*), the regulatory mechanism for fast bidirectional transport in live cells remains elusive (*17*, *46*, *47*).

As a first step in future studies of axonal transport with human disease-associated mutations, we also studied single cargo transport in human induced neurons (iNs). Human embryonic stem cells were trans-differentiated into human excitatory neurons using transient expression of Ngn2 (Fig. 2C) (*48*). Similar to rat DRG neurons, these iNs also exhibit robust axonal transport with comparable behaviors including the presence of stop-and-go, reversal, and track switching motions (Fig. 2C, Fig. S8). Our results showed that there are common dynamic features of axonal transport rat and human neurons.

### The number of dynein motor pairs changes during retrograde transport

We now show that the distribution of cargo displacements can be used to determine the number of active dynein motors being used to transport a cargo. Previously, pulldown experiments and electron microscope images of neuronal endosomes indicated the association of multiple dyneins (*46*, *49*).Attempts to measure the number of active dynein motors include measurement of the stall force using large lipid droplets or ~ 1-μm beads (*14*, *18*, *50*), but the large vesicle size and the high laser intensity required introduced large perturbations to the cells. Nanoparticle-assisted optical tethering of endosomes suggested that axonal endosomes can recruit up to 7 dyneins (*51*), but the addition of a large optical trap force could have caused additional dynein motors to become engaged.

Here we apply the fluctuation theorem (FT) to measure the number of active dynein motors used during retrograde transport. The FT compares the positive entropy production of the system to the less probable negative entropy production for a system (*52*, *53*),

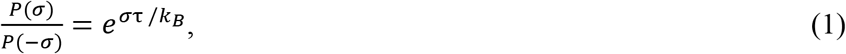

where *P*(*σ*) is the probability of observing an entropy production rate *σ* measured over a trajectory during a time *τ*, *P*(–*σ*) is the probability for the *time-reversed trajectory*, and *k_B_* is the Boltzmann constant. The FT is equally valid for a system in or out of thermal equilibrium with its surrounding environment. The FT has been recently used by Hayashi *et al*. in their measurement of the anterograde transport in ganglion neurons of individual dye-stained endosomes (*54*). However, our work differs significantly in how the FT is ultimately used to interpret the data. The FT has also been applied to measure the rotary torque of F_1_-ATPase (*55*).

We define our thermodynamic system as the cargo vesicle, dynein motors and associated cargo-attachment molecules. The change in entropy Δ*S* = *στ* in Equation 1 is measured experimentally as the translation of the cargo at position *x*(*t*) to a new position *x*(*t* + *τ*) = *x*(*t*) + Δ*x*(*τ*). The ability to track a ***single*** cargo continuously for extended periods of time allows the experimentally measurable probability distribution *P*(Δ*x*(*τ*)) to be determined precisely for individual cargos.

Time records of the displacement of one cargo in a DRG neuron at 37°C as it travels through the 900-μm microchannel are shown in Fig. 3A. For segments of roughly constant velocity (red curve in Fig. 3B), the normalized histograms *P*(Δ*x*(*τ*)) were calculated at different time delays *τ*. Fig. 3C plots *P*(Δ*x*(*τ*)) at *τ* = 10, 40, and 100 ms. The solid lines are fits to a Gaussian distribution 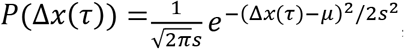, where *μ* and *s*^2^ = 〈*x*^2^〉 – *μ*^2^ are the mean and variance of the distribution.

**Figure 3.**
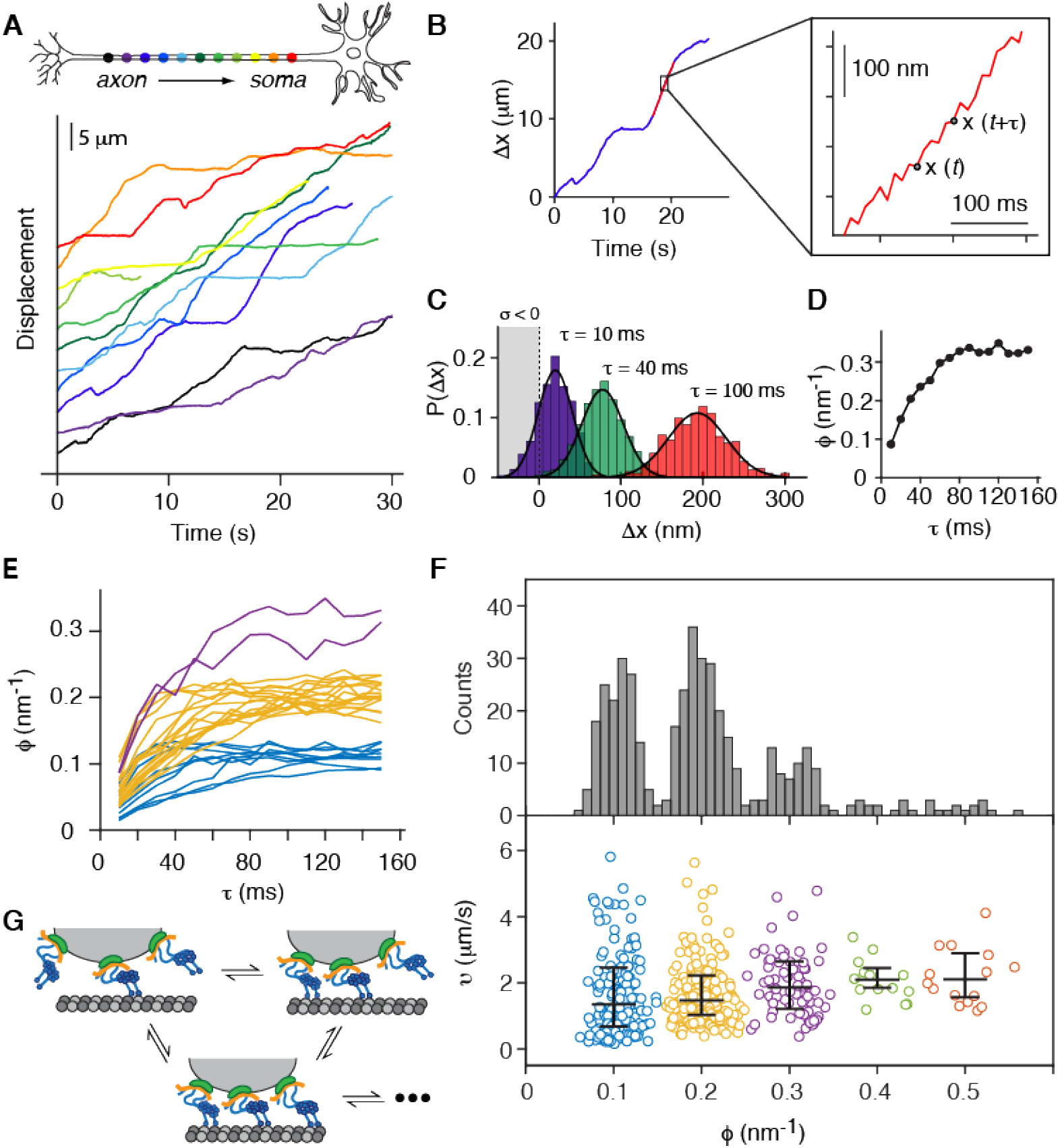
Fluctuation theorem analysis of retrograde transport. (A) Displacement curves of a single cargo being retrogradely transported in rat DRG neuron, from distal axon (dark purple) to soma (red) taken at 37°C. The curves obtained from separate movies were displaced vertically for better visualization. (B) Example of a constant velocity segment (red) used measure the set of displacements Δ*x*(*t*, *τ*) ≡*x*(*t* + *τ*) – *x*(*t*) for each starting time *t* and time delay *τ*. (C) The probability distribution *P_τ_*(Δ*x*) for a particular displacement labeled by the subscript *n*, Δ*x_τ_* (*t*, *τ*) = *x_τ_*(*t* + *τ*)–*x_τ_(t*) during different time delays *τ* = 10 ms (purple), 40 ms (green) and 100 ms (red). (D) Relaxation curve of the entropy change per unit length *ϕ* = 2*μ/s*^2^ in Equation 1. (E) All the relaxation curves of *ϕ* obtained from the displacement curves of the same cargo shown in (A). The curves are classified into three clusters using k-means and marked with different colors. (F) The histogram of 430 asymptotic *ϕ* values obtained from 12 single retrogradely transported cargoes, which shows clear asymptotic quantization of the entropy change per unit length. The corresponding velocities of the constant velocity segments are also plotted. The velocities are classified into 5 clusters using k-means clustering method. The central mark for each cluster indicates the median value, and the bottom and top lines indicate the 25^th^ and 75^th^ percentiles, respectively. (G) A cartoon model for our model where the number of independently engaged dynein motors stochastically changes during retrograde transport

The time-reversed displacement *P*(Δ*x*(–*τ*)) is found by making the substitutions Δ*x* → –Δ*x* and *μ* → –*μ*. Defining *ϕ* as the entropy change (normalized by *k_B_*) per unit length, Equation 1 becomes

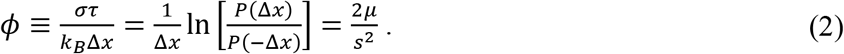

The value of *ϕ* (in units of nm’^1^) is seen to approach an asymptotic steady-state value, for times *τ* ≳ 120 *ms* in ~ 60% of the approximately constant velocity movements (Fig. 3D). The other 40% of the segments were rejected from further analysis. Fig. S9 is an example of a trajectory where *ϕ* does not reach a steady-state value, and Fig. S10 shows the entire trajectory of a single cargo. Remarkably, the resulting curves cluster into three discrete asymptotic distributions centered at *ϕ* = 0.1,0.2,0.3 nm^-1^ (Fig. 3E). This result indicates that the rate of entropy production per unit length is quantized. Since each dynein step requires hydrolysis of a certain amount of *ATP*, we interpret the three asymptotic values of *ϕ* as the increased entropy production due to the operation of 1 – 3 pairs of dynein motors (Fig. 3G).

The histogram for *ϕ* pooled from 12 different cargoes show well-defined and equally spaced peaks with the same quantized values of *ϕ*, even though it is expected that there is a distribution in the diameter of each vesicle (*46*). However, some cargoes exhibiting values of *ϕ* up to 0.5 *nm^-1^*, while other cargos exhibited only the two lowest values. (Fig. 3F and Figs. S11, S12). In sharp contrast, the corresponding velocity distribution exhibits large variations for the same asymptotic values of *ϕ* with only a modest increase in the average value with increasing *ϕ* (Fig. 3F). Similarly, the velocity distribution of the entire trajectory of a single cargo did not exhibit quantized peaks (Fig. S13). Our finding that the cargo velocity is not proportional to the number of motors is consistent with earlier *in vitro* experiments (*8*).

We interpret the steady-state quantization of *ϕ* as the number of active, independently tethered motor complexes. While it is clear that the entropy production per unit displacement would be proportional to the number of operating motors, it is less intuitive that increased entropy production would lead to a more “orderly” displacement, as indicated by *ϕ* = 2*μ/s*^2^. Note that we differ from the FT analysis of Hayashi *et al*. (*54*) who interpreted the quantization of *ϕ* as the quantization of their “force producing units” on each endosome.

We present a quantitative justification for this interpretation. If we assume that the displacement after many steps of *each* motor system *i* = 1,2 is described by Gaussian probability distributions 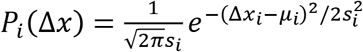. If the two motor systems attached to the same vesicle are assumed to operate independently, the combined probability distribution *P*(*x*) = *P*_1_(*x*)*P*_2_(*x*) can be shown to also be a Gaussian distribution. If the two motors are identical (*μ*_1_ = *μ*_2_ ≡ *μ*, 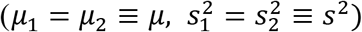), the mean displacement and dispersion of a vesicle driven by the two motors is *μ* and *s*^2^/2. For *N* equal independent motors, the mean displacement is unaffected, but the dispersion is reduced *s^2^→ s^2^/N* (*56*–*58*). Using the definition of *ϕ* (Equation 2), *ϕ*_*N* motors_ = 2*Nμ*/*s*^2^ = *Nϕ*_single motor_, which justifies that the equally spaced quantization of *ϕ* is due to the number of active motors.

The FT analysis of retrograde transport in human neurons found similar *ϕ* distributions (Figs. S14, S15), as expected from the high degree of structural conservation of cytoplasmic dynein among mammals. The deduced number of active dynein motors associated with both rat and human neuronal endosomes (average of 2.01 ±0.05 and 2.50 ±0.08 for rat and human neurons, respectively) are in good agreement with previous EM work and western blotting results (*46*, *49*).

### Measurement of single dynein steps in live neurons

Live cell transport measurements were made at 22°C, 30 °C and 37°C (Fig. 4). In order to time-resolve individual molecular steps of dynein in physiological conditions where the average velocity can be as high as 6 *μm* · *s*^-1^ (Fig. 3F), we synthesized brighter optical probes which allowed 1 *ms* time resolution. Larger UCNPs used to record the step-resolved trajectories were 66 *nm* (Fig. S16) for experiments at 22°C and 30 °C, and 160 × 90 nm (Fig. S17) for experiments at 37°C. With the largest UCNPs, ~5 × 10^6^ photons per second detected.

**Figure 4.**
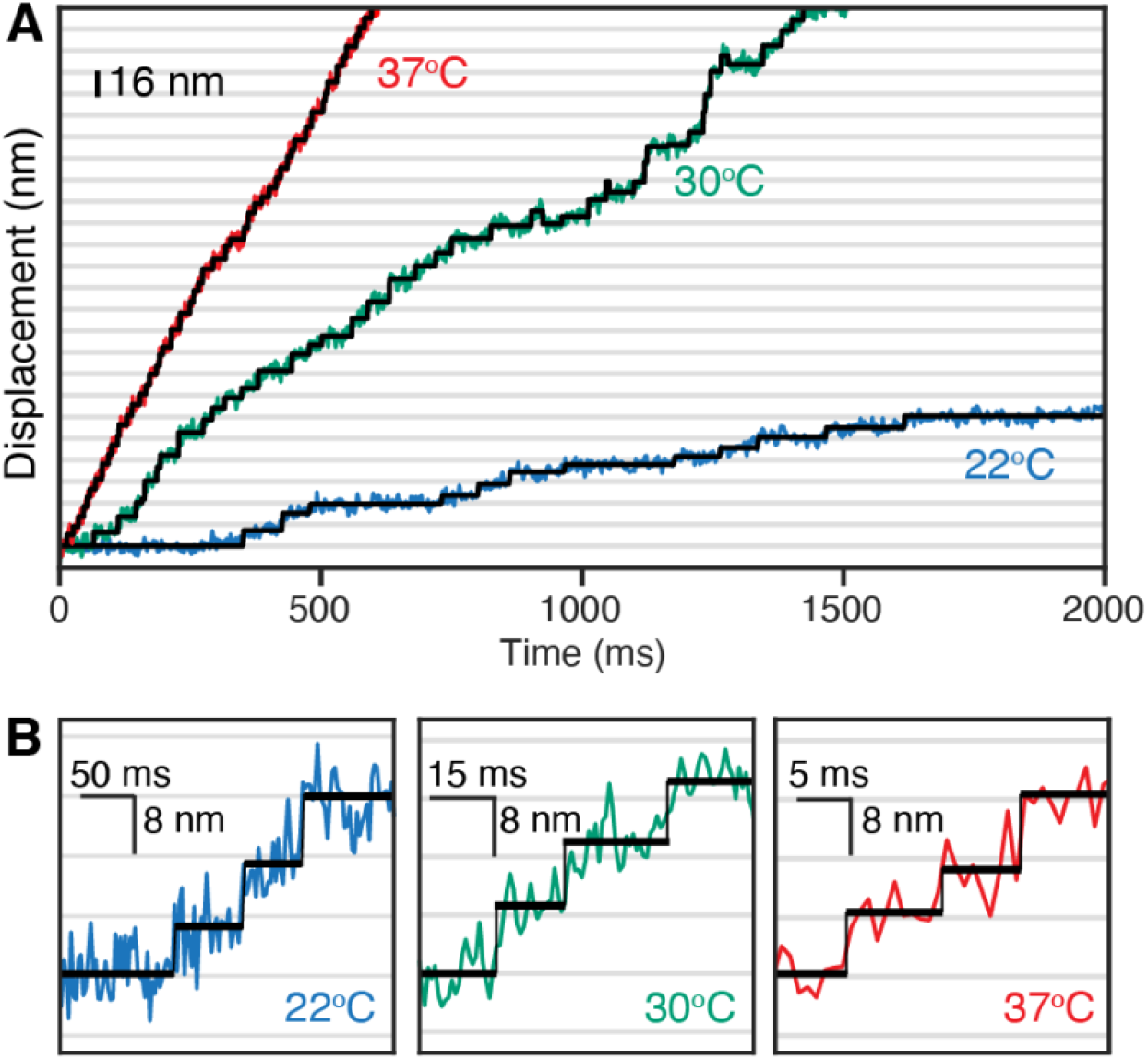
Dynein steps in live DRG neurons. (A) Stepping traces of retrogradely transported endosome labeled with larger probes at three different temperatures (red: 37 °C, green: 30 °C, blue: 22 C. We synthesized and used 80 × 60 nm core-shell-shell UCNPs (NaYbF_4_: 10% Gd, 8% Er@NaYbF_4_: 8% Er@NaYF_4_, Fig. S16) for experiments at 22° C and 30 °C. Larger 160 × 90 nm UCNPs (Fig. S17) were synthesized for experiments at 37 °C. The video frame rate of 1 msec consisted of a 0.8 ms integration time and 0.2 ms read-out time. (B) Expanded views of resolved single-step time traces taken at for 22°C, 30°C and 37°C. Individual motor steps were identified using a step-finding algorithm (black line).

The step-size histograms (Fig. 5A) display multimodal distribution with partially resolved peaks at ~ 8.3 nm and larger steps show that the step-size variability is temperature-dependent (Fig. S18 shows more example traces). At 22 °*C*, the histogram of live cell transport which involves full-length dynein, dynactin, cargo adaptor, vesicle, and dynein regulators is consistent with room temperature *in vitro* measurements of purified *tail-labeled* yeast dynein, but narrower than the distribution for head-labeled dynein (*58*, *59*). Previous *in vivo* step size measurements of dynein includes reports of a narrow distribution at 8 nm (*60*) while others observed broader distribution (*14*, *61*). As the temperature increases to 37 °C, the dynein velocity increased by ~7x (based on the decrease of the dwell-time between steps) and the step-size histogram became significantly narrower.

**Figure 5.**
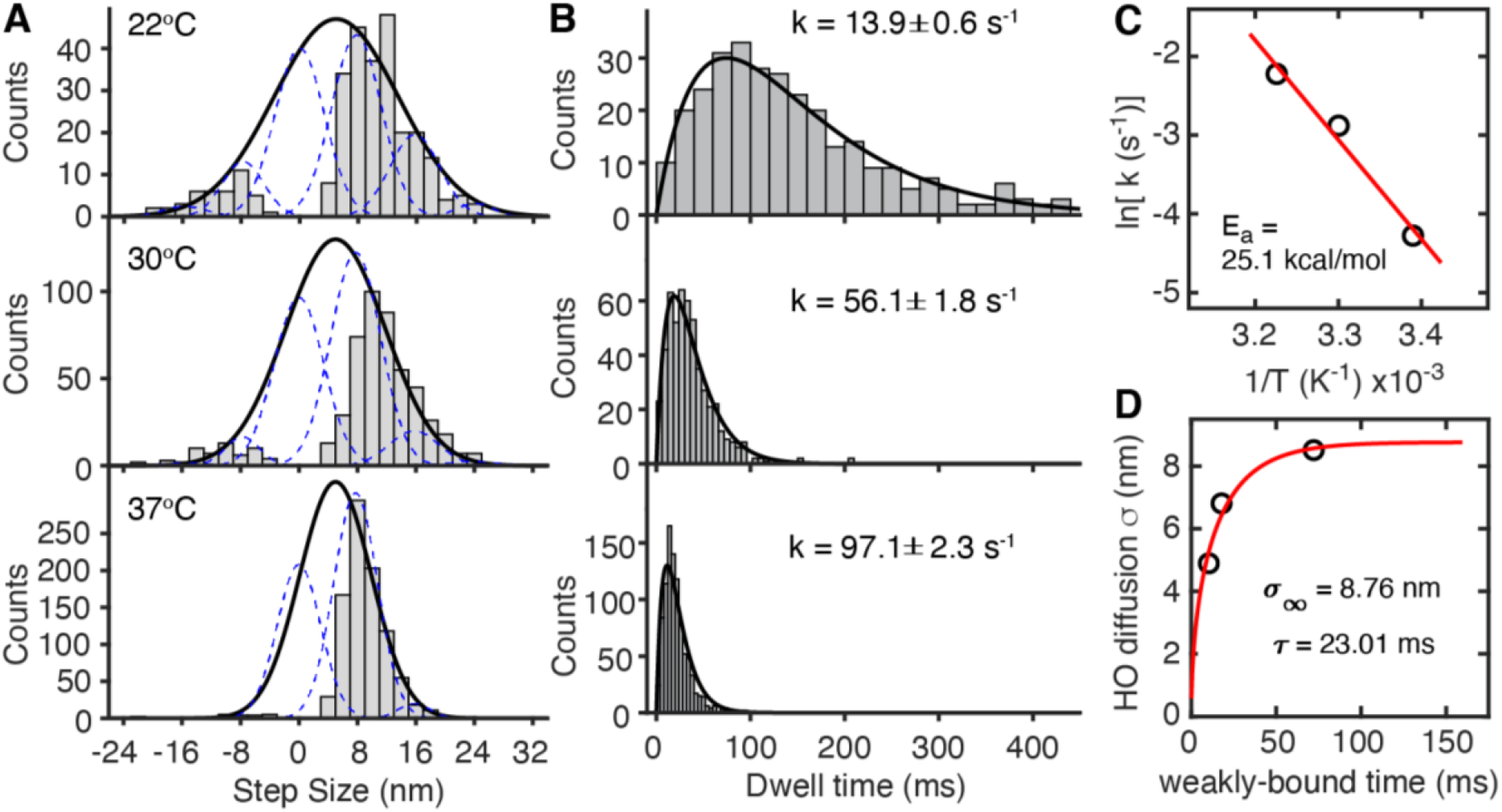
Dwell-time histograms and fit to two equal, thermally activated rate constants. (A) The step-size histogram at 22 °C displayed multimodal distribution with a major peak at 8.3 nm and larger (12-24 nm) The step-size histogram obtained from a single cargo showing variable step sizes with a major peak at 8 nm, where the width of Gaussian fits (solid black lines) is temperature dependent and indicative of a constrained random walk diffusion between power strokes. The step positions given by the blue dotted lines are displaced by increments of ~ 8 nm. The number of recorded steps in the histograms are N = 283, 278 and 920 steps for 22°C, 30°C and 37°C, respectively. The MTBD can only bind at certain locations on the MT due to the periodicity of tubulin subunits and the interactions between MTBD and MT, and the +5 *nm* distance represents the average probability of the cargo moving a step forward, backward or remaining in the same position after a full enzymatic cycle. (B) The temperature-dependent histograms of dwell time between steps. The solid lines are a convolution of two exponential functions with the same rate constants (*τk*^2^*e*^−*kτ*^, where *k*(22 °C) = 13.9 ± 0.6 *s*^-1^; *k*(30 °C) = 56.1 ± 1.8 *s*^-1^; *k*(37 °C)) = 97.1 ± 2.3 *s*^-1^). To be insensitive to the choice of the histogram bin size, the rate constants were determined by fitting the raw dwell times using Maximum Likelihood Estimates (MLE) as discussed in Figs. S19. The average time of a single step is calculated from the probability distribution 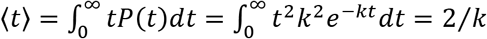 for each temperature. (C) Arrhenius plot of the equal rate constants *k* ≡ *k*_1_ = *k*_2_ at 22 °C, 30 °C, and 37 °C. Activation energy of 25.1 kcal/mol was obtained. (D) The temperature dependent histogram widths *σ*(*T*) fit (red line) to a model where the diffusive search of the stepping dynein monomer is constrained by a linear spring and when the MTBD is weakly bound to the MT.

The dwell times shown Figs. 4B and 5B decreases significantly with temperature to an average time ~20.6 msec at 37°C. The dwell-time histograms at all three temperatures fit well to a dynein cycle described by two sequential reactions that are formalized as two equal rate constants *k*_1_ = *k*_2_ ≡ *k* where the time *τ* to take the next step is given by *P*(*τ*) = *τk*^2^ exp(–*kτ*). The presence of two equal and thermally-activated rate (*~e^−E_a_/k_B_T^*, where *E_a_* is an activation energy, Fig. 5C) leads us to propose a new dynein stepping model in which two *ATP* molecules are hydrolyzed per dynein step. See Fig. S19 for discussion of the method used to fit *P*(*τ*) and uncertainties assigned to the fit. The *in vitro* single-molecule measurements show different dwell-time distributions when dynein was labeled at the head or tail (*58*); since the dynein monomers are joined at their tails, tail-labeled dynein may be less perturbative.

### A new model for dynein stepping

Before presenting the proposed model, we summarize some of previous insightful studies of dynein. The dynein motor consists of two > 4000 amino acid heavy chains (HC), each containing a motor domain built around a ring of six AAA+ (ATPases Associated with various Activities) and attached to a long tail (*47*), as indicated schematically in Fig. 6. The binding of *ATP* between the AAA1 and AAA2 domains (which will be referred to as the “AAA1 site”) has been identified as the primary catalytic site (*6*). The ATPase activity at the interface between AAA3 and AAA4 (the “AAA3 site”) has also been determined to be essential for dynein motility (*62*), whereas the cycling of *ATP* on the other AAA sites is not believed to be required (*5*, *6*, *63*).

**Figure 6.**
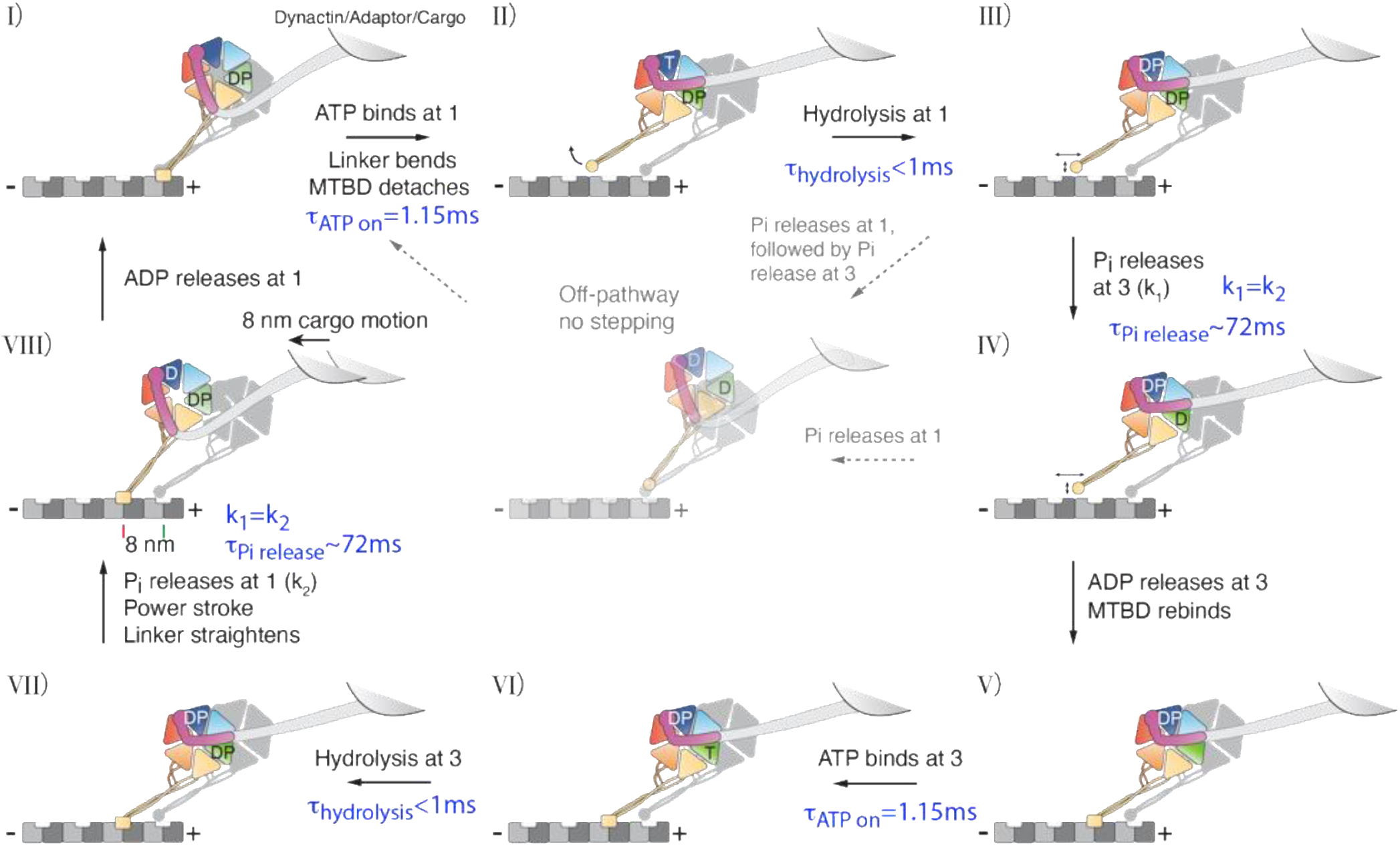
Proposed model for dynein stepping *in vivo*. State. **(I)** is the post power stroke with dynein at AAA1 is in the Apo state with ADP · Pi on AAA3. The rectangular shape of the MTBD denotes the dynein monomer strongly bound to the microtubule. The colored and grey dynein monomer tails are attached to the dynactin/adaptor/cargo structure. **(I) → (II)**: *ATP* binds at AAA1, inducing the gap between AAA1 and AAA2 to close dynein to rapidly unbind from the microtubule. With *ADP · Pi* (denoted here as DP) on the AAA3 site, the closure causes the stalk and buttress to rotate the MTBD so that the dynein enters into the weakly bound state. In this state the linker arm contact to AAA5 is broken. During the brief times when the MTBD is detached from the MT, is free to assume a variety of stress-free configurations as measured by negative stained and cryo-EM microscopy. **(II) → (III)**: *ATP* hydrolyzes to *ADP · Pi* in < 1 *ms* at AAA1. In states (II) and (III), the microtubule binding domain (MTBD) is in the weakly bound state. **(III) → (IV)**: The first rate-limiting step is *Pi* release at AAA3 with a rate constant of *k*_1_. *Starting from state (IV)*, *DoPi departs from the active-cycling model*. **(IV → V)**: ADP releases from AAA3 in ~1 *ms* and allows dynein to rebind to the MT in the *α*-registry. **Greyed-out off-pathway**: If *Pi* releases AAA1 before dynein rebinds to the MT, the power stroke is futile since the MT is still in the weakly bound state. **(V → VI → VII)**: *ATP* rapidly binds to AAA3 in ~1 *ms* and then rapidly hydrolyzes to DP. The rebinding of the microtubule binding domain (MTBD) to the MT is hypothesized to have backward allosteric communication to the *ATP* ase domain. We conjecture that the allosteric interaction causes sensor I loop at AAA1 to break its contact with *ADP* · *Pi* at the AAA1 site, which in turn allows the thermal desorption of *Pi*. **(VII → VIII)**: The release of *Pi* at AAA1 with rate constant *k*_2_ leads to the straightening of the linker and the power stroke that advances dynein by 8 *nm* along the MT. **(VIII → I)**: ADP at AAA1 is released in ~1 *ms* and returns the dynein to state (I). The listed Pi release times of 72 ms is for *T* = 22° *C*. Note that the dynactin/adaptor complex and the vesicle are not explicitly shown in this figure.

The “active-cycling” model requires the hydrolysis of *ATP* only at the AAA1 site for a single step (*64*). In this model, an *ADP* molecule remains bound at AAA3 and opens a gate that allows active cycling of *ATPs* at AAA1 (Fig. S20). In Supplementary Note S1, we present the two main arguments used in support of the active-cycling model, and a re-analysis that argues that the previous data does not favor the active-cycling model over a model that requires the hydrolysis of two *ATPs*. Also, optical tweezer measurements show that only a weak force is needed to detach dynein from the MT with *ADP* bound at AAA3 (Fig. S21), suggesting that *ADP* may not be bound to AAA3 during the power stroke (*65*).

Our *in vivo* step-resolving data reveal dynamics that were hidden in single-step *in vitro* studies taken at much lower *ATP* concentrations. We argue here that a 1-*ATP* model is incompatible with the two equal and thermally activated rate constants *k*(*T*) = *k*_1_(*T*) = *k*_2_(*T*). In considering all of the rates in *ATP* hydrolysis, *Apo* → *ATP* → *ADP* · *P*i → *ADP* → *Apo*, the hydrolysis of *ATP* and the dissociation of *ADP* have been found to occur on timescales of ≤ 1 *ms* in cellular conditions at room temperature (*66*). Single-molecule *in vitro* experiments have measured an *ATP* binding rate constant *k_MT_* = 0.29± 0.10 *μM^-1^ s^-1^ (67*). Assuming 3 *mM* of *ATP* concentration in the axon (*68*–*70*), the *ATP* binding rate is 870 *s*^-1^. Thus, we are led to assign the two rates to be the thermal desorption rates of two *Pi*’*s*, and propose a new model for the chemomechanical cycle of the dynein motor.

Our chemomechanical model of dynein requires the *ATP* hydrolysis on both AAA1 and AAA3 to complete a full step (Fig. 6). We show that the proposed model, abbreviated **DoPi**(for **D**ualdesorption **o**f ***Pi***), integrates much of the data from x-ray, negative-stained and cryo-EM structures, single-molecule motility and force measurements, and biochemical kinetics work with the new findings presented in this paper.

In State (I) AAA1 in the Apo state is strongly bound to the microtubule (*65*, *71*), with the linker arm rigidly connected to AAA1 and AAA5 (*64*). The AAA3 site is occupied by *ADP · Pi*, and allows the linker to bend upon *ATP* binding at AAA1 during the transition (I) → (II). *ATP* binding to AAA1 closes the AAA1-AAA2 cleft (*4*–*6*, *64*), causing the AAA5-stalk to rotate the microtubule binding domain (MTBD) from the strongly-bound (SB) *α*-registry to the weakly-bound (WB) *β*-registry (*72*). Once *ATP* binds to dynein initially in the *Apo* state, the dissociation of dynein from the MT is < 1 *ms* at [*ATP*] = 3 *mM* (*66*). The dynein linker in the ADP-vanadate state was shown to be in straight, partially-bent and fully-bent (*64*), while a large variability of the stalk-tail distance was seen in negatively stained EM structures (*73*). Since the linker remains straight in the AAA3_E/Q_ mutant (that allows *ATP* binding but not hydrolysis), the linker may not bend until *ATP* is hydrolyzed on AAA3 (*64*).

The transition (II) → (III) is believed to occur in < 1 *ms* (*66*), while the thermal desorption of *Pi* from AAA3 (III) → (IV) occurs with a rate *k*_1_(22°C) = 13.1 *s*^-1^. While this assignment is consistent with the rate deduced from optical tweezer measurements of dynein monomers (*67*), we note that the rate constants deduced from Ref. (*67*) explicitly assumes that the AAA1 was the only active site, so their *k*_*M*1_ is a mixture of our *k*_1_ and *k*_2_.

Once the *Pi* at the AAA3 site is released, the transition from (IV) → (V) occurs when *ADP* dissociates with a rate of ~ 1000 .s’^-1^ (*66*). *This transition is a critical step where our model departs from the active-cycling model that requires ADP to remain at AAA3*. We propose that the release of *ADP* from AAA3 opens the cleft between AAA3 and AAA4 (Fig. S22), which then allosterically reorients the dynein stalk by causing the stalk helices to rotate the MTBD into the SB *α*-registry. Optical tweezer force measurements of the dynein AAA3_K/A_ mutant which does not allow *ATP* to bind show that the dynein is strongly-bound to the MT (*65*).

The transitions (V) → (VI) → (VII) occurs quickly due to the high binding rate of *ATP* and its subsequent hydrolysis. Optical-tweezer experiments showed that the dynein AAA3_E/Q_ mutant (which is equivalent to State VI) was strongly bound (*65*, *74*). Our model suggests that States VII and VIII are also strongly bound to the MT binding. This prediction can be tested with single molecule force measurements made under saturating conditions of *ADP* · *Pi*. While there is no structural evidence that ties the linker conformation to the strength of MT binding, we believe that the two can be de-coupled and show the linker to be in the flexible state after dynein becomes strongly bound to the MT.

The transition (VII) → (VIII) occurs when *Pi* leaves the AAA1 site at a rate *k*_2_. After the release of *Pi*, the power stroke occurs (*4*, *6*, *78*, *79*). This assignment is based on structural studies which reveal that the linker is straight and rigid when dynein is saturated with 1.5 mM ADP (*4*, *6*, *78*, *79*). The rapid release of *ADP* from AAA1 returns the dynein back to state (I).

For the efficient operation of the dynein motor, there should be a time ordering of the two sequential *ATP* hydrolysis even though the two *Pi* off-rates are equal. For a productive power stroke, the MTBD should be strongly bound to the MT. If *Pi* prematurely leaves AAA1 before AAA3, the power stroke may cause the MTBD to detach form the MT rather than propelling the vesicle forward, as shown in the transition from (III) to the greyed-out “off pathway no stepping state”. A detailed discussion for the time ordering of *Pi* and supporting evidence is presented below. Sequential and time-ordered hydrolysis of four *ATP* s has also been observed for the DNA packaging motor bacteriophage *ϕ*29 (*75*).

### Molecular stepping via periods of transient binding and momentary diffusion

We present a sequence of proposed molecular motions based on our model in Fig. S23. The hydrolysis of *ATP* at each dynein monomer is assumed to be independent of the other monomer, but the motion of the vesicle is determined by the position of the dynactin and its connection to the cargo vesicle. While in States (II) – (IV), the dynein monomer alternates between being weakly bound to the MT and in a detached diffusing state. In this conformation, the linker and tail connection to dynactin is allows a large range of angular and longitudinal motion. Since there is no discernable diffusion of the vesicle after the previous power stroke, most spatial exploration by the dynein before the next power stroke is believed to occur while its MTBD is detached from the microtubule.

Each dynein monomer spends approximately half of the time in the weakly bound state (States II – III) and the other half the time in the strongly bound state (States V-VIII), but the linker arms remain in the flexible state to allow for more diffusive exploration. The bent linker depicted in States II and VII and the flexible tail structure are believed to allow the distance between the N-terminus attachment to dynactin and tip of the stalk to fluctuate. Thus, out of an average cycle time of 144 *ms*, each monomer spends 142 *ms* in the flexible state. The power stroke (VII→VIII) moves the tail of the stepping dynein forward momentarily (Fig. S23D, E), but the dynein quickly relaxes into the flexible state < 2 *ms* after *ADP* release and the binding of a new *ATP* (States I – II). This transient motion occurs too rapidly to be detected in our experiment, and the observed motion is the compromise position between the stepping and non-stepping monomer. We calculate that the return to this flexible state also positions the MTBD forward by +10.8 *nm*. This distance was derived from the known structural conformations of the dynein monomer and its orientation with respect to the MT in the pre-power stroke *ADP* · *Pi* state and in the post-power stroke *Apo* state.

If the leading and lagging dynein monomer become too separated, it is likely that the tension exerted on the lagging monomer will likely lagging motor to temporarily detach from the MT, and then reattach at a position closer to the motor the just stepped. Fig. S24 outlines how the mutual stress between the two monomers is relieved. Model is consistent with the single molecule force experiments that show very asymmetric detachment forces (*65*). The same stress relief should occur if two independent dynein motor systems become too separated.

We further propose that the range of the diffusive search width of the stepping dynein is twice width of the measured cargo displacement. Since the stepping dynein monomer begins its diffusive search +10.8 *nm* in front of the previous power stroke, the distribution of the next vesicle position is predicted to be +5.4 *nm* in front of the previous step position. The forward positioning of the dynein monomer should also be largely temperature independent since it arises from differences in the structural states of dynein.

The model prediction of +5.4 *nm* is in remarkable agreement with the center of +5.0 *nm* found by fitting a Gaussian curve to step-size histograms (Fig. 5A and Fig. S25). To link our detailed model to the observations, we simulate the detected motion in the presence of very low noise (0.1 *nm*) and with our observed 4 *nm* noise (Fig. S26). The simulated dynein stepping made with 1 *ms* time resolution and 0.1 *nm* spatial noise reveals all of the nano-motion is quantized in 4 *nm* increments. However, when 4 *nm* noise added is added to the simulation, the resulting step-size histogram is in remarkable agreement with our measurements and the previous *in vitro* measurements (*58*, *59*). In future work, these predictions can be tested by tracking the cargo with greatly increased spatial and temporal resolution.

Simulations were also used to investigate the motion of multiple dynein motor systems acting on the same vesicle in the context of our interpretation of the quantization of *ϕ* (Fig. S27). In the case of two motors, the non-stepping motors acts as stabilizing anchors (analogous to the non-stepping dynein monomer in the analysis above), and the net motion would be half to the step taken by the active motor. In the case of *N* independent motors, a step of length *δ_step_*, the motion would be (1/*N*) · *δ_step_*. While the effect of multiple anchors would still appear in the dispersion *s*^2^ → *s*^2^/*N* of the Gaussian distribution, the smaller steps would not be resolved.

The width of the experimental step-size histograms decreases with increasing temperature (Fig. 5D). Qualitatively, the narrowing of the histograms is due to the decreased the time allowed for the dynein to diffusively sample MT binding sites. To further quantitate our molecular model, the diffusive search is assumed to be constrained by an elastic spring *F_spring_*(*t*) = *κx*(*t*), where *κ* is an effective spring constant and *x*(*t*) is the departure from the initial starting position *x*_0_ = + 10 *nm* of the MTBD. The motor system and cargo experience a viscous drag force *F_d_* = *γẋ*(*t*). With these two constraints, the exploration of the MTBD of a dynein monomer detached from the MT is described by a diffusive motion 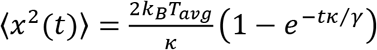 (*76*). In Fig. S25 and Note S2, we show that decreasing with of the step-size histograms is consistent the physical size of the dynein monomer, the viscosity of axonal cytosol, and the time the stepping monomer in State II is detached from the MT.

A lower limit of the force exerted by dynein *during* the power stroke is obtained by concatenating all of the single step data to form an average of all the time traces (Fig. S28). We note that the transition time during a molecular step is completely unresolved with the 1 ms frame rate used to record single steps. If we assume the time duration of the power stroke is less than 0.5 *ms* (Note S4), the lower limit to the minimum force *during* the power stroke force *F_P.S_* > 40 *pN*, which is significantly higher than the unbinding forces in both the forward and backward directions as measured by subjecting a dynein monomer to a slowly linear increasing optical tweezer linear ramping force (*65*, *77*). Higher time resolution measurements are needed to better estimate the instantaneous forces exerted on the MTBD during the power stroke, and to better elucidate the transient kinetic motions.

### Measurement of the randomness parameter supports the model that requires 2 ATPs per cycle

Block and Schnitzer have shown that the “randomness parameter” *r* ≡ (〈*t*^2^〉 – 〈*t*〉^2^〉/〈*t*〉^2^, the normalized distribution of dwell times in an enzymatic cycle is related to the number of kinetic sub-steps within a complete cycle (*78*). For a sequence of *M* forward only steps with rate constants *k_i_*, if *p* steps are comparable and the other *M* – *p* rates are much higher, *r* ~ 1/*p*. In general *r*^-1^ = *n_min_* < *M*, where *n_min_* is the minimum number of kinetic steps of the full kinetic cycle (*79*).

In our live cell measurements, the two thermal desorption rates *ADP* · *Pi* → *ADP* are the primary contributors to *r* = 1/*p* ~ 1/2 (Fig. S29), and all of the other rates are too fast to add to the dwell time dispersion. In the low [*ATP*] regime, if the rate limiting step is due to *ATP* binding at only the AAA1 (*58*, *59*), the *in vitro* experiments should give *r*~ 1, but if a dynein cycle requires 2 *ATPs*, *r*~ 1/2. In numerical simulation (Fig. S30), when the *ATP* on-rate is equal to the thermal desorption rate of *Pi*, *r* → 1/4, since the *ATP* hydrolysis and the *ADP* desorption rates will remain too fast to contribute to *r*. A measurement of *r* as a function of [*ATP*] would clearly differentiate between the 1-*ATP* active cycling model and our proposed 2-*ATP* model.

### The dynein motor system may operate out of thermal equilibrium

Another form of the FT is where fluctuations of the work Δ*W* done on the system in a thermal reservoir of temperature T also satisfies the relation,

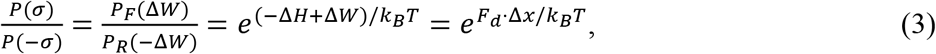

where Δ*H* = 0 is the net change in the Helmholtz free energy over a full enzymatic cycle, Δ*W* = *F*_d_ · Δ*x*, and *F*_d_ = *γv* is the Stokes drag force (*52*). Taking the logarithm of Equation 3 gives 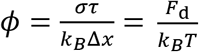. Using the asymptotic value of *ϕ* = 0.1 *nm*^-1^, *F_d_* = *ϕk_B_T* = 0.43 *pN* if *T* = *T_cell_*. Using an axonal viscosity *η_cargo_* = 1.34 *Pa* · *s* (see Note S4), and the range of velocities measured for *ϕ* = 0.1 *nm*^-1^ (Fig. 3F), the average inferred drag force *F_d_* ranges between 0.6 – 7.2 *pN*. The high values of *F_d_* = *ϕk_B_T* suggests that the motor system may be operating at and effective temperature *T_eff_* > *T_cell_*.

Hayashi *et al* (*54*) used the force-velocity relation for describing a very large effective stall force and obtain an effective temperature *T*_eff_ of 4200K. While we believe that the *ϕ* value should not be interpreted as the stall force, we agree with their conclusion that there is evidence that 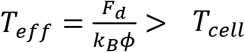. Our analysis of the motion of the cargo vesicle in the context of thermal and non-thermal equilibrium will be discussed in a companion paper.

### Dynein chemo-mechanical efficiency

The ability to determine the number of dynein motor systems used during the transport of individual vesicles allows us to estimate a range of efficiency that the dynein motor converts chemical energy into mechanical work. In segments where *ϕ* = 2*μ*/*s*^2^ = 0.1 *nm*^−1^, the observed dispersion *s*^2^ is interpreted to be due to fluctuations about the mean displacement *μ* arising from a single motor. The efficiency of converting chemical energy to mechanical work, defined here as *ξ* 〈*F_step_* · Δ*x*〉/2Δ*G_ATP_*, where 〈*F_step_* · Δ*x*) is the average work done per step. and 2Δ*G_ATP_* is the free energy of the hydrolysis of two *ATP*’s per stepping cycle. Under neuronal concentrations of [*ATP*], [*ADP*] and [*Pi*] (80, 81), Δ*G_ATP_*~51 *kJ* · *mol*^−1^.

The maximum mechanical work of a molecular motor occurs just below the stall force *F_stall_*. In our measurements, the highest recorded vesicle velocity for *ϕ* = 0.1 *nm*^−1^ is *ν_m_* = 6*μm* · *s*^−1^ (Fig. 3F). Setting *F_stall_* > *F_d_* = 6*πην_m_*, and using our estimated values of the axonal viscosity *η_axon_* = 1.34 *Ps* · *s*, a vesicle radius *a* = 100 *nm*, and the average displacement of 5*nm/step* (the centroid in the step-size histogram), the mechanical work in moving the vesicle is 〈*F_d_* · Δ*x*) ~ 7.58 × 10^−20^*J* · *step*^−1^. Thus, the estimated chemo-mechanical efficiency is of the order of *ξ* ≥ 0.45. The largest uncertainty in this estimate of *ξ* is the effective viscosity *η_axon_* for vesicles with *a* ~ 100 *nm*.

In the conversion of chemical energy into useful mechanical work, a portion of the energy has to overcome the internal “frictional” forces within the motor mechanism. For example, internal energy is dissipated as the stalk helices slide across each other to rotate the MTBD from the weakly-bound into the strongly-bound state. Similarly, in going from State VII to State I, energy is required to change relative angular orientation of the stalk-buttress structure with respect to the AAA ring during the power stroke, and this movement is also expected to dissipate energy within the motor.

A high chemo-mechanical efficiency also demands that the fraction of enzymatic cycles of futile power strokes is minimized (*82*). As previously discussed, this requires a time-ordering of Pi release from the AAA3 and AAA1 sites based on a sequence of allosteric events. Beginning with State I in Fig. 6, *ATP* binding and hydrolysis at the AAA1-AAA2 interface closes the gap. This movement causes the stalk helices to rotate the MTBD and allow it to release from the MT (*83*). We propose that this motion also causes Sensor I Loop at AAA3 to move to break two stabilizing contacts with the *Pi* at the AAA3 site (*84*, *85*). In Fig. S31A, the decrease in potential energy barrier that prevents *Pi* from being released is indicated schematically. After *Pi* thermally desorbs from AAA3, *ADP* quickly leaves, and the AAA3-AAA4 interface opens. The AAA ring motion induces the sliding motion of the stalk helices that rotate the MTBD into the strongly bound State V, consistent with single molecule force studies (*65*).

The thermal desorption of *ADP* at AAA3 cannot provide the energy to rotate the MTBD into to the strongly bound state, and we propose that the binding of *ATP* at AAA3 (State VI) provides the required energy. The newly bound *ATP* at AAA3 also establishes the Sensor I Loop contacts at AAA3 that prevents *Pi* from prematurely leaving the AAA3 site. However, the movement of the stalk helices also allosterically causes Sensor I loop at the AAA1-AAA2 interface to pull away from the AAA1 site protecting *Pi* from thermal desorption as shown in State VIII.

There is structural evidence supporting the time-ordering hypothesis. The structure of dynein obtained in the presence of A?P-vanadate reveal that the AAA3 site contains only ADP *without the vanadate* while the AAA1 site has a stable *ADP-vanadate* (*84*). This is consistent with our measured *Pi* release rate of 13.1 s^−1^, since the *Pi* would have left the AAA3 in the time it takes to form crystals of dynein after *Mg* · *ATP* and Na_3_VO_4_ are added. Examination of the environment surrounding the binding pocket of *ADP* at AAA3 and AAA1 shows that the *Pi* at AAA1 is stabilized by two contacts with the Sensor 1 (S1) Loop, while the S1 Loop at AAA3 has swung away from the binding site (*84*, *85*).

In Fig. 6, we note that both the States (III) and (VII) have *ADP* · *Pi* bound to AAA1 and AAA3, but they are in different MT binding states. The hysteresis in the dynein cycle is proposed to be the result of the allosteric interactions that affect the positioning of the S1 Loops at AAA1 and AAA3. In summary, in States (VII), (VIII) and (I), the S1 Loop at AAA3 is taken to be closed based on the AMPPNP structure (*64*). The S1 Loop contacts at AAA3 are proposed to be broken when ATP binds to AAA1 in the transition (I) → (II). Similarly, the S1 Loop contacts at AAA1 are broken after *ADP* releases from AAA3. The sum result is that *Pi* has to desorb from AAA3 before *Pi* is allowed to leave AAA1, and *Pi* has to desorb from AAA1 before *Pi* on AAA3 is allowed to leave the AAA3 site. It may be possible to falsify this hypothesis by determining the positions of Sensor I Loop at AAA1 and AAA3 in different nucleotide states with appropriate mutants and with specific time-ordering of *ATP, ADP* · *Pi* or *ADP* to capture the hysteresis.

## Concluding remarks

In *What is Life*, Erwin Schrodinger conjectured that the operation biological nanoscale machines “will be is different from a anything we have yet tested in the physical laboratory” (*86*). Unlike macroscopic machines which are constructed to minimize friction, biomolecular machines are embedded in an environment where fluctuations and dissipation dominate (*87*).

Rather than fighting friction, dynein exploits thermal fluctuations as an essential part of its operation in two different ways. Specifically, the Arrhenius activation probabilities with Sensor I Loop in its different positions is indicated schematically by the solid and dotted curves in Fig. S31A. The thermal desorption rates are proportional to *e^-E′_a_/k_B_T^* and *e*^-(*E_a_ + E_loop_)/k_B_T*^. If the thermal desorption of *Pi* from each AAA site takes ~ 10 *ms* at 37°C when Sensor I loop is not providing stabilizing contacts, the off-rate when Sensor I Loop, the probability of the incorrect time-ordering will occur (∝*e*^−(*E_loop_)/k_B_T*)^ is exponentially unlikely.

We further conjecture that the motions due to thermal fluctuations also provides a considerable fraction of the mechanical movement due to a motor step. In the pre-power stroke state, thermal drives the spatial exploration of the stepping dynein. During this time, a flexible linker arm allows considerable movement between the AAA ring and the dynactin that links the two dynein monomers, as portrayed schematically by the upper potential energy curve in Fig. S31B. When *Pi* is released from the AAA1 site, the potential energy is suggested to change into the potential energy surface that allows the linker arm to reattach to AAA5, but the movement that straightens the linker is driven by thermal fluctuations rather than the conversion of chemical energy into mechanical motion.

In Fig. S23, we have argued that the known molecular structures suggest that the AAA ring is moved forward by ~ 10.8 *nm* relative to its previous location. This distance is due to a combination of the change in the linker arm length and the change in angle between stalk and the MT from 31° to 55°.angle. The distance *l* ~ 15 *nm* between the AAA ring and the end of the MTBD adds a net forward movement of *l*(cos 55°– cos 31°) = 4.3 *nm*. Thus, the long length of the stalk serves to magnify the small change of distance of the AAA1-AAA2 interface into a large forward motion of the AAA ring, while the remainder of the net forward motion is supplied by Brownian motion.

A related mechanism of movement magnification was discussed in single molecule FRET studies of codon recognition of the ribosome during translation (*87*, *88*). A critical element in selecting correct ternary complex consisting of aminoacyl-tRNA, GTP, and elongation factor Tu(EF-Tu) is the “induced fit” between the decoding site of the ribosome and the tRNA. The induced fit mechanism was shown to position EF-Tu at the other end of the t-RNA to be more likely to make contact with the Sarcin-Ricin loop of the 50S subunit that triggers GTP hydrolysis. The large linear distance between the anti-codon region and the t-RNA EF-Tu complex amplifies the small molecular motion at the 30S. Also, a slight difference in positioning of EF-Tu results in an exponentially different probabilities of accommodation of t-RNA, and that the difference in positioning and thermal fluctuations constitutes the primary mechanism for the initial selection of tRNA.

In summary, the long-term tracking of individual vesicles in live cells with molecular step resolution has provided evidence suggesting that dynein requires the hydrolysis of two ATPs per step. Precise measurements of the statistical fluctuations of *single* vesicles are used to determine the number of motors used during the retrograde transport, and supports a detailed and falsifiable molecular model that is consistent with much of the previous structural, single-molecule force and biochemical kinetic data.

## Methods summary

Rare-earth doped upconversion nanoparticles were synthesized using protocols modified from (*30*). UCNPs were functionalized with a layer of silica, followed by conjugation to PEG500 and 10% PEG3400-biotin. For axonal transport experiments, the biotinylated-UCNPs were further conjugated to wheat germ agglutinin to target glycoproteins on axons. Human induced neurons were generated from human embryonic stem cells (H1 cell line) using a single transcription factor Neurogenin 2 (*48*). Neurons were cultured in microfluidic chambers and their axons grew through the 900-*μm* long microchannels. UCNPs were incubated in the axon chamber and imaging was done inside the microchannels. Single UCNP imaging was performed using a home-built wide-field microscope with a 976 nm fiber laser, and a 100X, NA = 1.49 oil-immersion objective (*30*). For fluctuation theorem analysis, single cargoes were imaged at 10 millisecond time resolution. Single dynein-step measurements were performed with 2.5 millisecond time resolution at 22 °*C* and 30 °*C*, and 1 millisecond time resolution at 37 °*C*. Detailed synthesis, functionalization, imaging methods, and data analyses are described in the materials and methods section of the supplementary materials.

## Supporting information

SI for SPT-neuron 1-5-22_V5

## Acknowledgements

We acknowledge the funding from the Gordon and Betty Moore Foundation (No. 4309), Stanford Neurosciences Institute (No. 119600), the National Institutes of Health (1R01GM128089-01A1). TEM and SEM imaging were performed at Stanford Microscopy Facility. The project described was supported, in apart, by ARRA Award Number 1S10RR026780-01 from the National Center for Research Resources (NCRR). Its contents are solely the responsibility of the authors and do not necessarily represent the official views of the NCRR or the National Institutes of Health. C.S.P. was supported by Stanford Cancer Translational Nanotechnology Training Grant T32 CA196585 funded by the National Cancer Institute, and NIH Grant 5K99AG065516 funded by the National Institute on Aging. The authors thank Luke Kaplan for assistance during the initial stage of axonal transport experiments. The authors thank Xianchuang Zheng, Guosheng Song, Jianghong Rao for early advice on functionalization of UCNPs. The authors thank Tianshe Yang for taking measurements of the UCNP spectra and X-ray diffraction.

C.S.P. and S.C. conceived the project, designed experiments, analyzed the data and interpreted the results. C.S.P. functionalized upconversion nanoparticles. C.S.P. and Y.Z. performed primary neuron dissection and imaging experiments. Q.L. synthesized upconversion nanoparticles. G.E.M contributed helpful discussions in the analysis. Y.-W.A.H. generated human induced neurons. T.C.S. supervised research on human induced neurons. B.C. supervised axonal transport experiments. C.S.P., T.C.S., B.C. and S.C. wrote the manuscript.

## References

1. S. Millecamps, J. P. Julien, Axonal transport deficits and neurodegenerative diseases. Nature reviews. Neuroscience 14, 161–176 (2013).

2. I. R. Gibbons, A. J. Rowe, Dynein: A Protein with Adenosine Triphosphatase Activity from Cilia. Science 149, 424–426 (1965).

3. B. M. Paschal, H. S. Shpetner, R. B. Vallee, MAP 1C is a microtubule-activated ATPase which translocates microtubules in vitro and has dynein-like properties. The Journal of cell biology 105, 1273–1282 (1987).

4. A. J. Roberts, T. Kon, P. J. Knight, K. Sutoh, S. A. Burgess, Functions and mechanics of dynein motor proteins. Nature reviews. Molecular cell biology 14, 713–726 (2013).

5. G. Bhabha, G. T. Johnson, C. M. Schroeder, R. D. Vale, How Dynein Moves Along Microtubules. Trends in Biochemical Sciences 41, 94–105 (2016).

6. M. A. Cianfrocco, M. E. DeSantis, A. E. Leschziner, S. L. Reck-Peterson, Mechanism and regulation of cytoplasmic dynein. Annual review of cell and developmental biology 31, 83–108 (2015).

7. S. L. Reck-Peterson et al., Single-Molecule Analysis of Dynein Processivity and Stepping Behavior. Cell 126, 335–348 (2006).

8. N. D. Derr et al., Tug-of-war in motor protein ensembles revealed with a programmable DNA origami scaffold. Science 338, 662–665 (2012).

9. L. Urnavicius et al., Cryo-EM shows how dynactin recruits two dyneins for faster movement. Nature 554, 202–206 (2018).

10. V. Belyy et al., The mammalian dynein-dynactin complex is a strong opponent to kinesin in a tug-of-war competition. Nature cell biology 18, 1018–1024 (2016).

11. S. Niekamp, N. Stuurman, N. Zhang, R. D. Vale, Three-color single-molecule imaging reveals conformational dynamics of dynein undergoing motility. Proc Natl Acad Sci U S A 118, (2021).

12. J. Huang, Anthony, Andres, Samara, Lis1 Acts as a “Clutch” between the ATPase and Microtubule-Binding Domains of the Dynein Motor. Cell 150, 975–986 (2012).

13. R. J. McKenney, W. Huynh, M. E. Tanenbaum, G. Bhabha, R. D. Vale, Activation of cytoplasmic dynein motility by dynactin-cargo adapter complexes. Science 345, 337–341 (2014).

14. P. A. Sims, X. S. Xie, Probing dynein and kinesin stepping with mechanical manipulation in a living cell. Chemphyschem: a European journal of chemical physics and physical chemistry 10, 1511–1516 (2009).

15. B. Cui et al., One at a time, live tracking of NGF axonal transport using quantum dots. Proc Natl Acad Sci U S A 104, 13666–13671 (2007).

16. L. Kaplan, A. Ierokomos, P. Chowdary, Z. Bryant, B. Cui, Rotation of endosomes demonstrates coordination of molecular motors during axonal transport. Science advances 4, e1602170 (2018).

17. S. E. Encalada, L. Szpankowski, C. H. Xia, L. S. Goldstein, Stable kinesin and dynein assemblies drive the axonal transport of mammalian prion protein vesicles. Cell 144, 551–565 (2011).

18. A. K. Rai, A. Rai, A. J. Ramaiya, R. Jha, R. Mallik, Molecular adaptations allow dynein to generate large collective forces inside cells. Cell 152, 172–182 (2013).

19. Y. M. Sigal, R. Zhou, X. Zhuang, Visualizing and discovering cellular structures with super-resolution microscopy. Science 361, 880–887 (2018).

20. I. Rasnik, S. A. McKinney, T. Ha, Nonblinking and long-lasting single-molecule fluorescence imaging. Nat Methods 3, 891–893 (2006).

21. R. B. Altman et al., Cyanine fluorophore derivatives with enhanced photostability. Nature Methods 9, 68–71 (2012).

22. T. A. Tsunoyama et al., Super-long single-molecule tracking reveals dynamic-anchorage-induced integrin function. Nature chemical biology 14, 497–506 (2018).

23. M. E. Tanenbaum, L. A. Gilbert, L. S. Qi, J. S. Weissman, R. D. Vale, A protein-tagging system for signal amplification in gene expression and fluorescence imaging. Cell 159, 635–646 (2014).

24. H. Liu et al., Visualizing long-term single-molecule dynamics in vivo by stochastic protein labeling. Proc Natl Acad Sci U S A 115, 343–348 (2018).

25. R. P. Ghosh et al., A fluorogenic array for temporally unlimited single-molecule tracking. Nature chemical biology 15, 401–409 (2019).

26. G. Xu et al., New Generation Cadmium-Free Quantum Dots for Biophotonics and Nanomedicine. Chem Rev 116, 12234–12327 (2016).

27. S. Wu et al., Non-blinking and photostable upconverted luminescence from single lanthanide-doped nanocrystals. Proceedings of the National Academy of Sciences 106, 10917–10921 (2009).

28. D. J. Gargas et al., Engineering bright sub-10-nm upconverting nanocrystals for single-molecule imaging. Nat Nanotechnol 9, 300–305 (2014).

29. Y. Liu et al., Amplified stimulated emission in upconversion nanoparticles for super-resolution nanoscopy. Nature 543, 229–233 (2017).

30. Q. Liu et al., Single upconversion nanoparticle imaging at sub-10 W cm2 irradiance. Nature Photonics 12, 548–553 (2018).

31. B. Tian et al., Low irradiance multiphoton imaging with alloyed lanthanide nanocrystals. Nature Communications 9, 3082 (2018).

32. F. Wang et al., Microscopic inspection and tracking of single upconversion nanoparticles in living cells. Light: Science & Applications 7, 18007 (2018).

33. J. Zhao et al., Single-nanocrystal sensitivity achieved by enhanced upconversion luminescence. Nat Nanotechnol 8, 729–734 (2013).

34. F. Wang et al., Tuning upconversion through energy migration in core-shell nanoparticles. Nat Mater 10, 968–973 (2011).

35. Y. Zhang et al., Multicolor Barcoding in a Single Upconversion Crystal. Journal of the American Chemical Society 136, 4893–4896 (2014).

36. L. Zhou et al., Single-band upconversion nanoprobes for multiplexed simultaneous in situ molecular mapping of cancer biomarkers. Nat Commun 6, 6938 (2015).

37. G. Chen et al., (alpha-NaYbF4:Tm(3+))/CaF2 core/shell nanoparticles with efficient near-infrared to near-infrared upconversion for high-contrast deep tissue bioimaging. ACS nano 6, 8280–8287 (2012).

38. S. Chen et al., Near-infrared deep brain stimulation via upconversion nanoparticle-mediated optogenetics. Science 359, 679–684 (2018).

39. S. H. Nam et al., Long-term real-time tracking of lanthanide ion doped upconverting nanoparticles in living cells. Angew Chem Int Ed Engl 50, 6093–6097 (2011).

40. H. S. Han et al., Spatial charge configuration regulates nanoparticle transport and binding behavior in vivo. Angew Chem Int Ed Engl 52, 1414–1419 (2013).

41. R. Abdul Jalil, Y. Zhang, Biocompatibility of silica coated NaYF(4) upconversion fluorescent nanocrystals. Biomaterials 29, 4122–4128 (2008).

42. N. Monnier et al., Inferring transient particle transport dynamics in live cells. Nat Methods 12, 838–840 (2015).

43. Y. Gu et al., Rotational dynamics of cargos at pauses during axonal transport. Nat Commun 3, 1030 (2012).

44. S. Yogev, R. Cooper, R. Fetter, M. Horowitz, K. Shen, Microtubule Organization Determines Axonal Transport Dynamics. Neuron 92, 449–460 (2016).

45. D. Bray, M. B. Bunge, Serial analysis of microtubules in cultured rat sensory axons. Journal of neurocytology 10, 589–605 (1981).

46. A. G. Hendricks et al., Motor coordination via a tug-of-war mechanism drives bidirectional vesicle transport. Current biology: CB 20, 697–702 (2010).

47. S. L. Reck-Peterson, W. B. Redwine, R. D. Vale, A. P. Carter, The cytoplasmic dynein transport machinery and its many cargoes. Nature reviews. Molecular cell biology 19, 382–398 (2018).

48. Y. A. Huang, B. Zhou, M. Wernig, T. C. Sudhof, ApoE2, ApoE3, and ApoE4 Differentially Stimulate APP Transcription and Abeta Secretion. Cell 168, 427–441.e421 (2017).

49. R. H. Miller, R. J. Lasek, Cross-bridges mediate anterograde and retrograde vesicle transport along microtubules in squid axoplasm. The Journal of cell biology 101, 2181–2193 (1985).

50. G. T. Shubeita et al., Consequences of motor copy number on the intracellular transport of kinesin-1-driven lipid droplets. Cell 135, 1098–1107 (2008).

51. P. D. Chowdary et al., Nanoparticle-assisted optical tethering of endosomes reveals the cooperative function of dyneins in retrograde axonal transport. Scientific reports 5, 18059 (2015).

52. G. E. Crooks, Entropy production fluctuation theorem and the nonequilibrium work relation for free energy differences. Physical review. E, Statistical physics, plasmas, fluids, and related interdisciplinary topics 60, 2721–2726 (1999).

53. G. M. Wang, E. M. Sevick, E. Mittag, D. J. Searles, D. J. Evans, Experimental demonstration of violations of the second law of thermodynamics for small systems and short time scales. Physical review letters 89, 050601 (2002).

54. K. Hayashi, Y. Tsuchizawa, M. Iwaki, Y. Okada, Application of the fluctuation theorem for non-invasive force measurement in living neuronal axons. Molecular biology of the cell, mbcE18010022 (2018).

55. K. Hayashi, H. Ueno, R. Iino, H. Noji, Fluctuation theorem applied to F1-ATPase. Physical review letters 104, 218103 (2010).

56. M. D. Springer, W. E. Thompson, The Distribution of Products of Independent Random Variables. SIAM Journal on Applied Mathematics 14, 511–526 (1966).

57. P. A. Bromiley, Products and Convolutions of Gaussian Probability Density Functions. Tina-Vis. Memo, 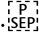; 3, pp. 1–13 (2014).

58. M. A. DeWitt, A. Y. Chang, P. A. Combs, A. Yildiz, Cytoplasmic dynein moves through uncoordinated stepping of the AAA+ ring domains. Science 335, 221–225 (2012).

59. S. L. Reck-Peterson et al., Single-molecule analysis of dynein processivity and stepping behavior. Cell 126, 335–348 (2006).

60. C. Kural et al., Kinesin and dynein move a peroxisome in vivo: a tug-of-war or coordinated movement? Science 308, 1469–1472 (2005).

61. X. Nan, P. A. Sims, X. S. Xie, Organelle tracking in a living cell with microsecond time resolution and nanometer spatial precision. Chemphyschem: a European journal of chemical physics and physical chemistry 9, 707–712 (2008).

62. C. Cho, S. L. Reck-Peterson, R. D. Vale, Regulatory ATPase sites of cytoplasmic dynein affect processivity and force generation. The Journal of biological chemistry 283, 25839–25845 (2008).

63. H. Schmidt, A. P. Carter, Review: Structure and mechanism of the dynein motor ATPase. Biopolymers 105, 557–567 (2016).

64. G. Bhabha et al., Allosteric communication in the dynein motor domain. Cell 159, 857–868 (2014).

65. M. P. Nicholas et al., Cytoplasmic dynein regulates its attachment to microtubules via nucleotide state-switched mechanosensing at multiple AAA domains. Proc Natl Acad Sci U S A 112, 6371–6376 (2015).

66. E. L. F. Holzbaur, K. A. Johnson, Microtubules accelerate ADP release by dynein. 28, 7010–7016 (1989).

67. Y. Kinoshita, T. Kambara, K. Nishikawa, M. Kaya, H. Higuchi, Step Sizes and Rate Constants of Single-headed Cytoplasmic Dynein Measured with Optical Tweezers. Scientific Reports 8, (2018).

68. A. Trevisiol et al., Monitoring ATP dynamics in electrically active white matter tracts. eLife 6, (2017).

69. L. Y. Shields, B. A. Mendelsohn, K. Nakamura, in Techniques to Investigate Mitochondrial Function in Neurons, S. Strack, Y. M. Usachev, Eds. (Springer New York, New York, NY, 2017), pp. 115–131.

70. D. Pathak et al., The Role of Mitochondrially Derived ATP in Synaptic Vesicle Recycling. Journal of Biological Chemistry 290, 22325–22336 (2015).

71. K. Imamula, T. Kon, R. Ohkura, K. Sutoh, The coordination of cyclic microtubule association/dissociation and tail swing of cytoplasmic dynein. 104, 16134–16139 (2007).

72. A. J. Roberts, T. Kon, P. J. Knight, K. Sutoh, S. A. Burgess, Functions and mechanics of dynein motor proteins. Nature Reviews Molecular Cell Biology 14, 713–726 (2013).

73. M. L. W. Stan A. Burgess*, Hitoshi Sakakibara†, Peter J. Knight* & Kazuhiro Oiwa, < Burgess-2003-Dynein-structure-and-power-stroke.pdf>. Nature 421, 715–718 (2003).

74. M. A. DeWitt, C. A. Cypranowska, F. B. Cleary, V. Belyy, A. Yildiz, The AAA3 domain of cytoplasmic dynein acts as a switch to facilitate microtubule release. Nature structural & molecular biology 22, 73–80 (2015).

75. J. R. Moffitt et al., Intersubunit coordination in a homomeric ring ATPase. Nature 457, 446–450 (2009).

76. S. F. Nørrelykke, H. Flyvbjerg, Harmonic oscillator in heat bath: Exact simulation of time-lapse-recorded data and exact analytical benchmark statistics. Physical Review E 83, (2011).

77. L. Rao, F. Berger, M. P. Nicholas, A. Gennerich, Molecular mechanism of cytoplasmic dynein tension sensing. Nature communications 10, (2019).

78. M. J. Schnitzer, S. M. Block, Statistical kinetics of processive enzymes. Cold Spring Harb Symp Quant Biol 60, 793–802 (1995).

79. J. R. Moffitt, C. Bustamante, Extracting signal from noise: kinetic mechanisms from a Michaelis-Menten-like expression for enzymatic fluctuations. Febs j 281, 498–517 (2014).

80. M. Tantama, J. R. Martinez-Francois, R. Mongeon, G. Yellen, Imaging energy status in live cells with a fluorescent biosensor of the intracellular ATP-to-ADP ratio. Nature communications 4, 2550 (2013).

81. M. E. I. A. Silver, A TP and Brain Function Maria. Journal Cerebral Blood Flow and Metabolism 9, 2–19 (1989).

82. A. Mogilner, T. C. Elston, H. Wang, G. Oster, in Computational Cell Biology, C. P. Fall, E. S. Marland, J. M. Wagner, J. J. Tyson, Eds. (Springer New York, New York, NY, 2002), pp. 320–353.

83. W. B. Redwine et al., Structural basis for microtubule binding and release by dynein. Science 337, 1532–1536 (2012).

84. H. Schmidt, R. Zalyte, L. Urnavicius, A. P. Carter, Structure of human cytoplasmic dynein-2 primed for its power stroke. Nature 518, 435–438 (2015).

85. T. Kon et al., The 2.8 Å crystal structure of the dynein motor domain. Nature 484, 345–350 (2012).

86. E. Schrodinger, What is Life?: With Mind and Matter and Autobiographical Sketches. Canto Classics (Cambridge University Press, Cambridge, 2012).

87. S. Chu, Is Life Based on the Law of Physics? M. L. C. R.Y. Chao, A.J. Leggett, W.D. Phillipos C.L. Harper, Ed., (Cambridge University Press, Cambridge England, 2011).

88. T. H. Lee, S. C. Blanchard, H. D. Kim, J. D. Puglisi, S. Chu, The role of fluctuations in tRNA selection by the ribosome. Proceedings of the National Academy of Sciences 104, 13661–13665 (2007).

