## Supplementary material for "Nanometer-Resolution Long-term Tracking of Single Cargos Reveals Dynein Motor Mechanisms": SI for SPT-neuron 1-5-22_V5

### Materials and Methods

Chemicals and reagents

All starting materials were purchased from commercial supplies. The rare-earth salt $YCl_3 \cdot 6H_2O$, $YbCl_3 \cdot 6H_2O$, $ErCl_3 \cdot 6H_2O$, $TmCl_3 \cdot 6H_2O$ and other chemicals $NH_4F$, NaOH, sodium oleate, cyclohexane, ethanol, octadecene (ODE) (>90%), oleic acid (OA) (>90%), Igepal CO-520, tetraethyl orthosilicate (TEOS), ammonia solution (28%) was purchased from Sigma-Aldrich. Streptavidin coated QD705 were purchased from Thermo Fisher Scientific. Silane-PEG(3.4k)-biotin was purchased from Laysan Bio, Inc. 2-[Methoxy-(polyethyleneoxy)propyl] trimethoxysilane was purchased from Gelest. FluoReporter Cell-Surface Biotinylation Kit (F-20650) was purchased from Molecular Probes. All chemical reagents of analytical grade were used directly without any further purification.

Nanoparticle synthesis

**Synthesis of 22 nm $NaYF_4$:$Yb_{0.2}Er_{0.02}$ used in single-particle binding assays:** This type of UCNPs were synthesized and reported in our previous publication (*1*). Given amounts of $YCl_3 \cdot 6H_2O$ (0.78 mmol), $YbCl_3 \cdot 6H_2O$ (0.20 mmol) and $ErCl_3 \cdot 6H_2O$ (0.02 mmol) were added into a 100 mL three-necked flask with 6 mL oleic acid (OA) and 15 mL 1-octadecene (ODE) inside. The mixture was heated to 160 $°C$ to form a clear solution, then cooled down to room temperature. Two milliliters of methanol solution containing NaOH (2.5 mmol) and $NH_4F$ (4 mmol) was added. The mixture was stirred for 30 min at room temperature, and then heated to 120 $°C$ and kept for another 30 min. Subsequently, the solution was heated to 300 $°C$ and maintained for 1 h in an argon atmosphere. After the solution was cooled naturally, 25 mL ethanol was added, and the resulting mixture was centrifugally separated (12000 rpm for 10 min) to a compact pellet, and the supernatant was discarded. The products were collected and washed with cyclohexane and ethanol (40 mL, 1:1, v/v) three times. The UCNPs (diameter 22.9 nm ± 0.9 nm) was stored in 3 mL cyclohexane.

**Synthesis of 29 nm core-shell-shell $NaYF_4$@$NaYbF_4$: 8% Er@$NaYF_4$ used in axonal transport experiments:** The CSS $NaYF_4$@$NaYbF_4$: 8% Er@$NaYF_4$ UCNPs were synthesized in our previous report (*1*).

**Synthesis of 66 nm $NaYbF_4$: 10% Gd, 8% Er@$NaYbF_4$: 8% Er@$NaYF_4$ used for step-size measurements:** Given amounts of $YCl_3·6H_2O$ (0.82 mmol), $GdCl_3·6H_2O$ (0.10 mmol) and $ErCl_3·6H_2O$ (0.08 mmol) were added into a 100 mL three-necked flask with 6 mL oleic acid (OA) and 15 mL 1-octadecene (ODE) inside. The mixture was heated to 160 $°C$ to form a clear solution, then cooled down to room temperature and 2 mL of methanol solution containing NaOH (2.5 mmol) and $NH_4F$ (4 mmol) were added. The mixture was stirred for 30 min at room temperature, then heated to 120 $°C$ and kept for another 30 min. Subsequently, the solution was heated to 300 $°C$ and maintained for 2 h in an argon atmosphere. After the solution was cooled naturally, 25 mL ethanol was added, and the resulting mixture was centrifugally separated (12000 rpm for 10 min) to a compact pellet, and the supernatant was discarded. The products were collected and washed with cyclohexane and ethanol (40 mL, 1:1, v/v) three times. The $NaYbF_4$: 10% Gd, 8% Er (diameter of 46.7 nm ± 1.0 nm) was stored in 3 mL cyclohexane. $Gd^{3+}$ doping was used for controlling size distribution and obtaining uniform UCNPs.

$NaYbF_4$: 10% Gd, 8% Er@$NaYbF_4$: 8% Er was synthesized using the following procedure: 0.40 mmol $RECl_3·6H_2O$ (92% $Yb^{3+}$, 8% $Er^{3+}$) was added into 3 mL oleic acid and 8 mL octadecene in a 100 mL three-neck flask. The solution was heated to 160 $°C$ to form a clear solution. After the mixture was cooled to room temperature, the prepared $NaYbF_4$: 10% Gd, 8% Er in cyclohexane and 2 mL methanol solution containing 1 mmol NaOH and 1.5 mmol $NH_4F$ were added into the reaction flask and stirred for 30 min. The solution was heated to 120 $°C$ to remove low-boiling solvents for 30 min, and then heated to 300 $°C$ and maintained for 1 hour under argon atmosphere. The subsequent purification steps are the same as used for $NaYF_4$: 20% Yb, 2% Er. The $NaYbF_4$: 10% Gd, 8% Er@$NaYbF_4$: 8% Er (diameter of 55.2 nm ± 1.8 nm) was stored in 3 mL cyclohexane.

The inactive $NaYF_4$ layer was synthesized following a similar procedure with that of $NaYbF_4$: 10% Gd, 8% Er@$NaYbF_4$: 8% Er. 0.40 mmol $YCl_3·6H_2O$ was used instead of 0.40 mmol $RECl_3·6H_2O$ (92% $Yb^{3+}$, 8% $Er^{3+}$), and $NaYbF_4$: 10% Gd, 8% Er@$NaYbF_4$: 8% Er was used instead of $NaYbF_4$: 10% Gd, 8% Er. The prepared $NaYbF_4$: 10% Gd, 8% Er@$NaYbF_4$: 8% Er@$NaYF_4$ (diameter of 65.2 nm ± 1.9 nm) was stored in 3 mL cyclohexane.

**Synthesis of 160 nm $NaYbF_4$: 10% Gd, 8% Er@$NaYbF_4$: 8% Er@$NaYbF_4$: 8% Er@$NaYF_4$ used for step-size measurements:**

$NaYbF_4$: 10% Gd, 8% Er: Given amounts of $YbCl_3·6H_2O$ (0.82 mmol), $GdCl_3·6H_2O$ (0.10 mmol) and $ErCl_3·6H_2O$ (0.08 mmol) were added into a 100 mL three-necked flask with 4 mL oleic acid (OA) and 10 mL 1-octadecene (ODE) inside. The mixture was heated to 160 ºC to form a clear solution, then cooled down to room temperature and 2 mL of methanol solution containing NaOH (2.5 mmol) and $NH_4F$ (4 mmol) were added. The mixture was stirred for 30 min at room temperature, then heated to 120 ºC and kept for another 30 min. Subsequently, the solution was heated to 300 ºC and maintained for 2 h in an argon atmosphere. After the solution was cooled naturally, 25 mL ethanol was added, and the resulting mixture was centrifugally separated (12000 rpm for 10 min) to a compact pellet, and the supernatant was discarded. The products were collected and washed with cyclohexane and ethanol (40 mL, 1:1, v/v) three times. The resulting $NaYbF_4$: 10% Gd, 8% Er nanocrystals were stored in 3 mL cyclohexane. The nanoparticles are hexagonal prisms with diameter of ~80 nm and height of 60 nm. $Gd^{3+}$ doping was used for controlling size distribution and obtaining uniform UCNPs.

NaYbF$_4$: 10% Gd, 8% Er@NaYbF$_4$: 8% Er: 0.40 mmol RECl$_3$·6H$_2$O (92% $Yb^{3+}$, 8% $Er^{3+}$) was added into 3 mL oleic acid and 8 mL octadecene in a 100 mL three-neck flask. The solution was heated to 160 °C to form a clear solution. After the mixture was cooled to room temperature, 1.5 mL of the prepared NaYbF$_4$: Gd$_{0.1}$, Er$_{0.08}$ in cyclohexane and 2 mL methanol solution containing 1 mmol NaOH and 1.5 mmol NH$_4$F were added into the reaction flask and stirred for 30 min. The solution was heated to 120 °C to remove low-boiling solvents for 30 min, and then heated to 300 °C and maintained for 1 hour under argon atmosphere. The subsequent purification steps were the same as used for $NaY_{0.78}F_4$:$Yb_{0.2}Er_{0.02}$. The resulting NaYbF$_4$: 10% Gd, 8% Er@NaYbF$_4$: 8% Er nanocrystals were stored in 3 mL cyclohexane. The nanoparticles have diameter of ~100 nm and height of 70 nm.

NaYbF$_4$: 10% Gd, 8% Er@NaYbF$_4$: 8% Er@NaYbF$_4$: 8% Er: the nanoparticles were synthesized by a similar procedure used for NaYbF$_4$: 10% Gd, 8% Er@NaYbF$_4$: 8% Er, except that NaYbF$_4$: 10% Gd, 8% Er@NaYbF$_4$: 8% Er was used as the precursor. The resulting hexagonal prisms have diameter of ~130 nm and height of 80 nm.

Finally, the inactive NaYF$_4$ layer was added using a similar procedure used for synthesizing NaYbF$_4$: 10% Gd, 8% Er@NaYbF$_4$: 8% Er, with 0.40 mmol YCl$_3$·6H$_2$O replacing 0.40 mmol RECl$_3$·6H$_2$O (92% $Yb^{3+}$, 8% $Er^{3+}$), and NaYbF$_4$: 10% Gd, 8% Er@NaYbF$_4$: 8% Er replacing NaYbF$_4$: 10% Gd, 8% Er. The purified NaYbF$_4$: 10% Gd, 8% Er@NaYbF$_4$: 8% Er@NaYbF$_4$: 8% Er@NaYF$_4$ was stored in 3 mL cyclohexane. The hexagonal prism UCNPs have diameter of ~160 nm diameter and height of 90 nm.

Nanoparticle characterization

X-ray diffraction (XRD) measurements were performed on a Bruker Single Crystal Diffractometer D8 Venture (Mo Kα radiation, λ = 0.70930 Å). The size and morphology of UCNP were determined at 100 kV using a JEOL JEM-1400 TEM. The prepared samples were dispersed in cyclohexane and dropped onto the surface of a copper grid for TEM analysis. The upconversion luminescence emission spectra were recorded on an Edinburgh LFS-920 instrument, but the excitation source using an external 0 - 1 W adjustable 980 nm semiconductor laser (Beijing Hi-Tech Optoelectronic Co., China) with an optic fiber accessory, instead of the Xeon source in the spectrophotometer. Upconversion luminescence lifetime was measured with a phosphorescence lifetime spectrometer (FSP920-C, Edinburgh) equipped with a tunable mid-band OPO pulse laser as excitation source (410-2400 nm, 10 Hz, pulse width ≤ 5 ns, Vibrant 355II, OPOTEK). All the photoluminescence studies were carried out at room temperature.

Silica coating of UCNPs

UCNPs were coated with silica using a reverse microemulsion method (*2, 3*). The protocol was modified for different starting materials of UCNPs as shown in Table S1. The protocol used for silica coating of the core-only NaYF$_4$: 20% Yb, 2 % Er is described in details here. Igepal CO-520 (1000 mg) was added to 10 mL cyclohexane and dispersed using sonication for 10 min. Then 200 µL of as-prepared oleic acid-capped UCNPs solution was added and the mixture was stirred vigorously for 1 hour. Subsequently, 150 µL of ammonia solution (28%) was added and the mixture was stirred overnight until a transparent emulsion was formed. TEOS (12 µL) was then added and the mixture was gently stirred for 2 days. The entire process was carried out at room temperature. Silica coated UCNPs were precipitated by adding 10 mL of ethanol and were collected by centrifugation (21,000 g, 20 min). The pellet was dispersed in 40 mL of ethanol by sonication and then collected by centrifugation (21,000 g, 20 min). The washing step was repeated again and the final purified nanoparticles were re-dispersed in 3 mL of ethanol for storage.

Surface modification of silica coated UCNPs

To incorporate PEG onto silica-coated UCNPs, 20 mg of 2-[Methoxy-(polyethyleneoxy)propyl] trimethoxysilane (silane-mPEG, molar mass ~ 500 g/mol) was dissolved in 3 mL of ethanol and dropwise added to silica-coated UCNPs in ethanol (1 mL) while stirring. For biotin-conjugated UCNPs, silane-PEG(3.4k)-biotin (4 mg) was dissolved in 500 μL of ethanol, and added to the mixture while stirring. Subsequently, 200 μL of water and 10 μL of ammonia (28%) were added to the mixture. The solution was heated to 50 $°C$ and stirred overnight. The PEGylated UCNPs were then cooled to room temperature. The solution was diluted to 10 mL in ethanol and collected by centrifugation (16,900 g, 20 min). The UCNPs were washed twice in ethanol and twice in potassium phosphate buffer (PBS, pH 7.4) to remove excess reagents. The particles were finally filtered through a 0.22 μm filter and stored in PBS buffer at 4 $°C$.

Binding assay of biotin-UCNPs and streptavidin-QDs

To test the specific binding of biotin-conjugated UCNPs (btn-UCNPs), 5 μL of 40 pM btn-UCNPs or UCNP@$SiO_2$-mPEG was sonicated and incubated with 10 μL of 20 nM streptavidin coated QD705. After 5 hours of incubation, the mixture was diluted 10x and drop cast on TEM grids for TEM imaging.

Single particle binding assay on biotinylated PEG coverslips

To test the specific binding of biotin-conjugated UCNPs (btn-UCNPs), we also performed single particle binding assay on PEGylated coverslips. PEGylated coverslips and sample chamber were prepared as described previously (*4*). Ten microliters of NeutrAvidin in PBS (1 μM) was added to the sample chamber and incubated for 5 min. Unbound NeutrAvidin was washed thoroughly with 100-200 μL of PBS buffer. Btn-UCNPs (30 pM) was injected in the sample chamber and incubated for 5 min before being washed thoroughly. For control experiments, NeutrAvidn was not added before btn-UCNPs incubation.

Cell surface binding assay

To examine the specific binding and nonspecific sticking of btn-UCNPs on cell surface. We biotinylated the cell surface of Hela cells and performed binding experiments similar to what was done on PEGylated coverslips. Hela cells were plated in Lab-Tek 8-well chamber at a density of 30,000 cells per well, and cultured with DMEM (high glucose), 10% fetal bovine serum (FBS), 2 mM L-glutamine, 100 U/mL penicillin-streptomycin. Cells were grown to 95% confluent before the experiment. The stock biotinylation kit containing biotin-XX sulfosuccinimidyl ester (0.2 mg/mL in DMSO) was diluted to 5 μg/mL in culture medium. The culture medium in the Lab-Tek well was aspirated and 150 μL of 5 μg/mL biotinlyation reagent was added. The cells were incubated for 15 min at room temperature. Subsequently, the cells were washed with cold PBS three times to remove unreacted reagent. One hundred microliter of NeutrAvidin in PBS (1 μM) was added to the cells and incubate for 10 min at room temperature. Unbound NeutrAvidin was washed thoroughly with cold PBS five times. One hundred microliter of btn-UCNPs (100 pM) was then added to the cells and incubate for 5 min at room temperature. Unbound UCNPs were washed thoroughly with cold PBS five times. For control experiments, NeutrAvidn was not added.

Rat DRG neuron culture

DRG neurons were dissected out of Sprague-Dawley rat embryos at embryonic day 18 (E18) in Hanks' buffered saline solution. The DRG cells were then treated with 5 mL of 0.25% trypsin for 30 min at 37°C followed by mechanical trituration. Equal volume of Dulbecco's modified Eagle's medium (+ 10% fetal bovine serum) was added to quench trypsin and the dissociated cells were spun down with 200 rpm for 8 min. Cells were resuspended in DRG maintenance media (neurobasal, B27 plus, 2 mM L-glutamin/GlutaMAX, 100 U/mL penicillin-streptomycin, and 50 ng/mL nerve growth factor). After counting the cell density, cells were spun down and resuspended in appropriate volume of maintenance media to reach final concentration of 10 million cells/mL. Ten microliters of cells were plated in the soma compartment of the microfluidic device. Microfluidic device (SND900 from Xona Microfluidics) was pre-assembled on 24 mm x 60 mm № 1.5 coverslips pre-coated with 0.5 mg/mL poly-D-lysine overnight. Cells were maintained at 37 $°C$ with 5 % $CO_2$ for up to 21 days. On 2 days-*in vitro* (DIV 2), the media was replaced with antimitotic media (maintenance media containing 1 μM cytosine arabinoside) in order to suppress glial cell proliferation. The media was replaced back to normal maintenance media on DIV 3. Rats used in this study were treated in accordance with Stanford University's institutional guidelines.

Human induced neuron

Human induced neuronal (iN) cells were generated from trans-differentiating human embryonic stem cells (WA01/H1 cell line, NIH registry 0043) as described in Huang et al (5). In brief, H1 cells were plated on matrigel-coated surface and transduced lentivirally to express the single transcription factor Neurogenin 2 (Ngn2) and the selection marker puromycin N-acetyl transferase (PuroR) in a tetracycline-inducible fashion, driven by tetracycline operator tetO promoter plus the co-expression of transactivator (tTA). After lentiviral infection (day -1), doxycycline (2 mg/L) was added on day 0 for three days, and puromycin (1 mg/L) selection was performed on day 1 and day 2. On day 3, differentiating human iN cells were detached, plated on one compartment of a matrigel coated microfluidic chamber (10 μL of 10 million cells/mL.), and maintained in Neurobasal A medium with B27 supplement.

UCNPs internalization in neurons

For both rat DRG and human iNs, live cell imaging was done between DIV 14 to DIV 21. Media in the axon or soma chambers (for retrograde or anterograde transport, respectively) were removed and 10 μL of 1 μM of biotin-WGA (Vector Laboratories B-1025) was added. After 5 min incubation, 150 μL of maintenance media was used to wash away free biotin-WGA. Subsequently, 10 μL of 1 μM of streptavidin was added for 3 min. Excess streptavidin was removed by washing with 150 μL of maintenance media twice. After washing, 1 nM of Biotin-UCNPs (diluted in maintenance media) was added and the cells were incubated at 37 $°C$ with 5 % $CO_2$ for 2 hours. Before imaging, free UCNPs were washed away with 150 μL of maintenance media

Live neuron imaging

Imaging was performed using a home-built microscope with wide-field epi-illumination of a 976 nm fiber laser through a 100X oil-immersion objective (NA=1.49, Nikon). A microscope stage-top incubator (STRF-WELSX-SET, Tokai Hit) was used for live cell imaging at 37 $°C$ and 5% $CO_2$. The upconverting luminescence signal was recorded on an EMCCD camera (iXon 897, Andor). To achieve sufficient spatial resolution at 10 ms exposure time for fluctuation theory analysis, live neuron imaging was performed at 6.5 kW/cm$^2$. However, 976 nm excitation with much lower power density could be used to image single cargoes at longer exposure times.

UCNP saturation curve measurement

Procedures for measuring UCNP saturation curves have been described previously (*1*). Briefly, approximately 400 ng/mL UCNPs in cyclohexane was drop cast onto a clean and dry № 1.5 cover glass pre-coated with 1% (w/v) poly-L-lysine. Cyclohexane was used to rinse off excess nanoparticles after 1 min incubation. For rigid support, the cover glass was attached to a standard microscope slide using double-sided tape. Custom IDL code was used to identify individual point spread function (PSF) and perform 2D Gaussian fit to determine the upconversion emission rate. In order to correct measured UCNP luminescence by the wide-field illumination profile, the microscope stage was scanned across the field of view (FOV) with equal step size of 1 μm. The position-dependent luminescence profile of a single UCNP was used to compute the wide-field illumination profile.

Correlative SEM and wide-field fluorescence imaging

Nanoparticles were drop cast as described above onto a glass coverslip with an alphanumerically labeled grid pattern marked in 50 μm increments (IBIDI grid-50, IBIDI, Germany). The sample was first characterized under wide-field optical illumination. Subsequently, a thin layer of 2 nm gold-palladium was sputter-coated (Denton Vacuum, USA) onto the same sample to enhance conductivity, and nanoparticles were imaged using a Zeiss Sigma Field Emission Scanning Electron Microscope (Carl Zeiss Microscopy, Germany) and InLens SE (Secondary Electron) detection. The fine grid pattern served as a navigation guide to locate the FOVs which had previously been optically characterized. Once registration was established between geometric patterns of the fluorescent image and electron micrograph, we then zoomed in to verify the oligomeric state and the size of each individual nanoparticles.

Single particle tracking

For single particle tracking, individual PSFs were localized and their time-trajectories were resolved using custom MATLAB scripts which performed 2D Gaussian fitting with multiple-target tracing (MTT) method (*6*, *7*).

Hidden Markov Model (HMM)-Bayes analysis

Single cargo trajectories were analyzed with HMM-Bayes, which infers stochastic switching between diffusive and directed transport states from measured particle displacement (*8*). The HMM-Bayes software written in MATLAB was downloaded from http://hmm-bayes.org/ and used with default parameters. The program automatically annotates when and where each type of motion occurs in space and time along the trajectory with single-step resolution.

Dynein step size determination

To determine the molecular step size in live DRG neurons, retrograde transport was imaged at 2.5 msec exposure time at 22 $°C$ and 30 $°C$, and at 1 msec frame rate (0.8 msec integration time and 0.2 msec data transfer time) at 37 $°C$. Single particle trajectories were obtained as described above. Step sizes were determined by a step-finding program developed by Kerssemakers et al (*9*). Step sizes smaller than the experimental noise of 4 nm were excluded.

### Supplementary Notes

#### Note S1. Analysis of evidence in support of the active cycling model

There are several arguments in the literature used to support of the active-cycling model:

(i) A bulk assay was used to measure the rate of $Pi$ generation due to ATP hydrolysis per dynein stepping under saturating concentrations of microtubules binding to dynein (*10*). The authors argue that an ATPase rate of $k_{cat}$~16 $Pi$ per second per dimer, if the step is ~ 8 $nm$ steps with a 0.8 probability of moving forward and a velocity of $100\ nm \cdot s^{-1}$ equates to 1.026 $Pi$ per step (*10*).

However, using their reported mean velocity of $85 \pm 30\ nm \cdot s^{-1}$, the number of $Pi's$ generated per nm is 0.188 (*10*). In a subsequent publication by the same group, $20\ Pi \cdot s^{-1}$ at a velocity of $69 \pm 22\ nm \cdot s^{-1}$ gives 0.29 $Pi$ per nm (*11*). The step-size histogram distribution of the tail-labeled dynein is ~ 5 $nm/step$ based on both our data (Fig. 4C) and the earlier *in vitro* data (*10, 12*). Our assignment of 5 $nm/step$ *explicitly includes* the probability of not moving forward or backward during a ATP hydrolysis dynein cycle. If the step size of a stepping monomer is assumed to be twice the step size of the dimer tail, as was done in Ref. (*10*), then the average value of the number of $Pi$ per step = 2.39. Thus, we find that the data of (*10, 12*) full stepping cycle uses =2 $Pi/step$ if ~ 84 % of the dynein were biologically active in their $Pi$ assay.

(ii) A Michaelis-Menten (MM) analysis of the dynein activity (velocity dependence of motility vs. $[ATP]$) agrees with a kinetic rate Hill coefficient of $n = 1$ that is expected if the activity is due to the cycling of a single $ATP$ (*13*). A fit to $n = 2$ (and a sigmoidal dependence of the enzyme kinetics on $[ATP]$ requires that there must be two $ATP$ binding sites. However, it has been shown that if the enzymatic cycle contains steps separated by effectively irreversible transitions (such as the hydrolysis of $ATP$ in live cells), the MM reaction rate can also be fit with a Hill coefficient $n = 1$, even if there are two or more $ATP$ binding sites (*14*). In Fig. S30, we explicitly show that if a single dynein step involves sequential hydrolysis of two ATPs, the observed $[ATP]$-dependent velocities reported in Ref. (*13*) satisfies the Michaelis-Menten equation with a Hill coefficient of $n = 1$. While we also note that another earlier velocity dependence was described by to two different Michaelis-Menten parameters, which suggests two $ATP$ binding sites (*15*), the primary conclusion is that Hill coefficient analysis is inconclusive.

(iii) In a series of experiments that analyzed transient pauses in motility induced by adding ATPγS, a slowly hydrolyzing ATP analog in wild-type (WT) and mutant forms of dynein (*13*), the authors concluded that AAA3 domain acts as a stepping gate. Their WT homodimer motility in the presence of ATPγS fits to MM Hill coefficient of $n = 2$. In the active-cycling model, each monomer contains one binding site, and *if* both dynein monomers are required to be active for motility, $n = 2$ would be expected. The group also studied the activity of an SRS-WT heterodimer, where movement can only be generated by the dynein monomer, and their data was consistent with $n = 1$. While suggestive, the motility of the mutant construct was compromised and there is no guarantee that the enzymatic activity reflects dynamics of the WT-homodimer. Furthermore, the data in Ref. (*13*) as pointed out above in (ii) does not rule out a model that demands the hydrolysis of 2 $ATP$'s on the active dynein monomer.

**Note S2. The quantification of the diffusion range of a dynein monomer as a function of temperature**

The diffusion $\langle x^2(t) \rangle$ in Fig. 5D is temperature dependent because the time a dynein monomer spends in State II of Fig. 5 of the Main Text is determined by the rate of $Pi$ release from AAA3. In Fig. 5C, we show that the desorption rate constants $k \equiv k_1 = k_2$ at 22 °$C$, 30 °$C$, and 37 °$C$ fit to an Arrhenius plot with a thermal activation energy equal to 25.1 kcal/mol. From Fig. 5B, the times spent in weakly bound State II are $t = 1/k_1 = 71.9\ ms$, $17.8\ ms$, and $10.3\ ms$.

The exploration of the next binding site is made during the time $ADP \cdot Pi$ is bound to both AAA1 and AAA3 of the stepping monomer, as shown in State III of Fig. 6. While in this state, the MTBD oscillates between being weakly bound to the MT, and in an unbound state when it is free to explore a new stepping position on the MT. To estimate the times $t_{bnd}$ and $t_{unbnd}$ spent in the bound and unbound states, we use the kinetic rate constants for $D \cdot MT \xrightarrow[k_{off}^{pre}]{} D + MT$ and $D + MT \xrightarrow[k_{on}^{pre}]{} D \cdot MT$ measured by Kinoshita, *et al.* (*16*). Although their analysis assumes that dynein requires the hydrolysis of a single $ATP$ per stepping cycle, the rate constants are related by the equilibrium constant $K_d = k_{pre}^{on} / k_{pre}^{off} \sim 34\ \mu M$, which was determined from the co-sedimentation measurements of dynein bound to a MT in $200\ \mu M\ ADP \cdot Pi$, and should still be applicable. The rates $k_{pre}^{off} = 34\ s^{-1}$ and $k_{pre}^{on} = 1\ \mu M^{-1} s^{-1}$ were measured at room temperature (*17*). The average time spent in the bound state is $t_{bnd} = 1/k_{pre}^{off} = 29\ ms$.

In order to determine the time $t_{unbnd}$ spent in the freely diffusing state, the effective microtubule concentration $[MT]$ has to be estimated. This is done by determining the search volume $\delta V = \delta x \delta y \delta z$ explored by the MTBD of the dynein monomer. The step-size histograms at $22°C$ is fit to a Gaussian curve (Fig. S25A-C) where longitudinal diffusion is given by the Gaussian width $\sigma$. If we define the distance $\delta x$ over which the area under the Gaussian curve is 50% of the total area,

$$\int_{\mu-\delta x/2}^{\mu+\delta x/2} e^{-(x-\mu)^2/2\sigma^2} dx = \frac{1}{2} \int_{-\infty}^{+\infty} e^{-(x-\mu)^2/2\sigma^2} dx, \tag{S1}$$

$\delta x = 1.35\sigma = 1.35 \times 2 \times 8.5\ nm$ at $22°C$. The $\sigma = 8.5\ nm$ distance is from the fit in Fig. S25E and the factor of 2 is because the exploration distance of a monomer is twice the distance of the recorded movement of vesicle, as discussed in Fig. S23. We assign $\sigma_y$ to be $6.5\ nm$ based on the sideway diffusion measurement (*18*). The vertical diffusion $\sigma_z$ is less certain because of the constraints that may be imposed by the p150 domain of dynactin, accessary molecules such as Lis, and/or the transient salt bridges between the MTBD and the MT (*19*). Since the MTBD may be "sliding" in the near proximity of the MT, we estimate that $\delta z \sim 1.5 \pm 1.0\ nm$. Therefore, the search volume $\delta V = \delta x \delta y \delta z = (1.35 \times 2 \times 8.5\ nm)(1.35 \times 6.5\ nm)(1.5 \pm 1.0\ nm) \sim 3.02 \times 10^{-25}\ m^3$. This is the search volume of one MTBD in the vicinity of a MT. Alternatively, we can use this volume occupied by a single MT to calculate an effective concentration of microtubules to be $[MT] = 5.5\ mM$. Thus,

$$t_{unbound} = \frac{1}{k_{pre}^{on} \times [MT]} = \frac{1}{(1\ \mu M^{-1} s^{-1}) \times 5.5 mM} = 182\ \mu s.$$

The number of cycles of transient binding and unbinding before $Pi$ is released from AAA3 is

$$\#\ of\ cycles = \frac{72\ ms}{29\ ms + t_{unbound}} = \frac{72\ ms}{29.182\ ms} = 2.47\ cycles,$$

and the total time available to exploratory diffusion is $\Delta t = 182\mu s \times 2.47 = 0.45\ ms$.

Next, the solution to the elastic-spring confined diffusion

$$\langle x^2(t) \rangle = \frac{2k_B T_{avg}}{\kappa}\left(1 - e^{-\Delta t \kappa/\gamma}\right) \tag{S2}$$

is used to infer a value for the viscous drag coefficient $\gamma$ in $F_d = v = 6\pi\eta a_{eff} v$. In Fig. S25 we used the fitted parameter $\sigma_\infty^2$ to calculate $\kappa = 1.1 \times 10^{-4}\ J/m^2$ for the vesicle diffusion. In our model, we assume that the stepping dynein monomer diffuses twice the distance of the net motion of the vesicle. Hence, $\sigma_{\infty,monomer}^2 = 2\sigma_\infty^2$ and $\kappa_{monnomer} = 2.75 \times 10^{-5}\ J/m^2$. The viscous drag of a dynein monomer is estimated from the crystal structures of the dynein monomer AAA ring, stalk and MTBD to be $\gamma = 6\pi\eta a_{eff} = 6\pi\eta(6nm)$. From the inferred value of $r_{rms} = \sqrt{\langle x^2(t) \rangle} = 17\ nm$, the axonal viscosity is estimated to be $\eta_{monomer} = 0.029\ Pa \cdot s$.

The viscosity of an object in a cell is size-dependent due to molecular crowding. For an object of diameter $d \sim 100\ nm$, the viscosity is in the range of $\eta_{cell} \sim 1 - 3\ Pa \cdot s$ (*20*), and for objects the size of organic dye molecules, $\eta_{cell}$ is close to that of $\eta_{H_2O} \sim 10^{-3} Pa \cdot s$. The estimated effective viscosity of an object the size of a dynein monomer is less certain. Mika, et al. reported $\eta \sim 0.056\ Pa \cdot s$ in a bacterial cell *L. lactis* (*21*), but eukaryotes are expected to have higher viscosity. Other estimates of cellular viscosity from $\eta \sim 0.005 - 0.05\ Pa \cdot s$ (*22*). Nanoscale viscosity measurement in HeLa cells give $\eta_{eff} \sim 0.006\ Pa \cdot s$ (*23*). In the confined geometry of axons, the increase in viscosity due to crowding and cell wall proximity could increase the axonal viscosity relative to cellular viscosities (*24*). In addition, as pointed out above, there may be additional sliding friction due to weak MTBD-MT interactions.

**Note S3. Estimates of the diffusion time of the dynein while in the weakly-bound state at higher cell temperatures**

Since there are no kinetic measurements at 30°C and 37°C, we assume the decrease in $\sigma$ at higher temperatures is due to the shortened diffusion time of the dynein monomer. With this assumption, the time in the freely diffusing state at 30°C and 37°C were calculated to be 250 $\mu s$ and 116 $\mu s$ respectively (Table S2). Direct kinetic measurements of the binding times of dynein aligned properly with the microtubule as in Ref. (*25*) could test the kinetic predictions of our microscopic model and spring constants of a dynein dimer $\kappa_{flex}$ under saturating conditions of $ADP$.

**Note S4. Estimate of the instantaneous viscous drag force $F_d = 6\pi\eta_{eff} a v$ due to the cargo vesicle during the power stroke**

The Stokes drag force is given by $F_d = 6\pi\eta_{eff} a v$, where $\eta_{eff}$ is the *effective* drag viscosity in the axon experienced by the vesicle of radius $a$ is the *effective* drag viscosity in the axon experienced by the vesicle of radius $a$ moving with velocity $v$ is the velocity of the system. Measurements of $\eta_{eff}$ using ~ 100 nm diameter particles (*20*) are in the range of $0.8 - 3.0\ Pa \cdot s$.

$$\tau_c = 23\ ms = \frac{\gamma}{\kappa} = \frac{6\pi\eta_{cargo}a_{cargo}}{\kappa_{cargo}} = \frac{6\pi\eta_{cargo}(100\ nm)}{1.1\times10^{-4}\ J/m^2} \rightarrow \eta_{cargo} = 1.34\ Pa\cdot s$$

For average velocity of 1 $\mu m/s$, the average drag force is $2.5\ pN$. The concatenation of the transition times (Fig. S25) is less that $1.0\ ms$, we may safely assume the duration of the power stroke $\Delta t_{P.S.} < 0.5\ ms$. For an 8 $nm$ step, the instantaneous velocity $v_{instant} > \frac{8\ nm}{0.5\ ms} = 16\ \mu m/s$, therefore, the instantaneous force exerted by the dynein during the power stroke is $F_{P.S.} > 40\ pN$. We stress that this estimate of the instantaneous force during the power stroke is a lower limit, and it is conceivable that $F_{P.S.} \geq 80\ pN$.

### Supporting information figures

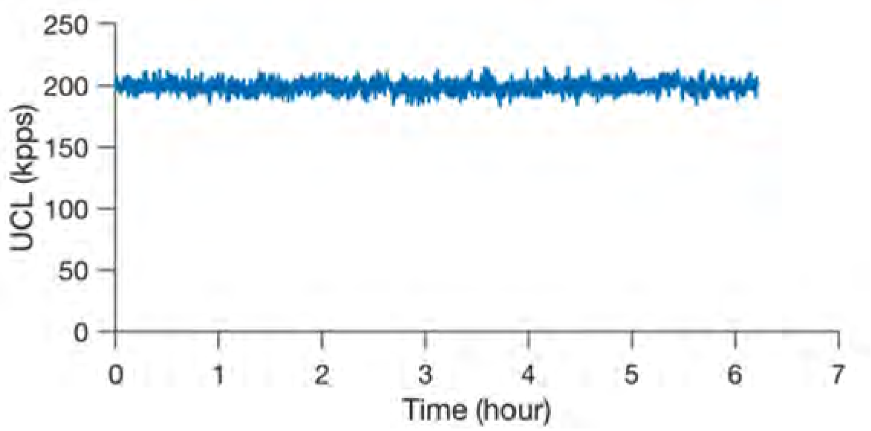


**Fig. S1.** Long-term stability of upconversion luminescence of single CSS nanoparticles, $NaYF_4$@ $NaYbF_4$: 8 % Er@$NaYF_4$ with 27 kW/cm$^2$ of 978 nm excitation. For intensities up to 6 $MWcm^{-2}$, we observed no evidence of photobleaching

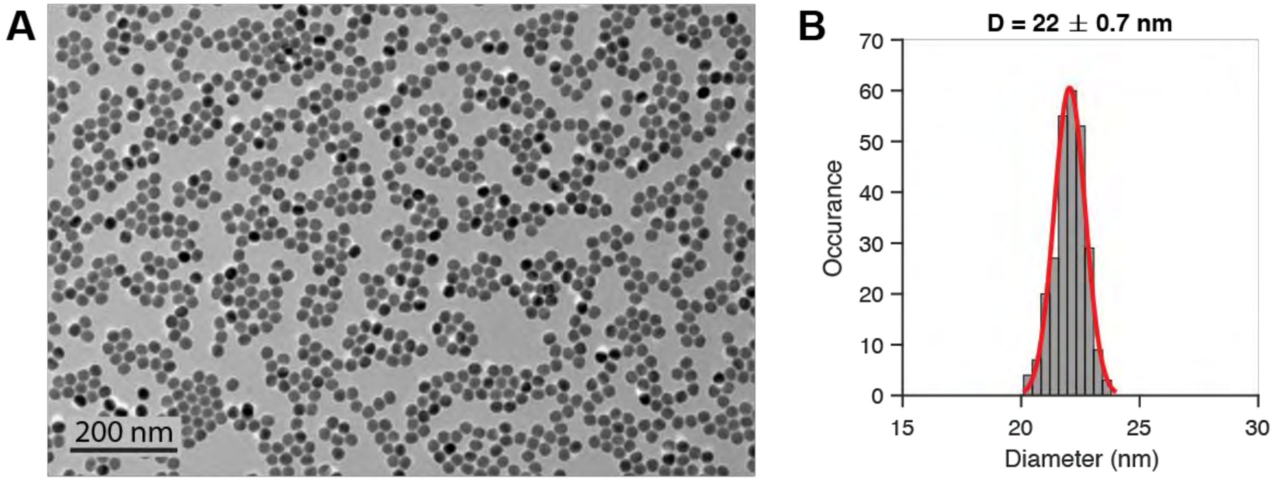


**Fig. S2.** TEM image (A) and size distribution (B) of the core-only $NaYF_4$: 20%Yb, 2% Er. Synthesis and detailed characterizations of this UCNP has been reported in our previous publication (*1*).

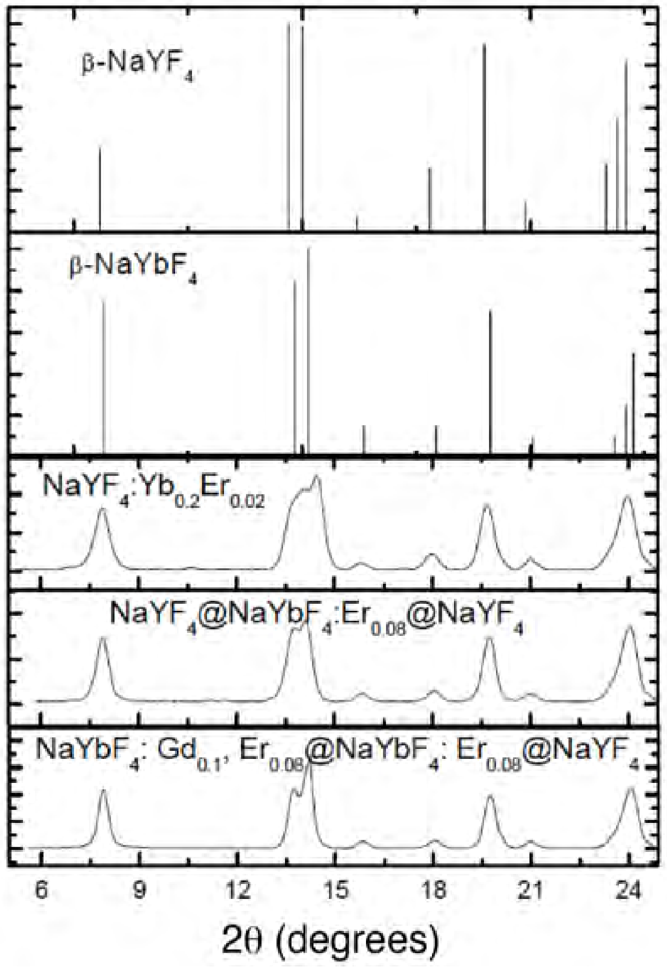


**Fig. S3.** XRD patterns of the UCNPs and standard β-phase $NaYF_4$ and $NaYbF_4$.

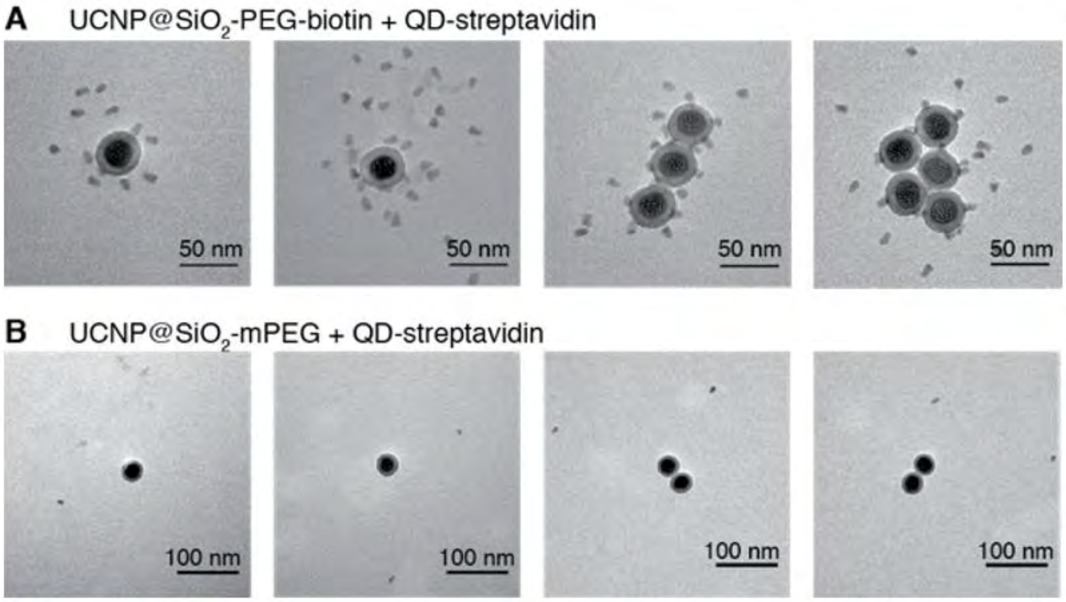


**Fig. S4.** TEM images of mixture of streptavidin coated quantum dots with (A) $NaYF_4$: 20% Yb, 2 % Er@$SiO_2$-PEG-biotin and (B) $NaYF_4$: 20% Yb, 2 % Er@$SiO_2$-mPEG. Larger field of views were displayed in (B) in order to show the presence of unbound QD-streptavidin, demonstrating that the bound QDs in (A) are due to specific biotin-streptavidin interactions rather than non-specific sticking.

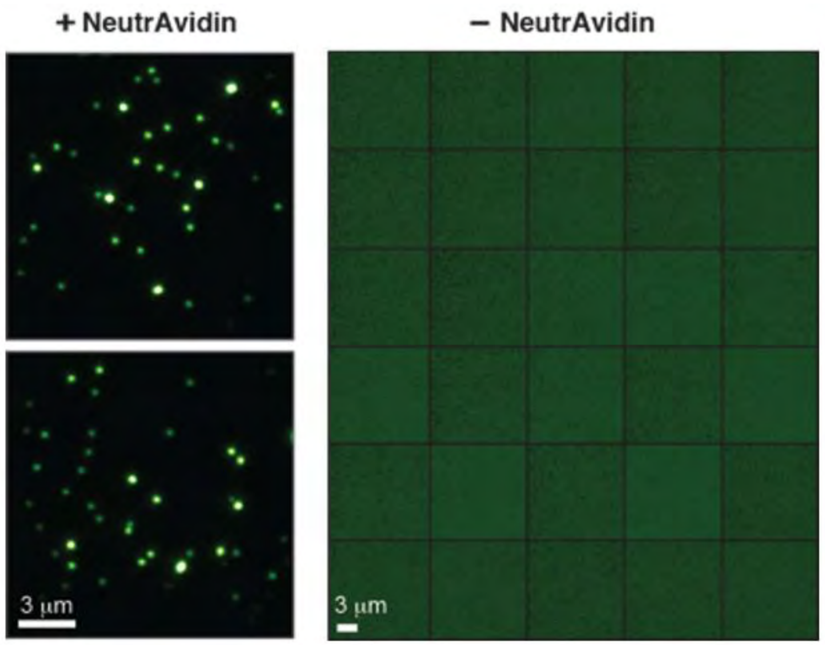


**Fig. S5.** Wide-field fluorescent images of $NaYF_4$: 20% Yb, 2 % Er@$SiO_2$-PEG-biotin on biotinylated PEG coverslips in the presence (left) and absence (right) of neutravidin. When btn-UCNPs were incubated on coverslips for 5 minutes without neutravidin and subsequently washed away, no particles were detected in at least thirty field of views. Under background-free imaging condition (976 nm excitation) and the absence of UCNPs, it was necessary to check the focus was maintained during image acquisition. For every five images taken, we switched to 532 nm excitation and used the autofluorescence from the coverslip surface to re-adjust the focus. The fact that no UCNPs were detected in 30 FOVs while on average 65 UCNPs were found in the presence of NeutrAvidin allowed us to establish at least 2000:1 specific versus non-specific binding of btn-UCNPs.

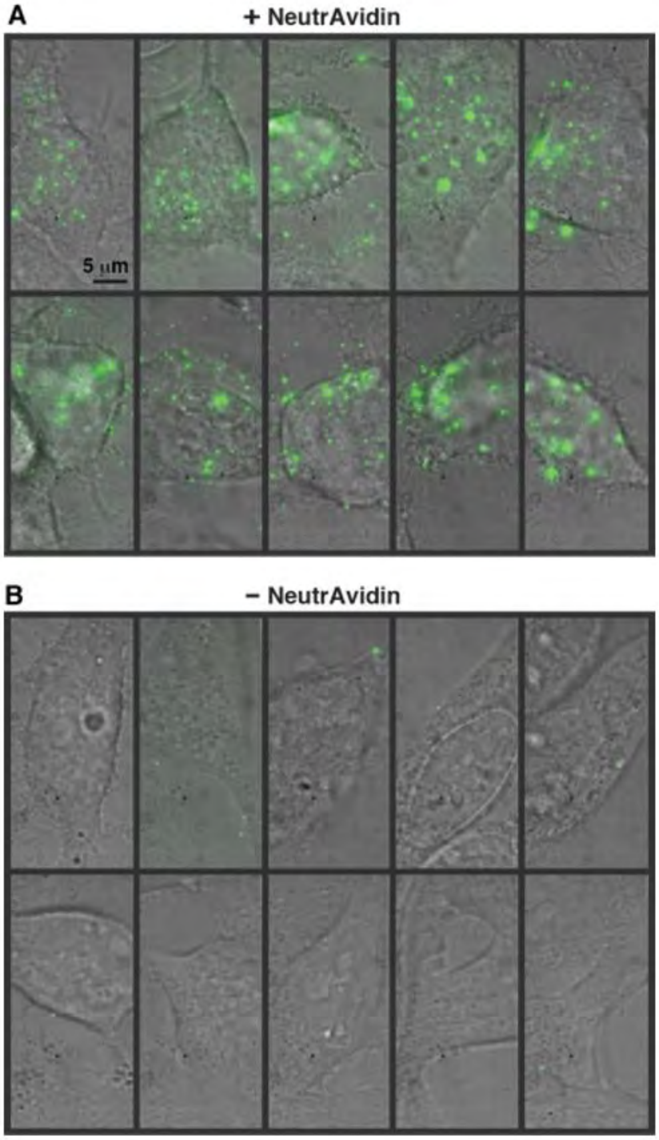


**Fig. S6.** Other examples of live-cell imaging of btn-UCNPs on HeLa cells whose membrane proteins have been biotinylated. Overlay of brightfield images of HeLa cells and maximum-intensity-projected luminescence images of biotinylated-UCNPs (btn-UCNPs) in the presence (A) and absence (B) of NeutrAvidin.

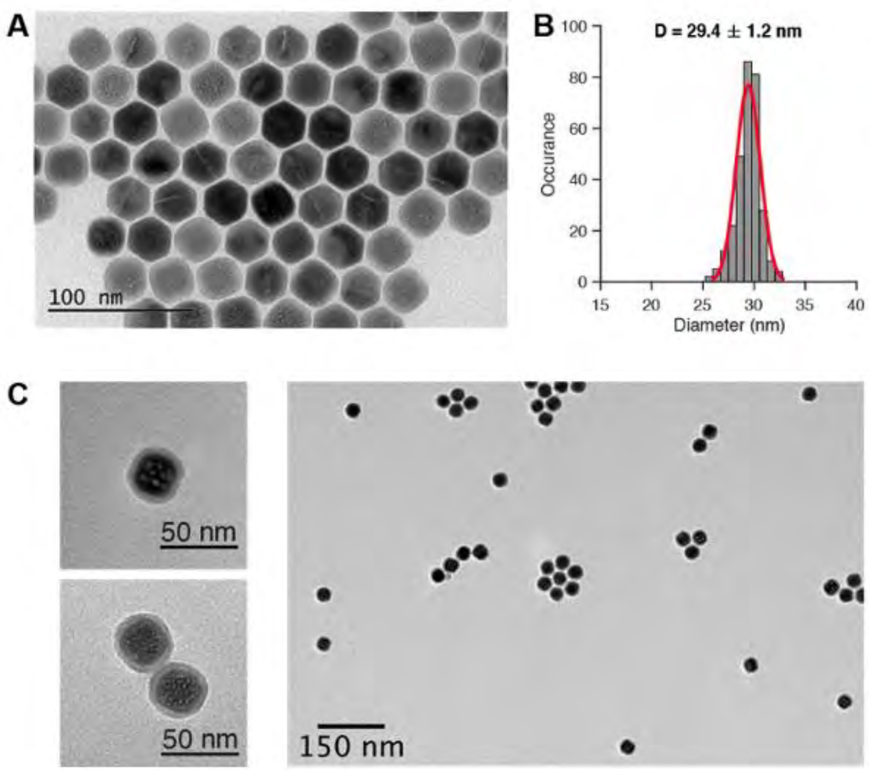


**Fig. S7.** TEM image (A) and size distribution (B) of the core-shell-shell $NaYF_4$@ $NaYbF_4$:8% Er@$NaYF_4$. Synthesis and detailed characterizations of this UCNP has been reported in our previous publication (*1*). (C) TEM images of silica coated UCNPs. The silica shell thickness is ~ 4 nm.

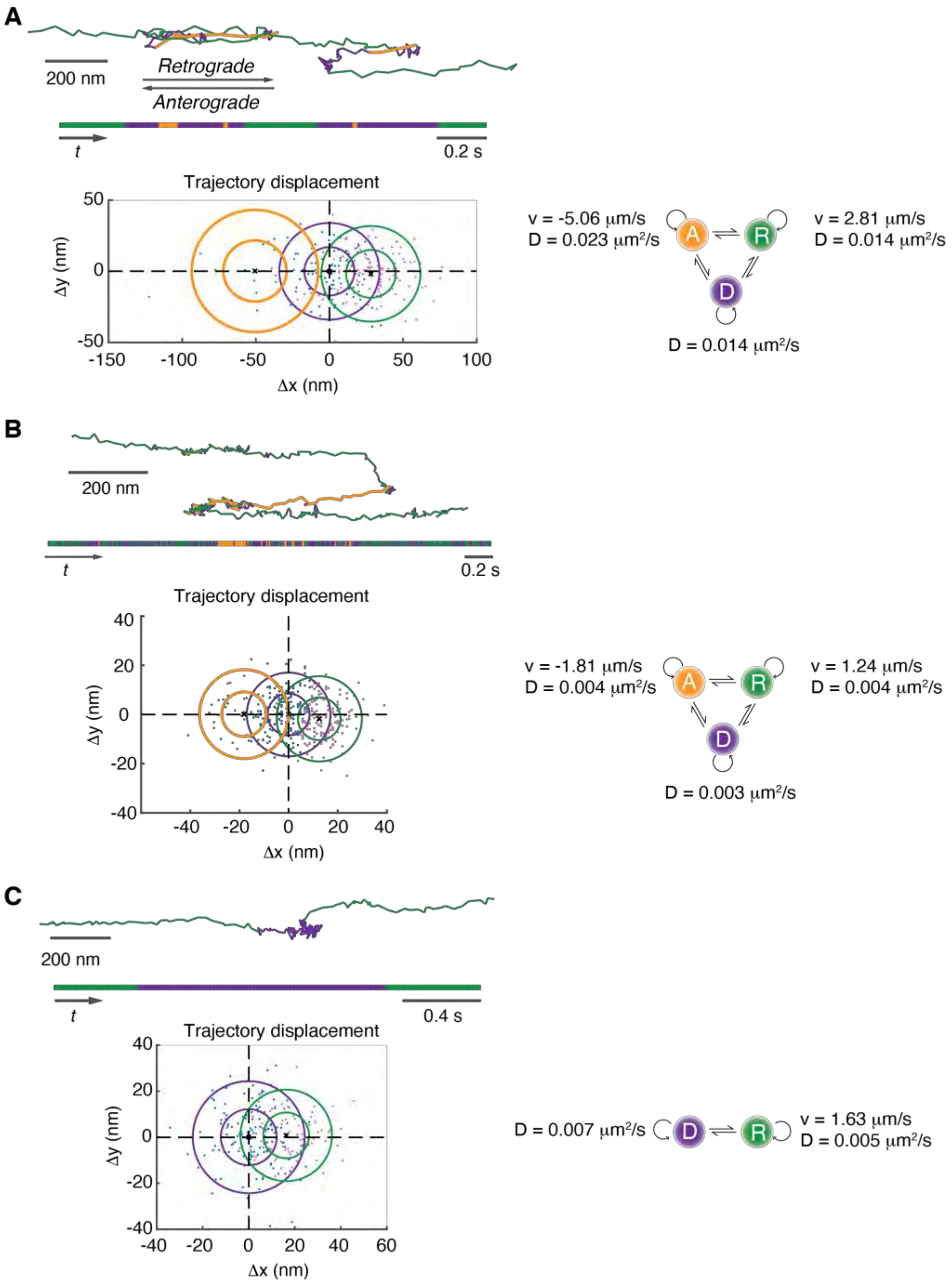


**Fig. S8.** HMM-Bayes analysis of the retrograde transport in rat DRG (A) and human iNs (B), corresponding to the trajectories shown in Fig. 2B and 2C. For each trajectory, scatterplots of the displacements are shown on the bottom and each point is colored blue, pink, and green indicating the inferred diffusive, retrograde or anterograde transport, respectively. Circles indicate one and two standard deviations of the displacements within each state. The inferred diffusion coefficients and velocities are shown on the right. (C) One example of iN trajectory that shows only retrograde and diffusion states.

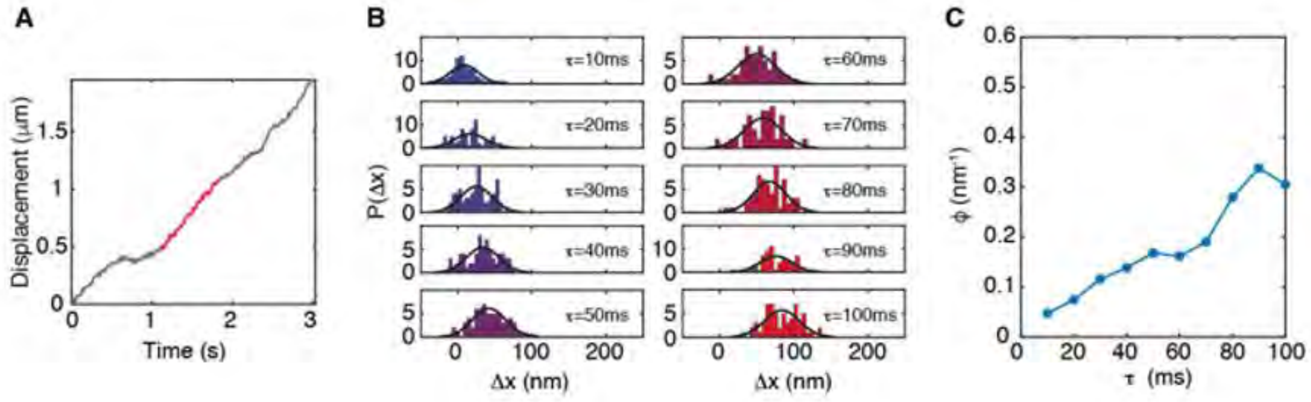


**Fig. S9.** One example of constant velocity segment that did not show clear where $\phi$ does not reach a steady-state asymptotic value. (A) Displacement curve of retrograde transport in rat DRG neurons. (B) Probability distribution of displacement $\Delta x$ at different time delays (10 ms to 100 ms). (C) Relaxation curve of the effective entropy $\phi$, calculated from the mean and variance of the Gaussian distributed $P(\Delta x)$.

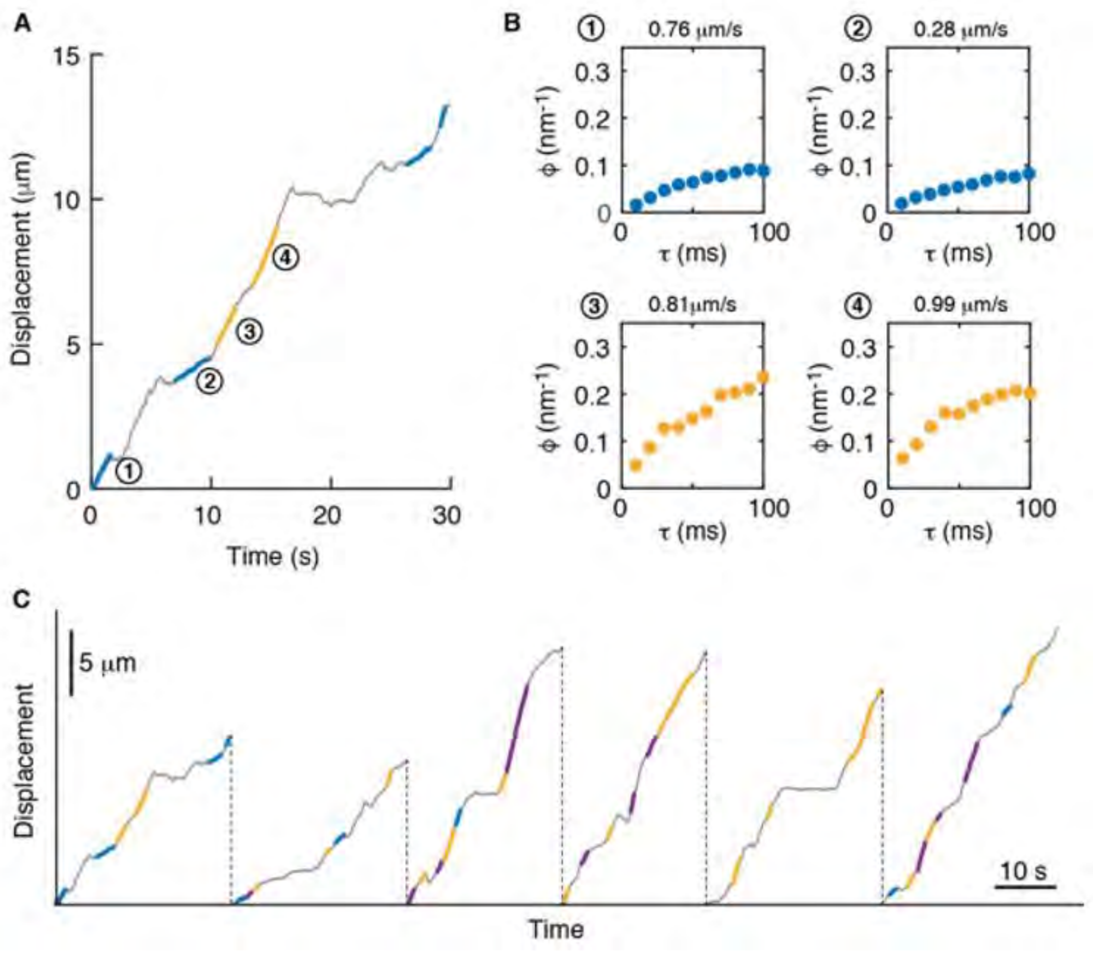


**Fig. S10.** Example showing switches between different $\phi$ values during the retrograde transport. (A) Displacement curve of retrograde transport in rat DRG neurons. Colored lines mark the constant velocity segments that show clear relaxation of $\phi$ curves. (B) Relaxation curves of the effective entropy calculated for each of the four constant velocity segments marked in (A). The average velocities of these segments are shown on the top of each panel. (C) More displacement curves from the same cargo. Color codes follow Fig. 3F where blue, yellow, and purple indicate one, two, and three pairs of dynein, respectively.

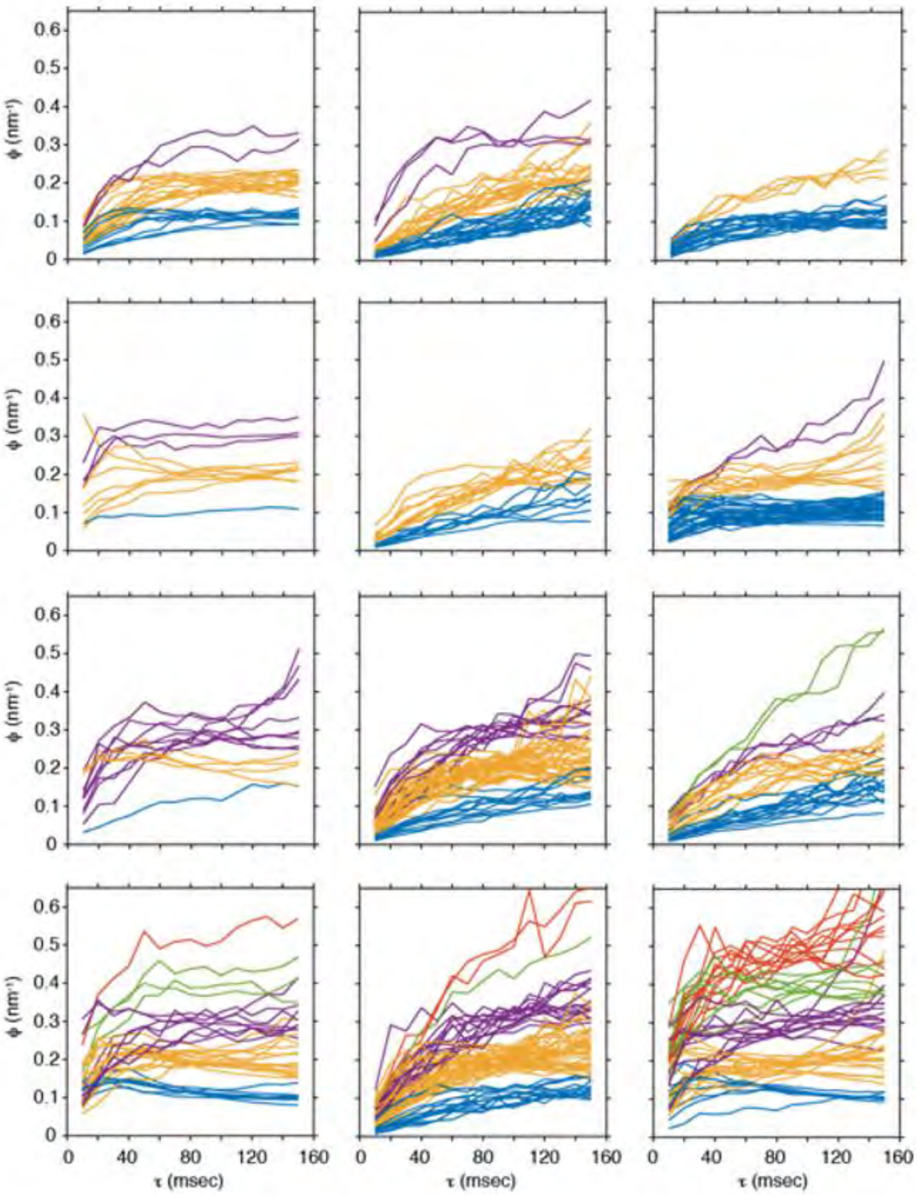


**Fig. S11.** Effective entropy $\phi$ curves for twelve individual retrograde endosomes in DRG neurons. In addition to the example shown in the main text which exhibits quantized $\phi$ values of 0.1, 0.2, and 0.3, some endosomes only show $\phi$ values of 0.1 and 0.2, while some endosomes have $\phi$ values up to 0.5. The number of $\phi$ curves for each endosome depends on the length of the recorded trajectories.

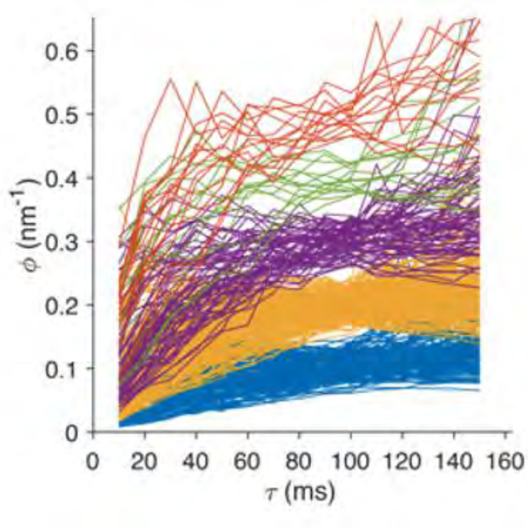


**Fig. S12.** Pooled effective entropy $\phi$ curves from 12 different retrograde cargoes in rat DRG neurons.

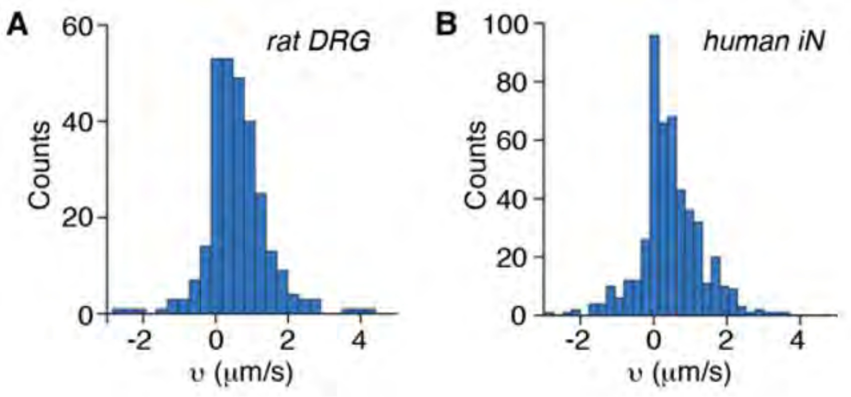


**Fig. S13**. Velocity distribution of single retrogradely transported cargo in (A) rat DRG and (B) human induced neurons. No clear quantized peaks were shown by these velocity distributions.

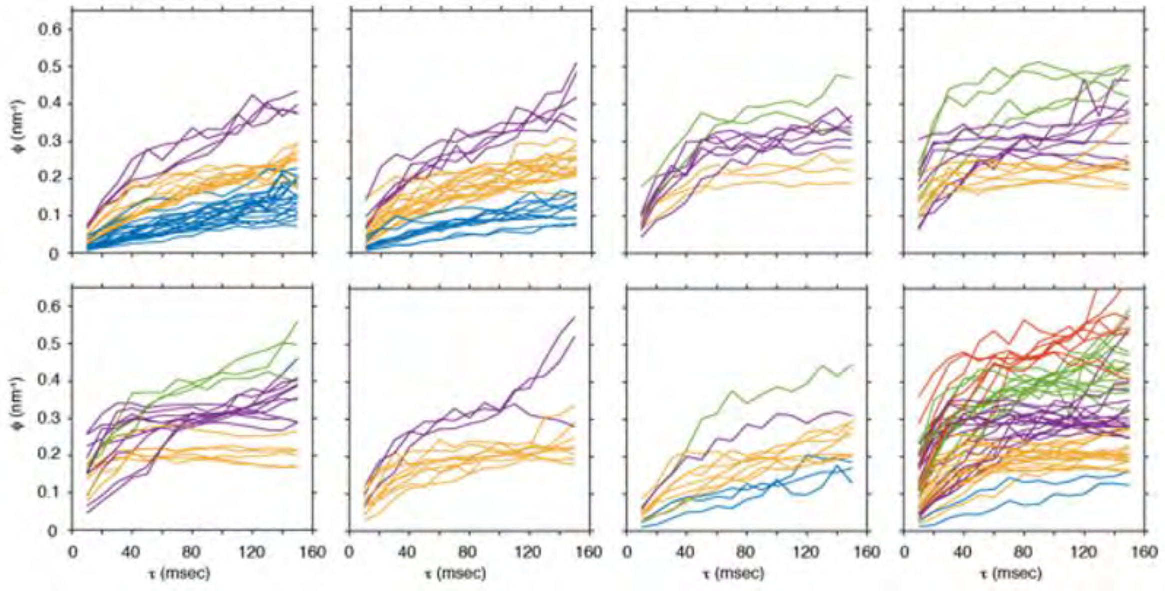


**Fig. S14.** Effective entropy $\phi$ curves for eight individual retrograde endosomes in human induced neuron.

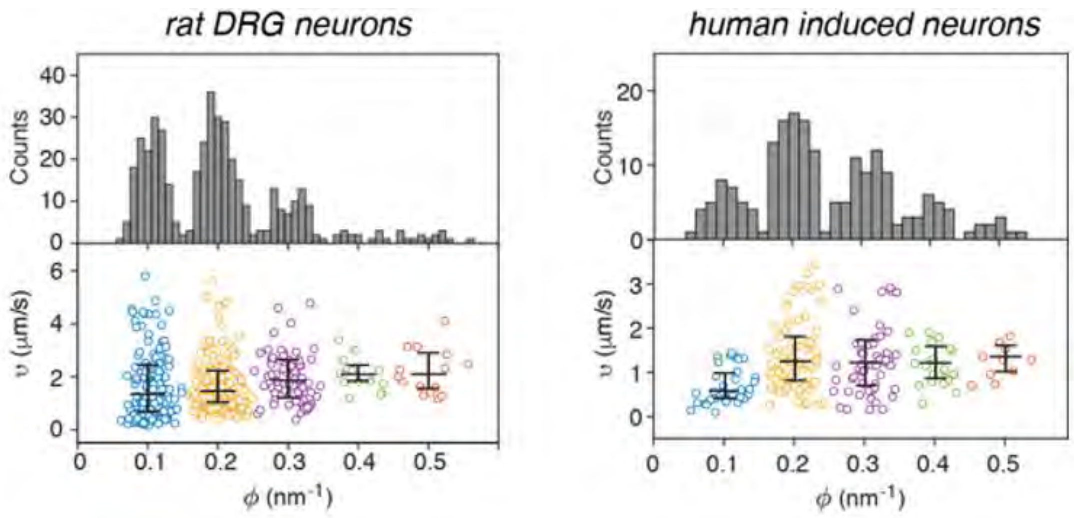


**Fig. S15.** Fluctuation theorem analysis results for rat DRG neurons and human induced neurons. The $\phi$ was averaged from $\tau$ = 80 msec to 120 msec. N = 430 and 194 for rat DRG and human iN, respectively. The average (± S.E.M.) pairs of dynein were 2.01 ± 0.05 and 2.50 ± 0.08 for rat DRG and human iNs, respectively.

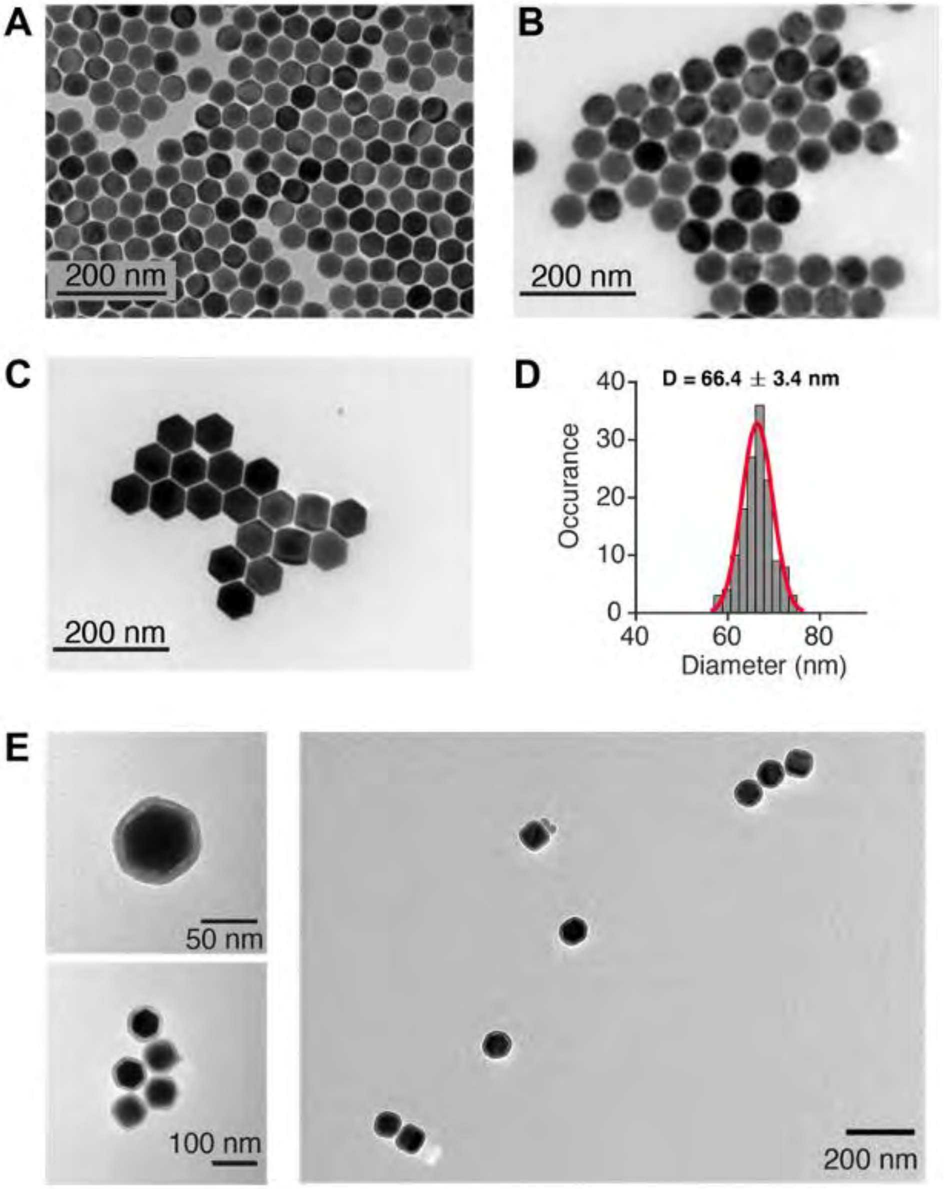


**Fig. S16.** TEM images of (A) $NaYbF_4$: 10%Gd, 8% Er, (B) $NaYbF_4$: 10% Gd, 8% Er@$NaYbF_4$: 8% Er, and (C) $NaYbF_4$: 10% Gd, 8% Er@$NaYbF_4$: 8% Er@ $NaYF_4$. (D) Size distribution of $NaYbF_4$: 10% Gd, 8% Er@$NaYbF_4$: 8% Er@ $NaYF_4$. (E) TEM images of silica coated $NaYbF_4$: 10% Gd, 8% Er@$NaYbF_4$: 8% Er@ $NaYF_4$. The silica shell thickness is ~8 nm.

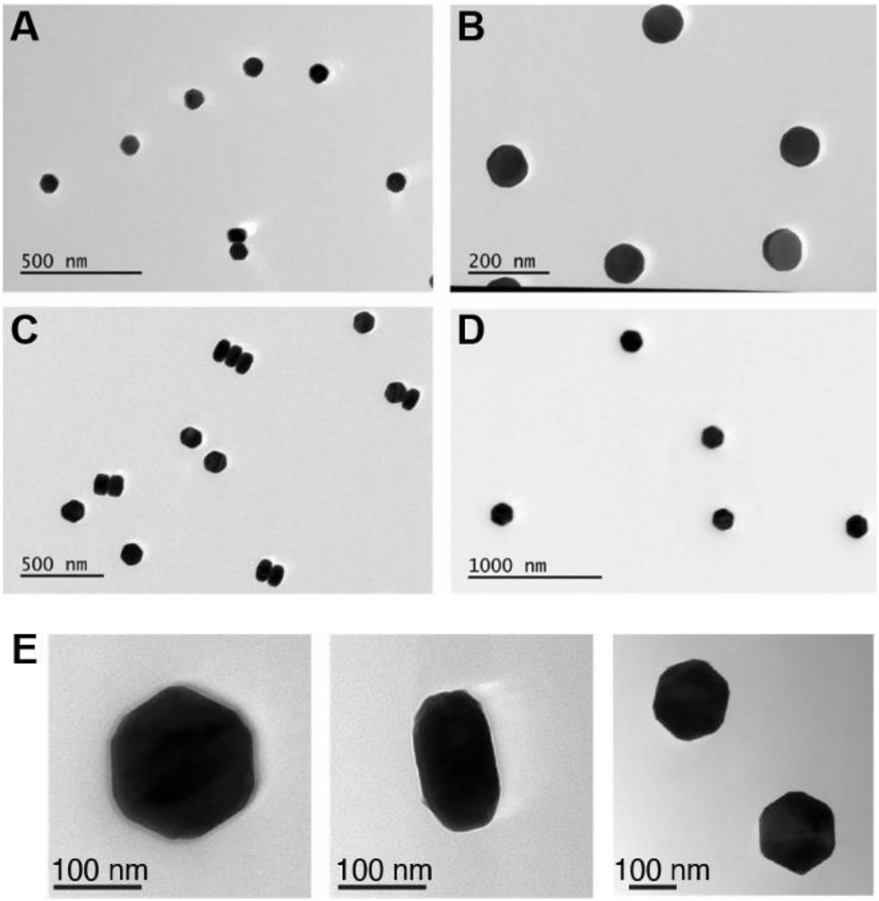


**Fig. S17.** TEM images of (A) $NaYbF_4$: 10% Gd, 8% Er (diameter of ~80 nm and height of ~60 nm), (B) $NaYbF_4$: 10% Gd, 8% Er@$NaYbF_4$: 8% Er (diameter of ~100 nm and height of ~70 nm), (C) $NaYbF_4$: 10% Gd, 8% Er@$NaYbF_4$: 8% Er@$NaYbF_4$: 8% Er (diameter of ~130 nm and height of ~80 nm), and (D) $NaYbF_4$: 10% Gd, 8% Er@$NaYbF_4$: 8% Er@$NaYbF_4$: 8% Er@$NaYF_4$ (diameter of ~160 nm and height of ~90 nm). (E) TEM images of silica coated $NaYbF_4$: 10% Gd, 8% Er@$NaYbF_4$: 8% Er@$NaYbF_4$: 8% Er@$NaYF_4$. The silica shell thickness is ~4 nm.

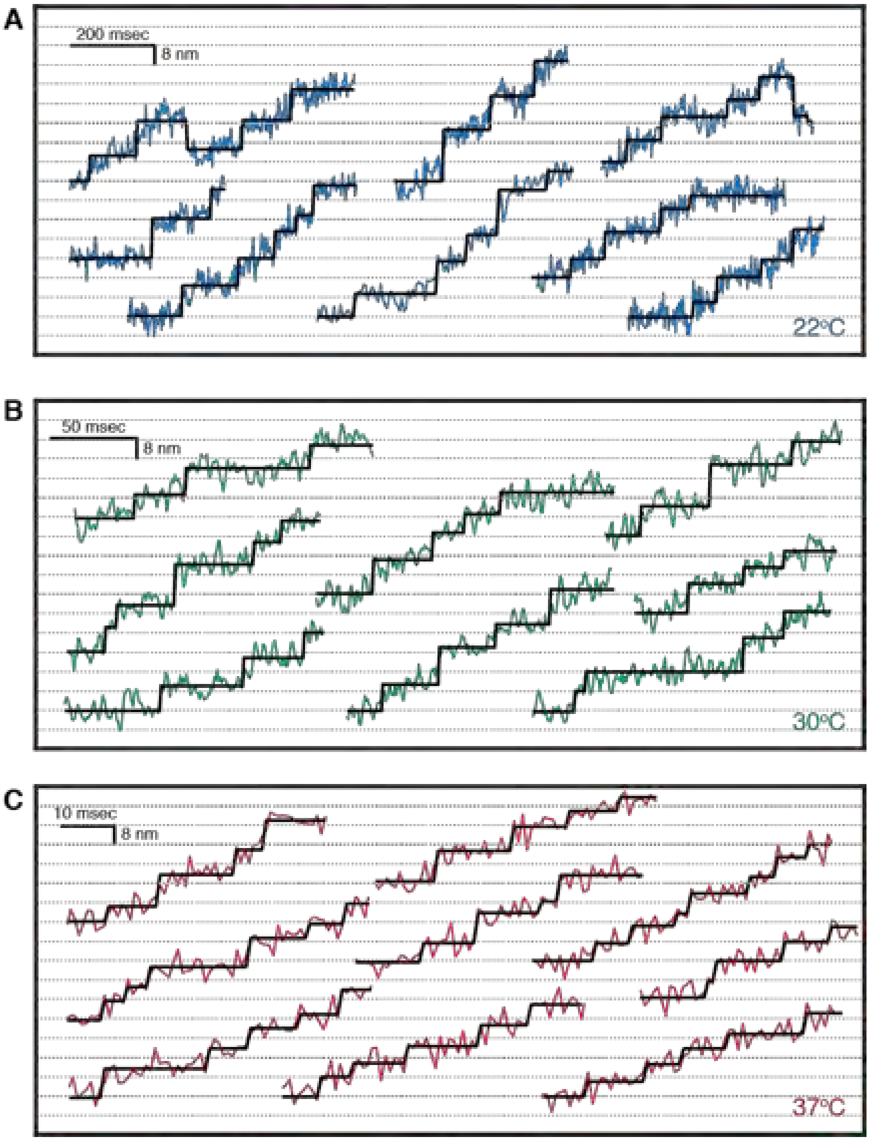


**Fig. S18.** More retrograde transport step-size traces as shown in Fig. 4. Displacement traces are broken into segments and displaced vertically for better visualization. The blue, green, and red lines are the raw trajectories at (A) 22 $°C$, (B) 30 $°C$, and (C) 37 $°C$, respectively. The black lines mark the steps detected by step-finding algorithm. Dynein steps exhibit 8 nm and larger steps. Notice that the different time scales are used at different temperatures.

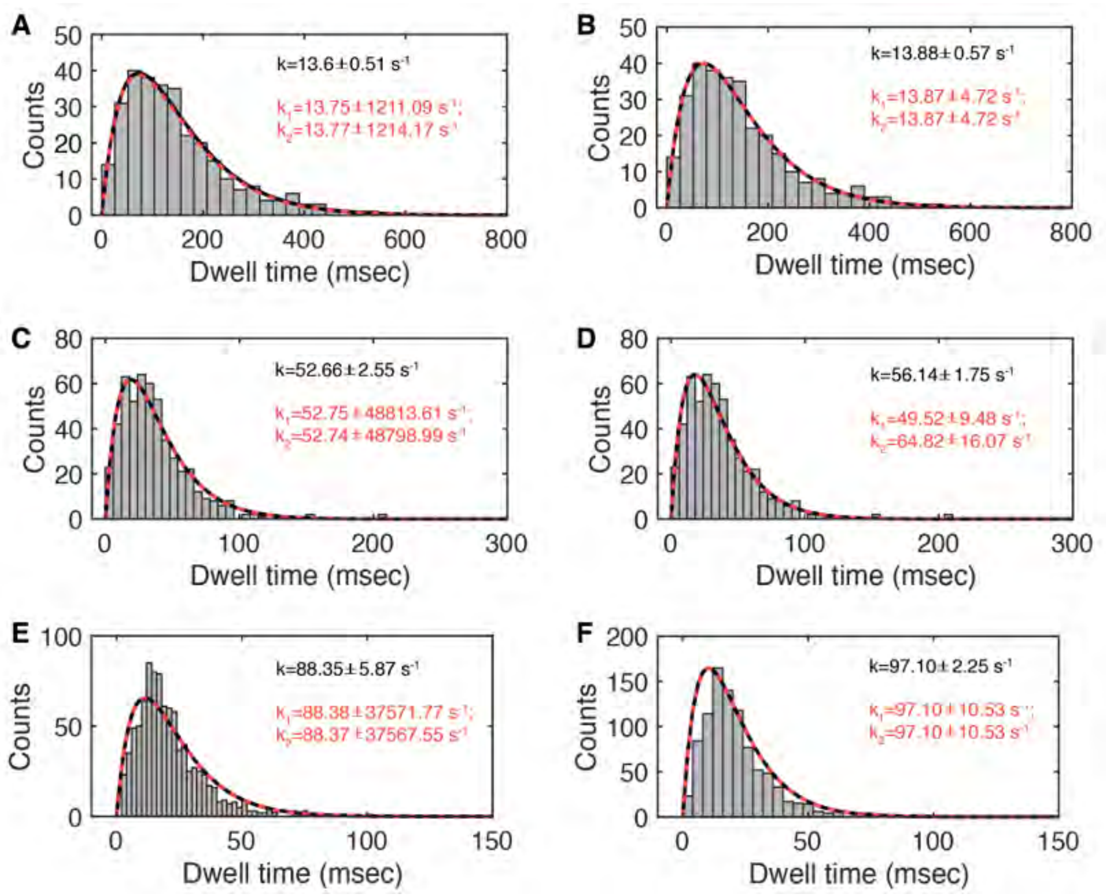


**Fig. S19.** Dwell-time histograms for retrograde transport in live rat DRG neurons at 22 $°C$ (A, B), 30 $°C$ (C, D), and 37 $°C$ (E, F). These histograms were fit to a convolution of two exponential functions, either with the same rate $\tau k^2 e^{-k\tau}$ (black curves), or with different rates $\frac{k_1 k_2}{k_1 - k_2}\left(e^{-k_2\tau} - e^{-k_1\tau}\right)$ (orange dash lines). Fitting was done using two different methods: (1) fit to binned histogram using *lsqnonlin* function in Matlab (A, C, E); and (2) directly fit the raw dwell times using Maximum Likelihood Estimates (MLE) (B, D, F). We note that the fitting results using (1) can vary with the choice of bin size, whereas the fitting results using (2) does not depend on the bin size because it directly fits to the raw dwell time data points. The error bars reported are 95% confidence interval for (1) and standard error for (2), which is the square root of the covariance matrix from MLE. In all cases except for (D), the fitting results indicated a model of the same rates. Even in (D), the single rate constant ($k = 56.14 \pm 1.75\ s^{-1}$) is within the error bars of the two different rates ($k_1 = 49.52 \pm 9.48\ s^{-1}; k_2 = 64.82 \pm 16.07\ s^{-1}$). Therefore, we constrained the model to having the same rate, $P(\tau) = \tau k^2 e^{-k\tau}$, which resulted in smaller error bars.

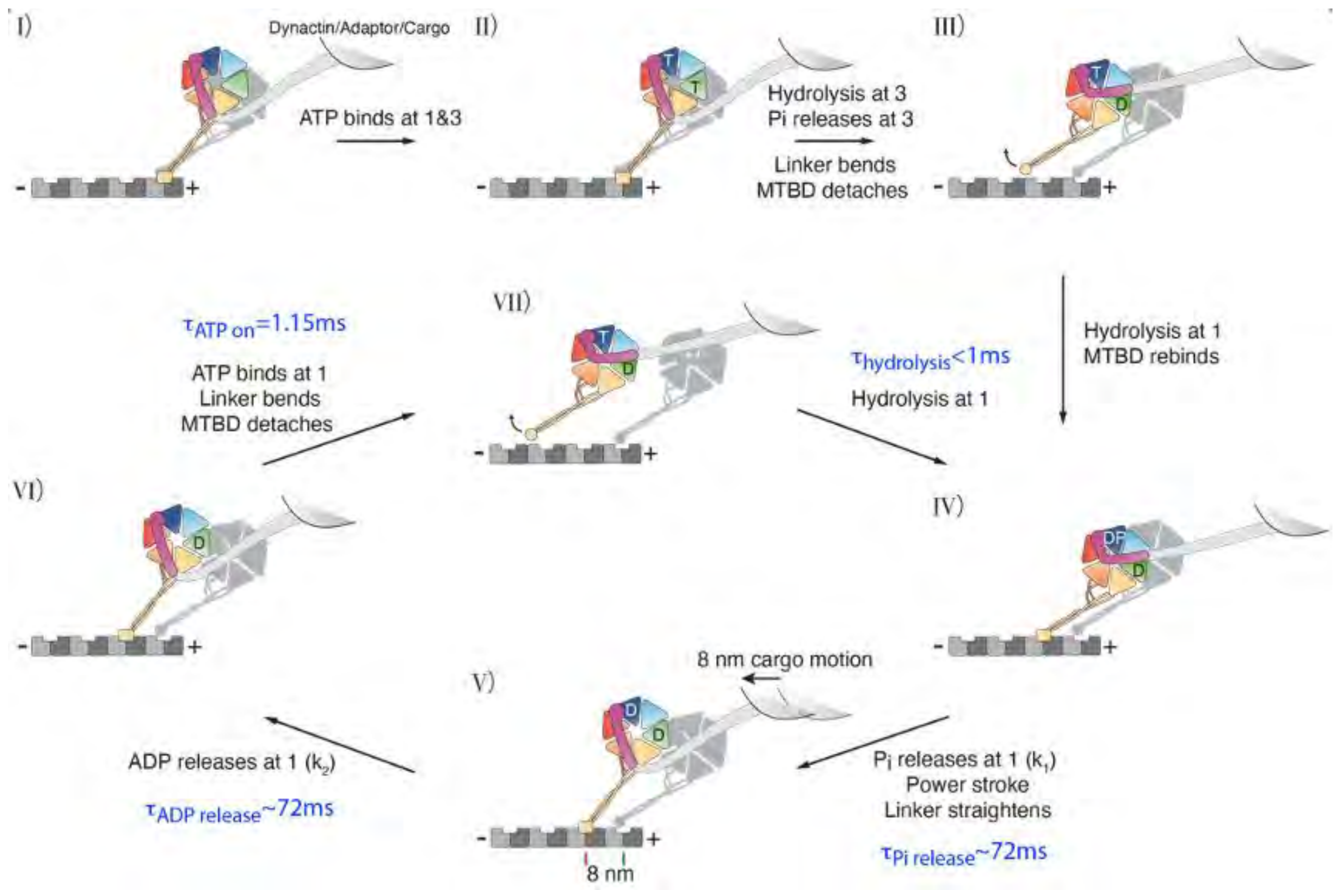


**Fig. S20**. Schematic of active cycling-model where only ATP hydrolysis at AAA1 is required for dynein stepping.

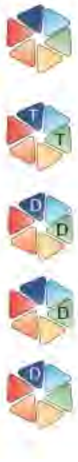

| Conditions | Mean backward force [pN] | 95% confidence interval [pN] | MT affinity |
|---|---|---|---|
| WT, apo | 3.3 | [3.1, 3.6] | S. B. |
| AAA1 E/Q + AAA3 E/Q, 1 mM ATP | 0.9 | [0.8, 0.9] | W. B. |
| WT, 2 mM ADP | 2.4 | [2.3, 2.5] | W. B. |
| AAA1 K/A, 2 mM ADP | 1.8 | [1.6, 2.0] | W. B. |
| AAA3 K/A, 2 mM ADP | 3.8 | [3.5, 4.1] | S. B. |

**Fig. S21**. Single-molecule unbinding force of the dynein motor in different nucleotide states reported in Ref (*25*). S.B. and W.B. microtubule (MT) affinity stands for strongly-bound and weakly-bound states, respectively.

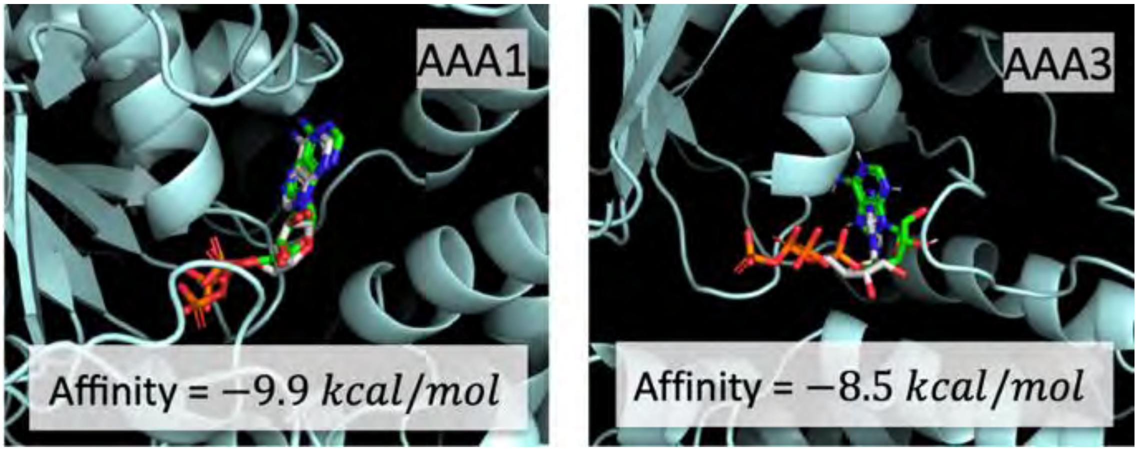


**Fig. S22**. Molecular docking of ADP at AAA1 and AAA3 sites using the ADP-dynein structure (PDB: 5NUG). The active cycling model requires that $ADP$ remains of the AAA3 site during many rounds of $ATP$ cycling. This suggest that $ADP$ is more tightly bound at AAA3 compared to AAA1. To examine the affinity of $ADP$ at AAA3, we performed molecular docking of $ADP$ at both the AAA1 and AAA3 sites. It was found that the binding energy of $ADP$ at the AAA1 and AAA3 sites were 9.9 kcal/mol and 8.5 kcal/mol, respectively. Since the off-rate $k_{\text{off}} \propto e^{-\Delta E_b/k_B T}$, $ADP$ is estimated to have a ~ 10-fold higher probability of release from AAA3 compared to AAA1.

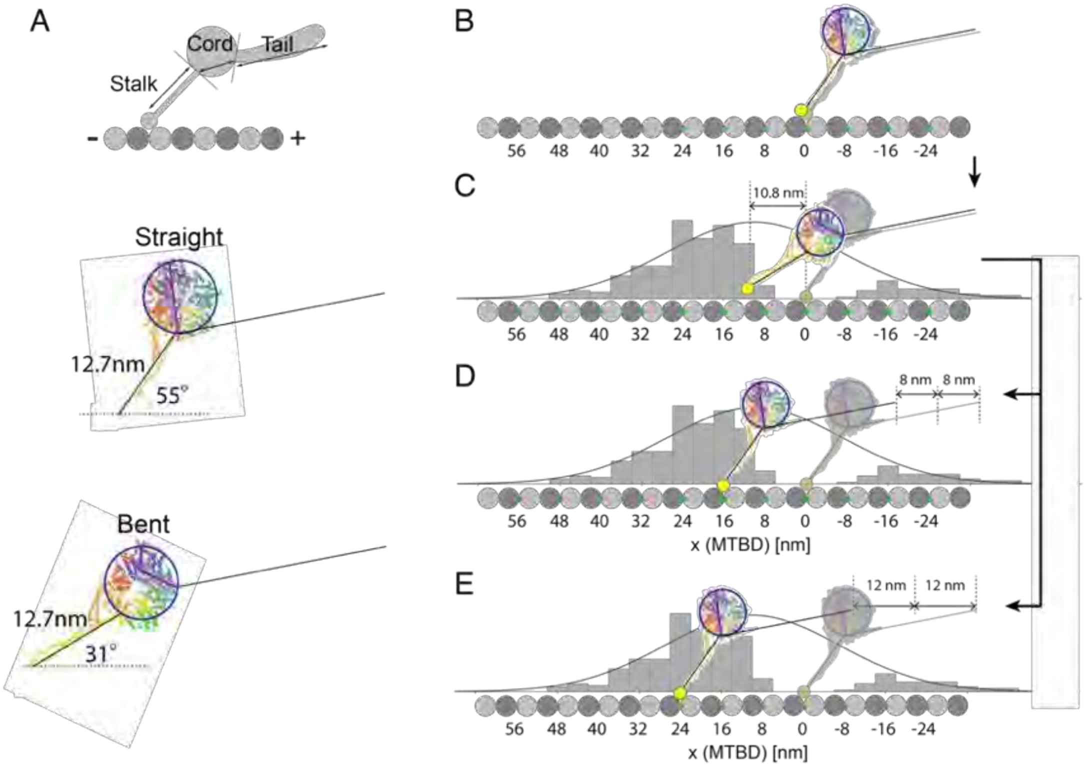


**Fig. S23.** Stepping model for dynein in the context of molecular structures. (A) Geometry of dynein in the "straight-linker" and "bent-linker" configurations based on the crytstal structures obtained in the $ADP$ (PDB: 3VKH) and $ADP \cdot Pi$ (PDB: 4R7H) states, respectively. The angles between the stalk and the MT are obtained from the negatively stained (*26*) and cryo-EM (*27*) structures. Note that in the proposed dynein cycle (Main Text, Fig. 5), the average cycle time is $144\ ms$. Each monomer spends $142\ ms$ in the flexible, pre-power stroke "bent linker" state. (B) The initial state assumes that a dynein monomer just completed a power stroke. The companion dynein monomer is in its flexible state, and bends to follow the motion. Both monomers are at $x = 0\ nm$. (C) Within $1\ ms$, the stepping monomer enters the weakly-bound, flexible state with the MTBD positioned $10.8\ nm$ forward based on the geometry shown in (A). The step-size histogram for the MTBD was approximated using the measured step-size histogram at $22°C$, but with twice the width. The $+10.8\ nm$ position matches the center of the Gaussian fit. Once the dynein monomer is in the WB state, it can diffuse and land at any of the allowed MT binding sites (denoted by the green circles) within the diffusion range. We show two examples (D) and (E) here. (D) The MTBD lands $+16\ nm$ forward and transitions to the strongly-bound state. The power stroke moves the stepping dynein tail forward by $+16\ nm$, but then quickly relaxes into the flexible state. This transient motion occurs too rapidly to be detected and the net observed motion is $+8\ nm$, half-way between the stepping and the non-stepping monomers. (E) In another scenario, the leading monomer undergoes a power stroke at a position $+24\ nm$ forward of its initial position.

Single-molecule force experiments have shown that dynein has asymmetric response to tension with a much weaker binding to the MT when pulled towards the MT minus end (*13, 25*). Even with less than $2\ pN$ forward tension, the trailing monomer can be released from the MT with a release rate of $20\ s^{-1}$ (measured for AAA1 E/Q mutant monomers in the presence of ATP) (*13*). It appears likely that the detachment force asymmetry allows the trailing monomer to detach from the MT briefly to relieve the mutual stress if the two MTBD positions become too separated.

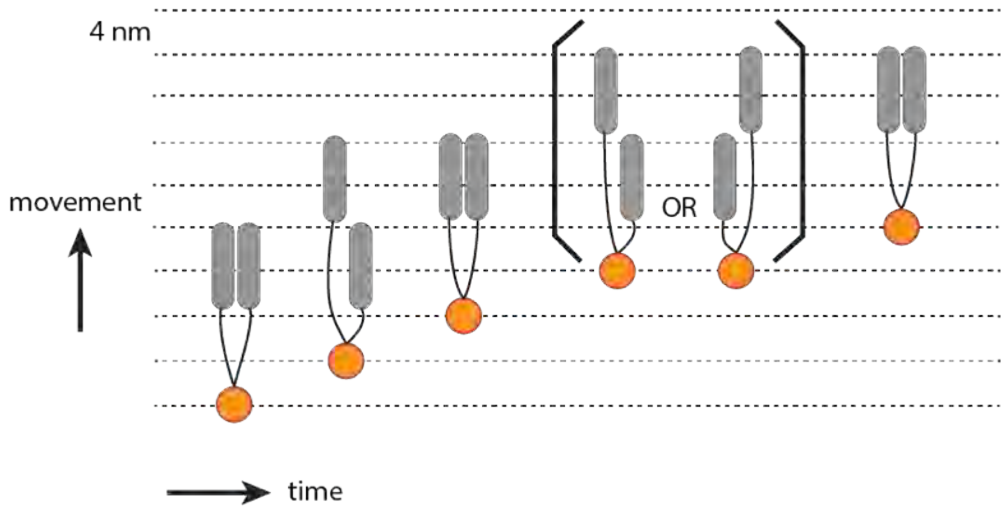


**Fig. S24**. Top view of the proposed dynein stepping model. In Fig. S23, we argued that the measured cargo position (orange circle) is the compromised position of the two dynein monomers. *In vitro* optical tweezers measurements showed asymmetric unbinding force of dynein from the microtubule (MT) (*13, 25*). After a forward step, the lagging dynein monomer will be subject to a forward force that could be large enough to cause it to detach and reattach as shown in the second- and third-time steps. Once the lagging monomer catches up, either one of the two monomers is allowed to step forward. It is still possible for the leading monomer to take consecutive steps before the lagging monomer catches up (*10, 12*), but as the distance between the leading MTBDs increases, those configurations become less likely. Note that we show the monomer taking 8-nm steps for simplicity, but it is allowed to take larger steps as shown in Fig. S23.

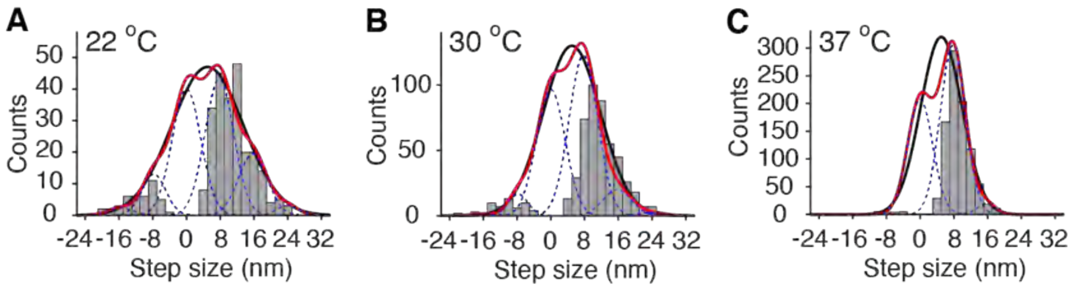


D

| 22°C | | | | | | | |
|---|---|---|---|---|---|---|---|
| | Single | Multiple Gaussians | | | | | |
| $x_0$ [nm] | 5 | -15.8 | -7.8 | -0.2 | 7.8 | 15.8 | 24 |
| $\sigma$ [nm] | 8.5 | 3 | 2.8 | 3.4 | 3.4 | 3.4 | 2.5 |
| Amp | 47 | 2 | 13 | 40 | 43 | 19 | 3 |
| **30°C** | | | | | | | |
| | Single | Multiple Gaussians | | | | | |
| $x_0$ [nm] | 5 | -15.8 | -7.8 | 0 | 7.8 | 15.8 | 23.8 |
| $\sigma$ [nm] | 6.8 | 3.1 | 2.3 | 3.4 | 3.4 | 3.4 | 3.4 |
| Amp | 130 | 2 | 17 | 97 | 123 | 20 | 2 |
| **37°C** | | | | | | | |
| | Single | Multiple Gaussians | | | | | |
| $x_0$ [nm] | 5 | | -8.3 | 0 | 7.8 | 15.8 | |
| $\sigma$ [nm] | 4.9 | | 2.1 | 3.0 | 3.0 | 2.1 | |
| Amp | 300 | | 0.1 | 208 | 305 | 20 | |

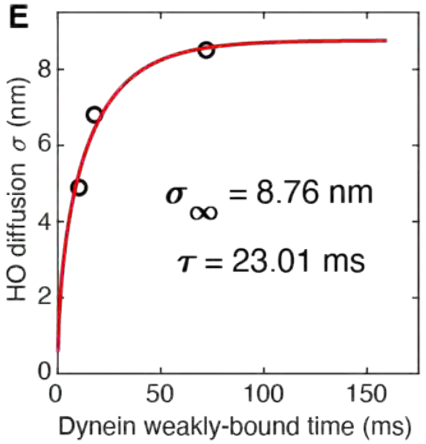


**Fig. S25**. Dynein step-size histograms measured at (A) 22 °$C$, (B) 30 °$C$, and (C) 37 °$C$, respectively. The solid black line is a single Gaussian fit to the overall distribution. The center ($x_0 = 5\ nm$) of the black line Gaussian fits is temperature independent. The dotted blue lines represent multiple Gaussian fits to the discrete peaks, suggesting that the vesicle motion is quantized in partially resolved 8 $nm$ steps. However, we believe the true quantization are 4 $nm$ steps, which would be observable it the spatial resolution were 0.1 $nm$ as explained to Fig. S26. (D) Table of all the fit parameters. (E) The constrained diffusion fit (red line) to the temperature-dependent widths of the step-size histograms and dynein spent dynein spends in the weakly bound state.

At the molecular scale, inertial effects are negligible, and the motor and cargo system experience a viscous drag force $F_d = \gamma \dot{x}(t)$, where $\gamma = 6\pi\eta a$ for a sphere of radius $a$ immersed in medium with viscosity $\eta$. It can be shown that the average diffusive motion is $\langle x^2(t)\rangle \equiv \sigma^2(t, T_{avg}) = \frac{2k_B T_{avg}}{\kappa}\left(1 - e^{-t\kappa/\gamma}\right)$ (*28*), where the time $t$ is the time spent in State II and $T_{avg} = 303\ K$ is the average temperature of 22°C, 30°C and 37°C in degrees Kevin. In (E), the three measurements are fit to $\langle x^2(\infty)\rangle \equiv \sigma_\infty = (8.76\ nm)^2$ and $\tau = \gamma/\kappa = 23\ ms$. For $t \ll \gamma/\kappa$, the free Brownian diffusion is described by the Einstein equation $\langle x^2(t)\rangle = 2Dt$. For long times, $\langle x^2\rangle_{t\to\infty} = 2k_B T_{avg}/\kappa$, and $\kappa = 1.1\ \times 10^{-4} J \cdot m^{-2}$.

The time $t$ in Fig. S25E is the time the dynein is in the weakly-bound state (State II). While in the weakly-bound state, the dynein is allowed to explore the position of its next step only when its MTBD is detached from the MT. In Note S2, we show that the fitted parameters $\tau = \gamma/\kappa$ and $\sigma_\infty$ are in agreement with the physical size of a dynein monomer, reasonable estimates of viscosity of an object the size of the dynein in axonal cytosol, and the kinetic binding and unbinding rates of dynein in the pre-power stroke state (*17*).

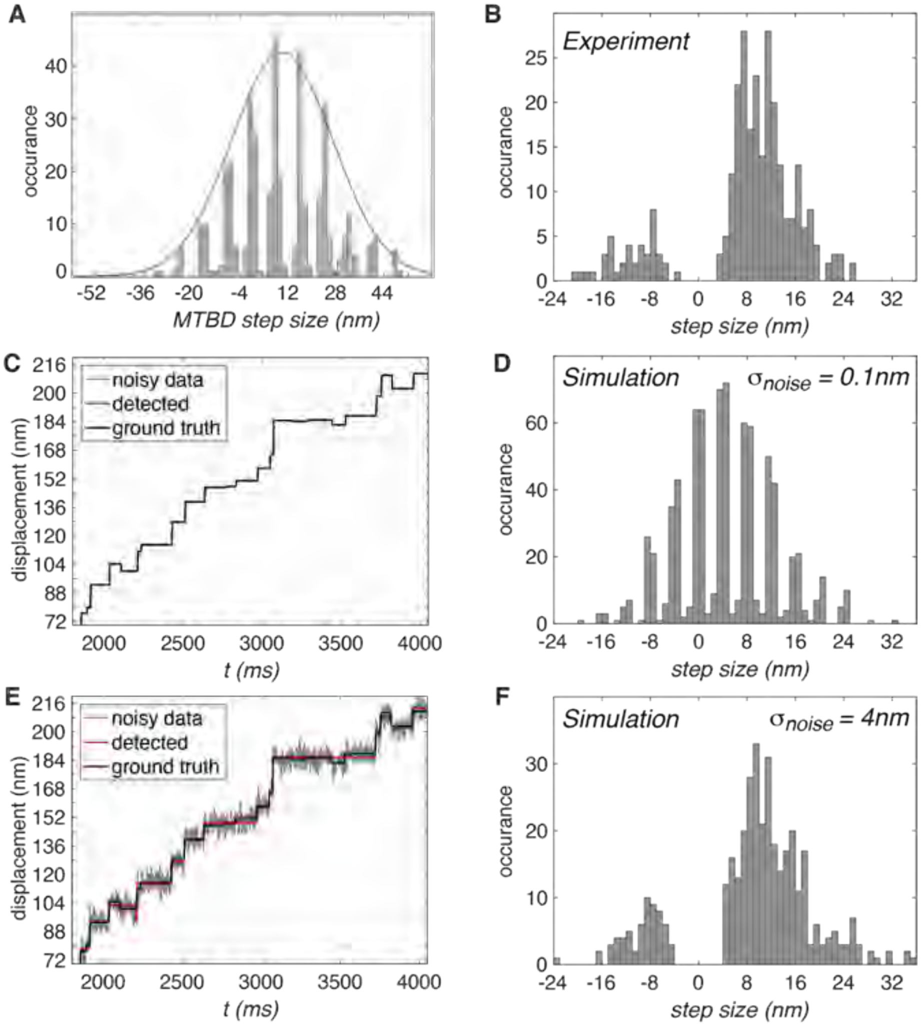


**Fig. S26.** (A) The simulation input of the MTBD step-size histogram, which was generated based on the experimentally measured cargo step-size histogram at $22°C$ shown in panel (B). The width of the probability distribution was taken to be twice of that of the cargo distribution. Multiple Gaussian distributions with a $\sigma = 1\ nm$ were placed at every $8\ nm$ to represent the allowed binding sites on the MT. The center of the overall distribution (red curve) is at $10\ nm$. (C) Simulated trajectory for a system with two independent dynein monomers. Noise of $\sigma_{noise} = 0.1\ nm$ was added. (D) The detected step-size historgram from the trajectory in (C). (E) Simulated trajectory with a noise of $\sigma_{noise} = 4\ nm$. (F) The detected step-size historgram from the trajectory in (E).

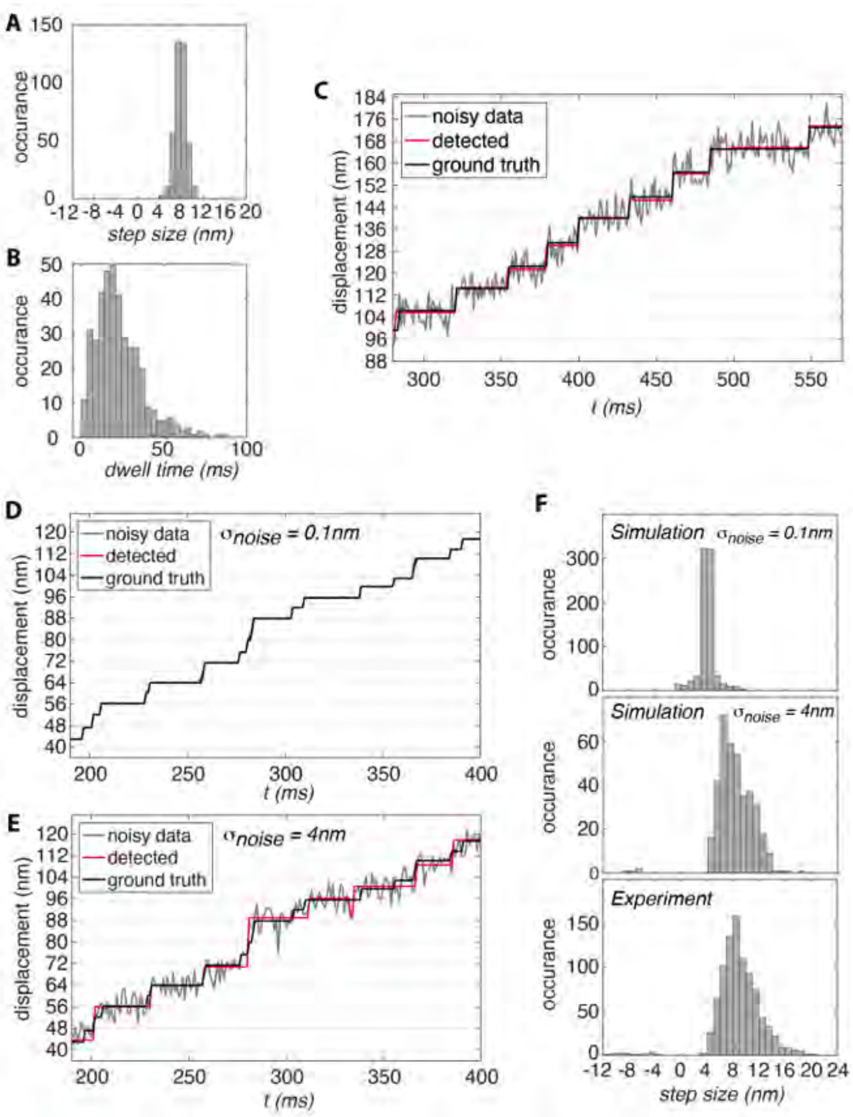


**Fig. S27.** (A) Simulation input step-size histogram centered at 8 $nm$. (B) Simulation input dwell-time histogram based on the experimentally measured rates at $37°C$ and $[ATP] = 3\ mM$. (C) Simulated traces for a *single* motor using the step-size histogram (A) and dwell-time histogram (B). The solid black line is the simulated ground truth trace. The gray line is the simulation with 4 $nm$ of added experimental noise. The red line shows the detected stepping trace using the step-finding algorithm, which accurately determined the occurrences of individual steps. (D) The simulated stepping trace of a system with *two* motors. The two motors were simulated to step independently, and the cargo position was taken as the average of the two motor positions. Experimental noise of 0.1 $nm$ was added. (E) Simulation of the two independent motor, but with experimental noise of 4 $nm$. Note that most of the 4 $nm$ steps were missed by the step-finding algorithm. (F) Comparison of the experimentally measured and the detected step-size histograms for the simulations with $\sigma_{noise} = 0.1\ nm\ or\ 4\ nm$. Note that the major peak is at 4 $nm$ with $\sigma_{noise} = 0.1\ nm$ due to the two motors splitting the 8 $nm$ steps. The detected step-size histogram for $\sigma_{noise} = 4\ nm$ shows a major peak at 8 $nm$, which is similar to our experimental step-size histogram at $37°C$.

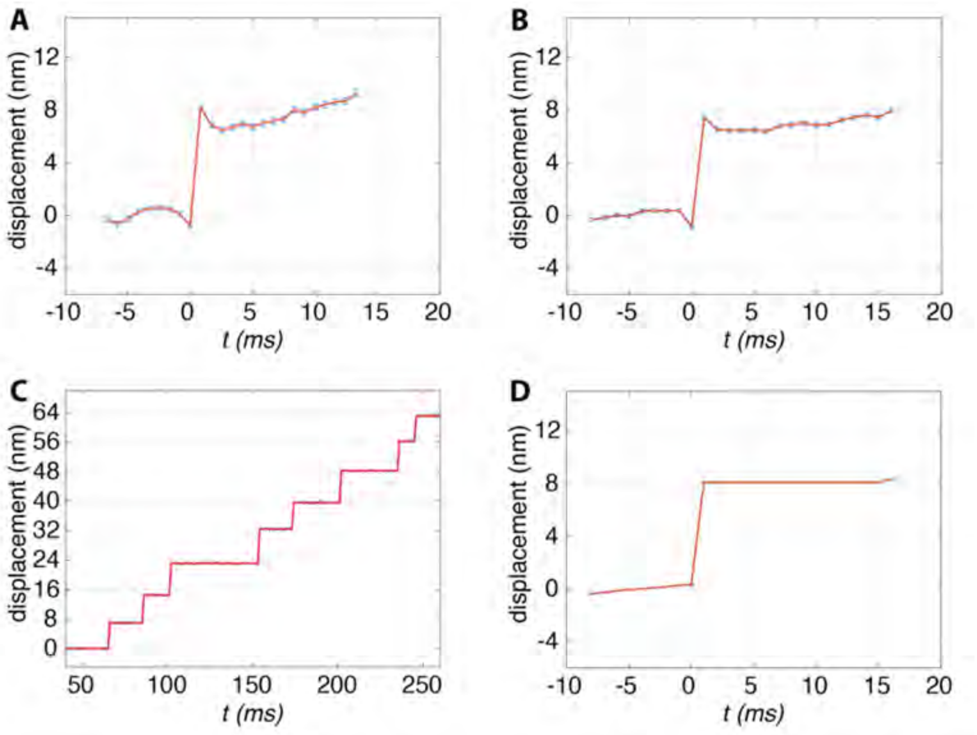


**Fig. S28.** (A) Concatenated and averaged stepping transition of the experimental trace at $37°C$. The errors in the mean are shown as the error bars. (B) Concatenated and averaged stepping transition of the simulated trace in Fig. S25E with $\sigma_{noise} = 4\ nm$. The slight dip and peak in the time bins immediately before and after the step transition, is a good approximation to the actual averaged data shown in (A). These features are mostly artifacts of the step-finding algorithm when applied to data with large noise since the algorithm would tend to favor bigger transitions. (C) Simulated trace of a single motor with $\sigma_{noise} = 0.1\ nm$, showing that the step-finding algorithm can perfectly identify all the transitions. (D) The concatenated and averaged stepping transition of the trace shown in (C) does not result in the artifact features around the transition.

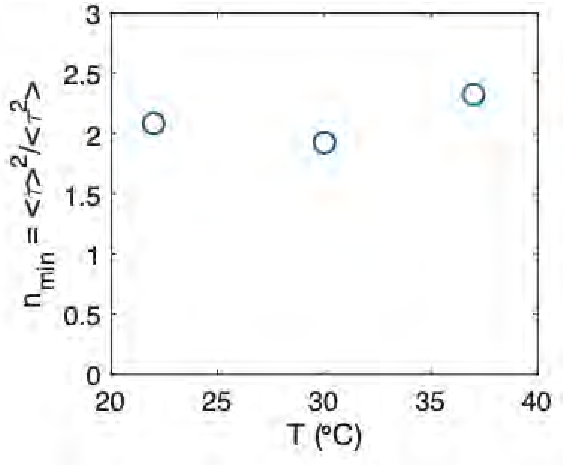


**Fig. S29**. Experimentally measured $n_{min}$ values from the dwell time distributions at $T = 22°C, 30°C$, and $37°C$.

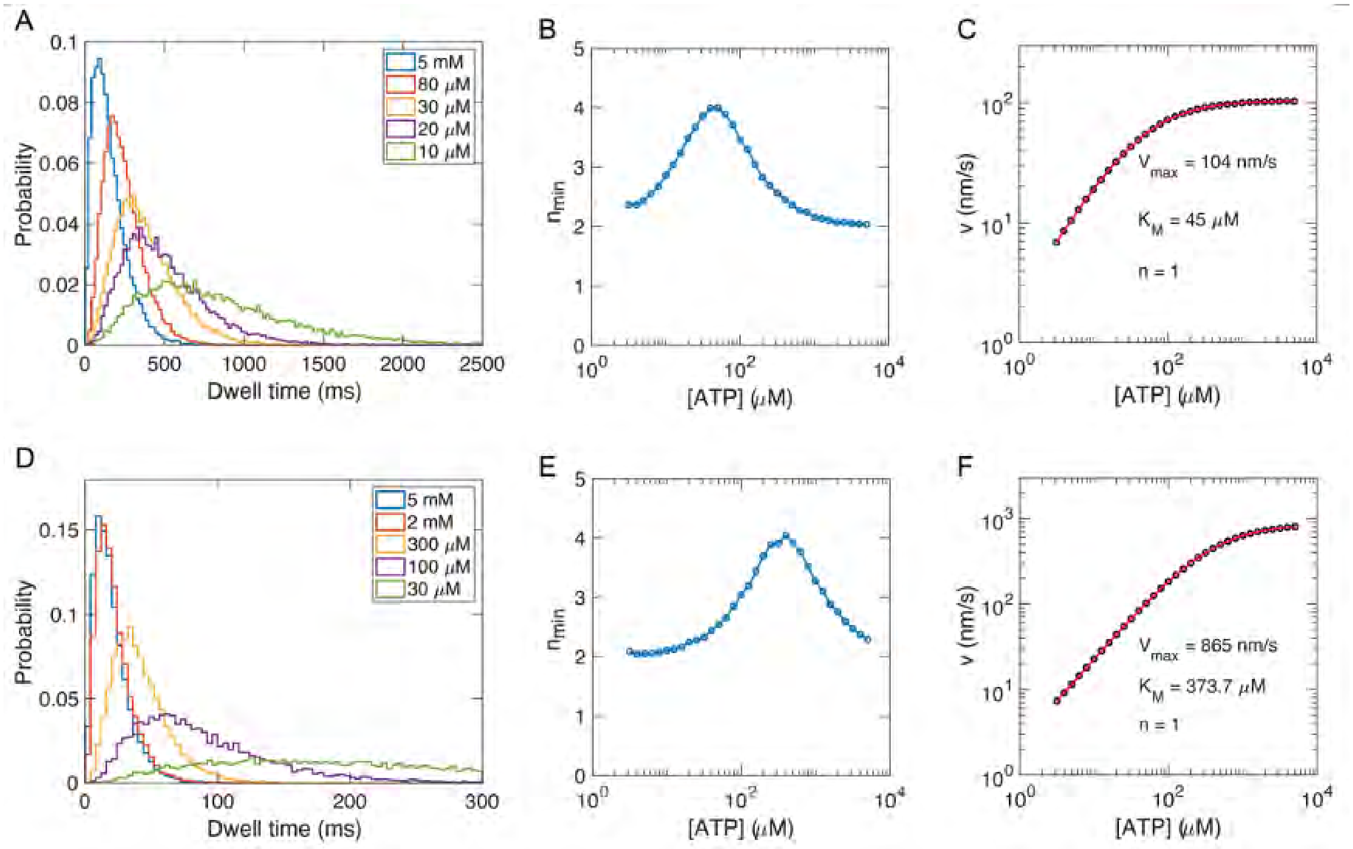


**Fig. S30**. Simulated dwell time distributions (A, D), randomness factors (B, E), and velocities (C, F) at various ATP concentrations. The simulations were performed for $T = 22°C$ (A-C) and $T = 37°C$ (D-F). In the limit of low or high $[ATP]$ ($< 5\ \mu M$ or $> 1\ mM$) $n_{min} \rightarrow 2$. In order to clearly see the fast rise in the dwell-time distribution, sufficient time resolution and fine time binning of the histogram are required. As discussed in Fig. S19, the parameters of a dwell-time distribution function can be determined independent of the choice of the histogram bin size by fitting the raw dwell times using Maximum Likelihood Estimates (MLE).

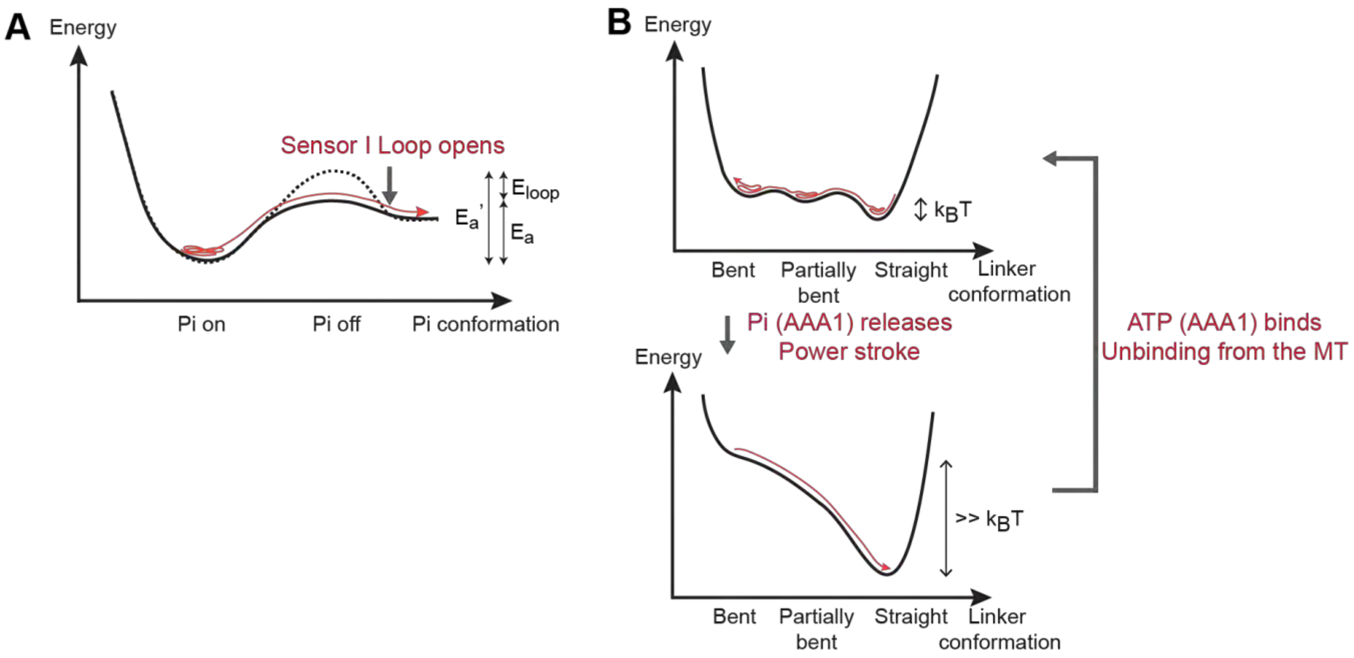


**Fig. S31**. (A) Potential energy surfaces for the $Pi$ configurations before and after the Sensor I Loop opens. While Sensor I Loop has contacts with the $Pi$, the energy barrier prevents thermal desorption. The opening of the Sensor I Loop lowers the energy barrier for $Pi$ to release, which is a thermally activated process. The energy surfaces apply to both the AAA1 and AAA3 sites. (B) Potential energy surfaces for the dynein linker conformations before and after the power stroke. The linker conformations are based on the cryo-EM structures of the dynein in the $ATP \cdot Pi$ state (*29*).

### Supplementary Tables

**Table S1.**

Preparation of silica-coated UCNPs.

| Reagents | UCNP-1 | UCNP-2 | UCNP-3 | UCNP-4 |
|---|---|---|---|---|
| Igepal CO-520 (mg) | 1000 | 500 | 1500 | 1500 |
| UCNPs (μL) | 200 | 100 | 333 | 100 |
| Ammonia solution (μL) | 150 | 150 | 70 | 70 |
| TEOS (μL) | 12 | 12 | 100 | 5 |

UCNP-1: 22 nm $NaYF_4$: 20% Yb, 2 % Er
UCNP-2: 29.4 nm $NaYF_4$@$NaYbF_4$: 8 % Er@$NaYF_4$
UCNP-3: 66.4 nm $NaYbF_4$: 8 % Er, 10% Gd@$NaYbF_4$: 8 % Er@$NaYF_4$
UCNP-4: 160 nm $NaYbF_4$: 10% Gd, 8% Er@$NaYbF_4$: 8% Er@$NaYbF_4$: 8% Er@$NaYF_4$

**Table S2.**

Predicted freely-diffusing time at $22\ °C$, $30\ °C$ and $37\ °C$.

| Temperature | $\sigma_{cargo}$ | Unbound time |
|---|---|---|
| $22°C$ | 8.5 nm | $450\ \mu s$ |
| $30°C$ | 6.8 nm | $250\ \mu s$ |
| $37°C$ | 4.9 nm | $116\ \mu s$ |

### References


1. Q. Liu *et al.*, Single upconversion nanoparticle imaging at sub-10 W cm2 irradiance. *Nature Photonics* **12**, 548-553 (2018).
2. R. Abdul Jalil, Y. Zhang, Biocompatibility of silica coated NaYF(4) upconversion fluorescent nanocrystals. *Biomaterials* **29**, 4122-4128 (2008).
3. Q. Liu, W. Feng, T. Yang, T. Yi, F. Li, Upconversion luminescence imaging of cells and small animals. *Nature protocols* **8**, 2033-2044 (2013).
4. M. Kyoung, Y. Zhang, J. Diao, S. Chu, A. T. Brunger, Studying calcium-triggered vesicle fusion in a single vesicle-vesicle content and lipid-mixing system. *Nature protocols* **8**, 1-16 (2013).
5. Y. A. Huang, B. Zhou, M. Wernig, T. C. Sudhof, ApoE2, ApoE3, and ApoE4 Differentially Stimulate APP Transcription and Abeta Secretion. *Cell* **168**, 427-441.e421 (2017).
6. A. Serge, N. Bertaux, H. Rigneault, D. Marguet, Dynamic multiple-target tracing to probe spatiotemporal cartography of cell membranes. *Nat Methods* **5**, 687-694 (2008).
7. J. Chen *et al.*, Single-molecule dynamics of enhanceosome assembly in embryonic stem cells. *Cell* **156**, 1274-1285 (2014).
8. N. Monnier *et al.*, Inferring transient particle transport dynamics in live cells. *Nat Methods* **12**, 838-840 (2015).
9. J. W. Kerssemakers *et al.*, Assembly dynamics of microtubules at molecular resolution. *Nature* **442**, 709-712 (2006).
10. S. L. Reck-Peterson *et al.*, Single-molecule analysis of dynein processivity and stepping behavior. *Cell* **126**, 335-348 (2006).
11. A. P. Carter *et al.*, Structure and Functional Role of Dynein's Microtubule-Binding Domain. *Science* **322**, 1691-1695 (2008).
12. M. A. DeWitt, A. Y. Chang, P. A. Combs, A. Yildiz, Cytoplasmic dynein moves through uncoordinated stepping of the AAA+ ring domains. *Science* **335**, 221-225 (2012).
13. M. A. DeWitt, C. A. Cypranowska, F. B. Cleary, V. Belyy, A. Yildiz, The AAA3 domain of cytoplasmic dynein acts as a switch to facilitate microtubule release. *Nature structural & molecular biology* **22**, 73-80 (2015).
14. J. R. Moffitt *et al.*, Intersubunit coordination in a homomeric ring ATPase. *Nature* **457**, 446-450 (2009).
15. J. L. Ross, K. Wallace, H. Shuman, Y. E. Goldman, E. L. F. Holzbaur, Processive bidirectional motion of dynein–dynactin complexes in vitro. *Nature Cell Biology* **8**, 562-570 (2006).
16. Y. Kinoshita, T. Kambara, K. Nishikawa, M. Kaya, H. Higuchi, Step Sizes and Rate Constants of Single-headed Cytoplasmic Dynein Measured with Optical Tweezers. *Scientific reports* **8**, 16333 (2018).
17. Y. Kinoshita, T. Kambara, K. Nishikawa, M. Kaya, H. Higuchi, Step Sizes and Rate Constants of Single-headed Cytoplasmic Dynein Measured with Optical Tweezers. *Scientific Reports* **8**, (2018).
18. J. Ando *et al.*, Small stepping motion of processive dynein revealed by load-free high-speed single-particle tracking. *Scientific reports* **10**, 1-11 (2020).
19. W. B. Redwine *et al.*, Structural Basis for Microtubule Binding and Release by Dynein. *Science* **337**, 1532-1536 (2012).
20. D. Wirtz, Particle-tracking microrheology of living cells: principles and applications. *Annu Rev Biophys* **38**, 301-326 (2009).

21. J. T. Mika, P. E. Schavemaker, V. Krasnikov, B. Poolman, Impact of osmotic stress on protein diffusion in Lactococcus lactis. *Mol Microbiol* **94**, 857-870 (2014).
22. A. S. Verkman, Solute and macromolecule diffusion in cellular aqueous compartments. *Trends in Biochemical Sciences* **27**, 27 - 33 (2002).
23. K. Kwapiszewska *et al.*, Nanoscale Viscosity of Cytoplasm Is Conserved in Human Cell Lines. *The journal of physical chemistry letters* **11**, 6914-6920 (2020).
24. J. C. Wortman *et al.*, Axonal transport: how high microtubule density can compensate for boundary effects in small-caliber axons. *Biophysical journal* **106**, 813-823 (2014).
25. M. P. Nicholas *et al.*, Cytoplasmic dynein regulates its attachment to microtubules via nucleotide state-switched mechanosensing at multiple AAA domains. *Proc Natl Acad Sci U S A* **112**, 6371-6376 (2015).
26. S. A. Burgess, M. L. Walker, H. Sakakibara, P. J. Knight, K. Oiwa, Dynein structure and power stroke. *Nature* **421**, 715-718 (2003).
27. S. Can, S. Lacey, M. Gur, A. P. Carter, A. Yildiz, Directionality of dynein is controlled by the angle and length of its stalk. *Nature* **566**, 407-410 (2019).
28. S. F. Nørrelykke, H. Flyvbjerg, Harmonic oscillator in heat bath: Exact simulation of time-lapse-recorded data and exact analytical benchmark statistics. *Physical Review E* **83**, (2011).
29. G. Bhabha *et al.*, Allosteric communication in the dynein motor domain. *Cell* **159**, 857-868 (2014).
